# T-cell receptor specificity landscape revealed through de novo peptide design

**DOI:** 10.1101/2025.02.28.640903

**Authors:** Gian Marco Visani, Michael N. Pun, Anastasia A. Minervina, Philip Bradley, Paul Thomas, Armita Nourmohammad

**Affiliations:** Paul G. Allen School of Computer Science and Engineering, University of Washington, 85 E Stevens Way NE, Seattle, WA 98195, USA; Department of Physics, University of Washington, 3910 15th Avenue Northeast, Seattle, WA 98195, USA; St. Jude Children’s Research Hospital, Memphis, TN, USA; Fred Hutchinson Cancer Center, 1241 Eastlake Ave E, Seattle, WA 98102, USA; Department of Applied Mathematics, University of Washington, 4182 W Stevens Way NE, Seattle, WA 98105, USA

## Abstract

T-cells play a key role in adaptive immunity by mounting specific responses against diverse pathogens. An effective binding between T-cell receptors (TCRs) and pathogen-derived peptides presented on Major Histocompatibility Complexes (MHCs) mediate an immune response. However, predicting these interactions remains challenging due to limited functional data on T-cell reactivities. Here, we introduce a computational approach to predict TCR interactions with peptides presented on MHC class I alleles, and to design novel immunogenic peptides for specified TCR-MHC complexes. Our method leverages HERMES, a structure-based, physics-guided machine learning model trained on the protein universe to predict amino acid preferences based on local structural environments. Despite no direct training on TCR-pMHC data, the implicit physical reasoning in HERMES enables us to make accurate predictions of both TCR-pMHC binding affinities and T-cell activities across diverse viral epitopes and cancer neoantigens, achieving up to 0.72 correlation with experimental data. Leveraging our TCR recognition model, we develop a computational protocol for de novo design of immunogenic peptides. Through experimental validation in three TCR-MHC systems targeting viral and cancer peptides, we demonstrate that our designs—with up to five substitutions from the native sequence—activate T-cells at success rates of up to 50%. Lastly, we use our generative framework to quantify the diversity of the peptide recognition landscape for various TCR-MHC complexes, offering key insights into T-cell specificity in both humans and mice. Our approach provides a platform for immunogenic peptide and neoantigen design, as well as for evaluating TCR specificity, offering a computational framework to inform design of engineered T-cell therapies and vaccines.

## I. INTRODUCTION

T-cells play a key role in the adaptive immune system, contributing to immune surveillance and response to pathogens. T-cells recognize pathogen-derived epitopes in the form of short protein fragments (peptides) displayed on specific molecules known as Major Histocompatibility Complexes (MHCs). The binding between a T-cell receptor (TCR) and a peptide-MHC (pMHC) is essential to mediate a protective T-cell response to an infection. To defend the host against a multitude of pathogens, a highly diverse TCR repertoire is generated through a somatic VDJ recombination process, and selected to target infected cells with remarkable sensitivity and specificity.

Predicting the TCR-pMHC binding specificity is an important step in characterizing the efficacy of the immune system in countering different pathogens. Such understanding would aid with disease diagnoses and development of diagnostic tests for early detection of autoimmune diseases [1]. A predictive model for TCR-pMHC specificity can be used to design engineered TCRs that recognize cancer antigens [2], enhancing the effectiveness of adoptive cell transfer and Chimeric Antigen Receptor (CAR) T-cell therapies [3, 4], as well as the design of soluble TCRs and bispecific T-cell engager therapeutics [5]. Moreover, it can enable the design of optimized antigens for vaccines to elicit robust T-cell responses against specific pathogens or tumors [6–8].

One bottleneck in deciphering the TCR-pMHC code is the lack of well-curated datasets. High-throughput immune repertoire sequencing, combined with comparative analyses of hosts facing similar immune challenges, and the development of functional assays (e.g., pMHC-tetramer staining [9] or MIRA [10]), have enabled the identification of diverse TCR pools with potential reactivity to specific antigens [11], collected in repositories like Immune Epitope Database (IEDB) [12] and VDJdb [13]. However, the current databases are unbalanced, with only a handful of dominant peptides (e.g. derived from SARS-CoV-2, CMV, EBV, Influenza, etc.) and MHC alleles present, each linked to hundreds of TCRs.

As a result, statistical inference and machine learning methods trained on such data to predict TCR-pMHC reactivities lack generalizability beyond their training data [14–16]. On the structural side, only a few hundred TCR-pMHC crystal structures have been experi-mentally generated to date. This is in part because TCR-pMHC interactions are typically of low affinity and are difficult to stabilize to the level needed for CryoEM or X-ray crystallography [17]. Importantly, this scarcity hinders computational prediction methods like AlphaFold in modeling reliable TCR-pMHC structures [18, 19]. Machine learning models, such as large language models, pre-trained on vast unannotated protein datasets can generate meaningful data representations [20, 21]. Fine-tuning these models on smaller, labeled datasets for specific tasks (e.g., predicting mutation impacts on stability or function) has proven highly effective [22–25]. Pre-trained sequence-based protein language models have been employed to predict reactivities of antibodies to antigens [24, 25], or TCRs to pMHC complexes [16, 26]. However, lack of generalizability due to sparse and skewed training data has remained a challenge in these models.

Protein-protein interactions in general, and immune-pathogen interaction in particular, rely on the complementarity in the 3D structures and amino acid compositions of the interacting protein pairs. Even though the interacting amino acids can be far apart in a protein sequence, they are close to each other in structure, resulting in local interactions that can determine immune-pathogen recognition. Machine learning models trained on protein structures instead of sequences can learn local structural representations for amino acid statistics, which tend to be more generalizable across different protein families and beyond their training sets [27, 28, 31–34].

Here, we present a structure-based approach to predict interaction affinities between TCRs and peptides presented on MHC class I, and to design reactive peptides for specific TCR-MHC-I complexes (TCR-MHC for short). Due to the limited availability of structural data for TCRs, we use HERMES– a physics-guided, 3D-equivariant machine learning model trained on the entire protein universe to predict amino acid preferences based on their local structural environments [27, 28].

We demonstrate that HERMES accurately predicts both binding affinity of TCR-pMHC Class I complexes and the functional responses and T-cell activities induced by a range of peptides presented on MHC molecules. Affinity of TCR-pMHC complexes, measured as the dissociation constant *K*_*d*_ by surface plasmon resonance, is labor intensive to obtain. Functional readout used to measure T-cell activities, including measurements of T-cell proliferation, labeled cellular activation markers, and cytokine (e.g., IFN-*γ*) release, are easier to obtain than affinity measurements. Yet, functional readouts are shaped not only by the TCR-pMHC affinity, but also by other cellular factors such as the T-cell’s differentiation state (e.g., naive or memory), self-reactivity of the TCR [35, 36], and the avidity effects driven by TCR/co-receptor clustering and pMHC density on antigen-presenting cells [37]. Although HERMES is more suited to predict molecular affinities, we show that its scores nonetheless can well track T-cell activities, making it a reliable tool for predicting T-cell response to antigens.

Lastly, we show that HERMES can be used to design novel peptides binding specific TCR-MHC complexes, achieving up to 50% experimental validation accuracy across diverse TCR-MHC systems. By leveraging our design algorithm, we characterize the specificity of a diverse range of TCRs, providing a quantitative measure for the diversity of the peptide-MHC antigens that a TCR can recognize in humans and mice.

### MODEL

T-cell response is mediated by the interactions between TCRs and pMHC complexes. To model the TCR-pMHC interaction, we use HERMES [27], a 3D rotationally equivariant structure-based neural network model, trained on the protein universe; see Fig. 1A. HERMES predicts the amino acid propensity from its surrounding 3D protein environment, a measure that we have previously shown to serve as a reliable proxy for assessing the effect of mutations on protein stability or binding [27, 28].

**Figure 1.**
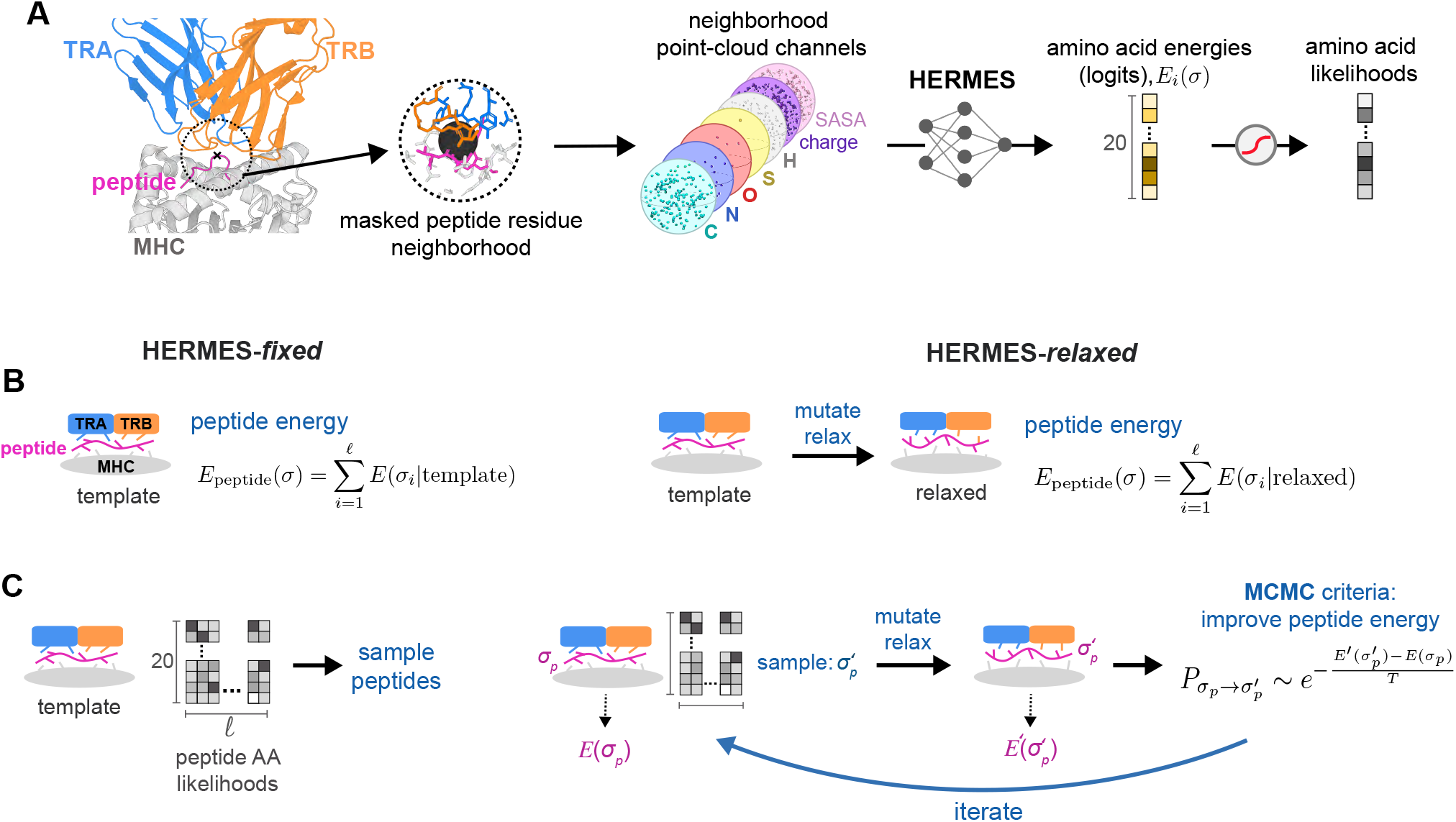
HERMES model for predicting and designing TCR-pMHC interactions. **(A)** HERMES is a structure-based, rotationally equivariant neural network pre-trained on the protein universe to predict amino acid preferences at a residue from its 3D structural environment [27, 28]. It takes as input all atoms within 10Å of a focal residue (with that residue’s atoms masked), featurizing them by atom type, charge, and SASA. HERMES then outputs energies *E*_*i*_(*σ*) for the 20 possible amino acids *σ* at residue *i*, which serve as logits in a softmax to estimate amino acid likelihoods. **(B)** To predict TCR-pMHC affinity, we use two protocols: HERMES-*fixed* (left) and HERMES-*relaxed* (right). In HERMES-*fixed* we use a fixed structural template closest to the TCR-pMHC of interest as input to HERMES, and evaluate the peptide energy as the sum of HERMES-computed per-residue energies for the peptide. In HERMES-*relaxed* mutations are introduced to the template’s peptide to match the peptide of interest, followed by local repacking in PyRosetta [29]. Peptide energy is then evaluated similar to HERMES-*fixed*, but using the relaxed structure. **(C)** We also use two protocols for peptide design. Similar to (B) we fix the closest structural template in HERMES-*fixed*, and use HERMES’ likelihood matrix for possible amino acids at each peptide position, which we sample to generate new peptide sequences. In HERMES-*relaxed* (right), we begin with a random peptide *σ*, and locally pack it with PyRosetta, and also generate an amino acid likelihood matrix (similar to HERMES-*fixed*) for this packed structure. We sample a new sequence *σ*′ from this likelihood matrix, and pack it with PyRosetta. We evaluate peptide energies *E*_peptide_(*σ*), and *E*_peptide_(*σ*′) for both peptides, using their respective PyRosetta-relaxed structures, and we use these energies as the criterion to sample improved sequences with MCMC (SI). This procedure is then iterated until convergence. Peptides designed by both approaches are then filtered by TCRdock [18] or Alpha-Fold 3 [30] to assure the stability of the complex.

For TCR-pMHC complexes, we seek to determine how changes in a peptide sequence impact the binding affinity of a TCR-pMHC complex, and ultimately, the T-cell response to the antigen. To do so, we characterize a score, which we term peptide energy, for a peptide *σ*, given its surrounding TCR and MHC structural environment,

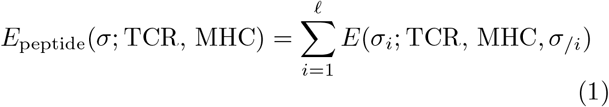

We assume that each peptide residue contributes linearly to the total energy by the amount *E*(*σ*_*i*_; TCR, MHC, *σ*_*/i*_), where *σ*_*i*_ is the amino acid at position *i* of the peptide. A residue’s energy contribution *E* is evaluated by HERMES (logits of the network), taking in as input the atomic composition of the surrounding TCR, MHC, and the rest of the peptide *σ*_*/i*_ (excluding the *i*^*th*^ residue) within a 10 Å of *σ*_*i*_’s *α*-Carbon; see Fig. 1 and SI for details.

To compute peptide energy, we input to HERMES an experimentally or computationally determined structure of a specified TCR-MHC complex bound to at least one peptide variant. Experimental data on the impact of peptide substitutions on the TCR-pMHC structure is limited, and computational models often fail to capture subtle conformational changes in the structure due to amino acid substitutions in a peptide. Given these constraints, we adopt two approaches to estimate peptide energies across diverse peptide variants for a given TCR-MHC complex:

- HERMES-*fixed*: In the simplest approach, we choose a TCR-pMHC structure with the same TCR and MHC as our query and a peptide sequence closest to the peptide of interest. This structure is then used as our template in HERMES to compute the peptide energy as described in eq. 1 (Fig. 1B). This method does not alter the underlying structure and assumes that peptide amino acid substitutions do not significantly change the conformation of the TCR-pMHC complex.
- HERMES-*relaxed*: Since amino acid substitutions can locally reshape a protein complex, we introduce a more involved protocol to account for these structural changes. We begin with the closest available structure for the TCR-pMHC complex. We then mutate the original peptide to the desired state and apply a relaxation procedure using PyRosetta to pack the substituted amino acid [29] (SI). During relaxation, side-chain and backbone atoms of the peptide are allowed to move, while only the side-chain atoms of the MHC and TCR chains within a 10 Å radius of the peptide are flexible. We then calculate the peptide energy with HERMES (eq.1), using the relaxed structure as input (Fig. 1B). Since PyRosetta’s relaxation procedure is stochastic, we average the peptide energy across 100 realizations of these relaxations. For ablation purposes, we also include results obtained by selecting the relaxed conformation with the lowest (most favorable) Rosetta energy, and using its HERMES peptide energy.

## RESULTS

### Predicting binding affinity of TCR-pMHC complexes

Binding between TCRs and pMHC complexes is necessary to mediate a T-cell response. The binding affinity of a natural TCR-pMHC complex is relatively low, with a dissociation constant *K*_*d*_ in the range of 1–100*µ*M [17, 43, 44], in contrast to nanomolar affinities for antibody-antigen complexes. Engineered TCRs can achieve much higher affinities in a nanomolar [45] to picomolar [46, 47] range.

Surface plasmon resonance (SPR) spectroscopy provides accurate measurements of the dissociation constant (*K*_*d*_) for specific TCR-pMHC systems. In these experiments, one protein–either the TCR or the MHC–is immobilized on a conducting plate, and the other is introduced in solution to bind to it. This binding alters the local refractive index near the plate, affecting the resonance signal. We used a published dataset of SPR-measured affinities for two TCR-MHC complexes binding to an ensemble of peptides, for which crystal structures of the complexes with at least one peptide are available [48, 49]; see Table S1 for details.

In our first example, we examined TCR 1G4 binding to NY-ESO-1 peptide variants, which is a cancer-testis antigen commonly expressed in many cancers and is targeted by immunotherapies [50]. We used two structural templates, one with the wild-type peptide SLLMWITQC (NY-ESO-1), and the other with a more immunogenic peptide, in which the Cysteine at position 9 is substituted by Valine (C9V) [38, 51] (Fig. 2A). For variants differing by a single amino acid, our predictions with HERMES*relaxed* correlated well (Spearman *ρ* = 0.72) with experimentally measured affinities from refs. [48, 49]; see Fig. 2B, C.

**Figure 2.**
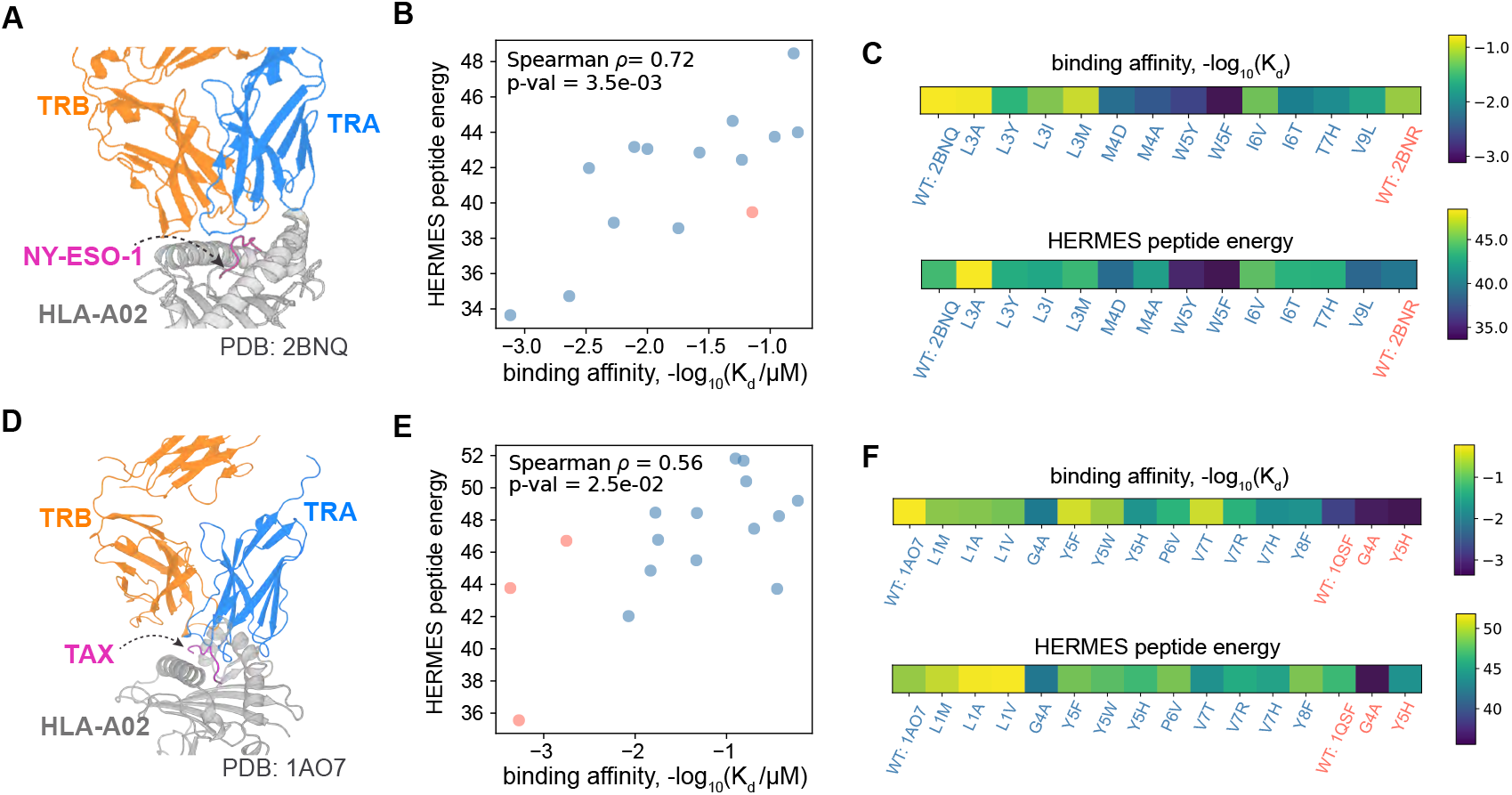
Predicting binding affinity of TCR-pMHC complexes. **(A)** The structure of the NY-ESO-1 peptide (purple), in complex with HLA-A*02:01 and a purified specific 1G4 TCR is shown (PDB ID: 2BNQ [38]). This structure together with that of a peptide variant with C9V mutation (PDB ID: 2BNR [38]) are used as templates to predict the binding affinity of the TCR-MHC complex to other peptide variants. **(B)** The binding affinities (*–*log_10_(*K*_*d*_*/µ*M)) of TCR-MHC complexes with 14 distinct peptides, measured using surface plasmon resonance, are shown against the predicted HERMES-*relaxed* peptide energies, using the closest structural template for each variant; 2BNQ is used for blue points and 2BNR for the orange point. The Spearman *ρ* between predictions and measurements is reported in the panel. **(C)** The heatmaps show the experimental affinity values (top) and predicted peptide energies (bottom) for all the peptide variants, with the identity of the mutations color coded based on the closest structural template. **(D-F)** Similar to (A-C) but for the Tax viral peptide in complex with HLA-A*02:01 and a specific human TCR. Two structural templates are used [39] for evaluating peptide energies with PDB IDs: 1AO7 (shown in D, and used for blue points), and 1QSF (used for orange points).

Next, we tested the A6 TCR binding to the HTLV-1 Tax peptide LLFGYPVYV. We employed two structural templates: one with the wild-type, the other with the mutant peptide with substitution Y8A [39] (Fig. 2D and Table S1). Again, the HERMES-*relaxed* peptide energies correlated well (Spearman *ρ* = 0.56) with the affinities, measured in ref. [49] (Fig. 2E, F).

Overall, HERMES predictions aligned closely with these limited experimental affinity measurements, with HERMES-*relaxed* showing a better agreement with affinity measurements than HERMES-*fixed*. We compared our approach for scoring peptides to alternative methods, including the baseline substitution matrices (BLOSUM62 [40] and the TCR-specific substitution matrices from ref. [52]), the score from the TCR-pMHC structure prediction algorithm TCRdock [18], the sequence-based machine learning algorithms for TCRs (TAPIR [41], NetTCR [53], and TULIP [16]), and the structure-based inverse folding models (ESM-IF1 [42] and Protein-MPNN [31]); see Table I for key benchmarking results and Table S2 and Fig. S1 for a more extensive comparison of different methods, including the different HERMES models, and different scoring schemes by ProteinMPNN.

**Table I.**
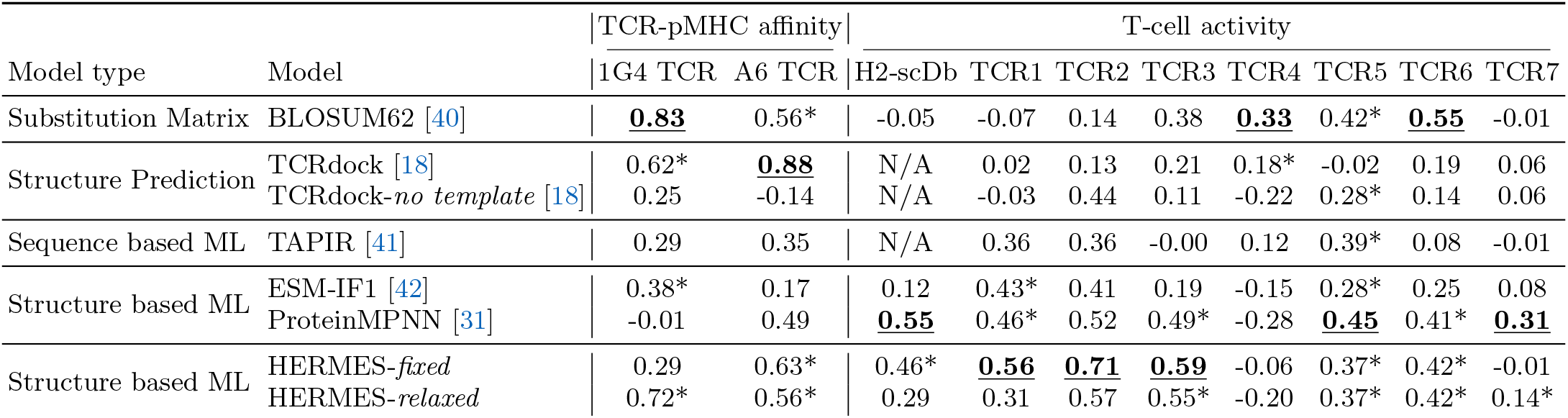
Benchmarking of models for predicting TCR-pMHC binding affinities and T-cell activities. The table lists the Spearman *ρ* of models’ predictions against experimentally measured binding affinities (Fig. 2) and T-cell activities (Fig. 3) in response to peptide variants. The HERMES-*fixed* and HERMES-*relaxed* performance are from models with no noise (0.00) in training data, and the ProteinMPNN performance is from the model with a small noise amplitude of 0.02 Å; see SI for details. We show in **bold** the best model for each dataset, and any model whose performance is *not* significantly different from the best model (p-value*>* 0.05 associated with the difference in the Fisher’s z-transformed Spearman correlations) is indicated with an asterisk (*). See Tables S2 and S3 in the SI for *p*-values and additional models.

For the 1G4 TCR, BLOSUM62 [40] achieves the highest correlation with experimental affinity data, whereas template-guided TCRdock [18] leads for the A6 system; in contrast, TCRdock without using a homologous structure as a template underperforms. In both systems, HERMES achieves performance that is statistically indistinguishable from the top performers (Fisher’s z-test p-value *>* 0.05). Overall, BLOSUM62, template-guided TCRdock, and HERMES rank strongly as predictors of TCR-pMHC affinities across the tested datasets. In addition, for the 1G4 system, ESM-IF1 [42] and the TCR-specific substitution matrices from ref. [52] also perform competitively. However, due to the limited number of affinity measurements, we lack statistical power to resolve finer differences among the models.

It should be noted that although BLOSUM62 is one of the best contenders for affinity prediction, its sequence-averaged construction ignores local structural context and therefore assigns identical substitution scores regardless of a residue’s environment (Fig. S2); similar limitations apply to the TCR-specific substitution matrices from ref. [52]. HERMES, by contrast, is structure-aware and can attribute distinct scores to the same mutation when it occurs in different conformational settings–an ability that is critical for nuanced mutational forecasts (Fig. S2). Determining the biophysical circumstances under which such context-agnostic scoring schemes suffice, and when structure-aware approaches become necessary, will require systematic evaluation on broader, more diverse datasets.

Lastly, we probed HERMES’s robustness to potential inaccuracies in structures generated by the widely used AlphaFold3 (AF3) algorithm [30], replacing the crystal structure inputs with AF3 models; see SI for details. Both the template-guided and the de novo AF3 structures remained within 1.1 Å RMSD of the respective crystal structures (Table S4). For both HERMES and ProteinMPNN, replacing experimental structures with AF3 predictions altered predictions for the 1G4 and A6 TCRs binding to pMHC variants, but the changes were not statistically significant (Fisher’s z-test p-value *>* 0.05); see Fig. S4. It is nonetheless expected that the predictions should suffer in the absence of reliable structural templates, as seen also in other methods such as TCRdock [18] (Tables I, S2).

### Predicting T-cell response to peptide antigens

T-cell activation is shaped by factors beyond TCR-pMHC affinities, including the T-cell’s differentiation state, its cross-reactivity to self-antigens [35, 36], and avidity effects from TCR/co-receptor cross-linking [37]. Nonetheless, because readouts of T-cell activiy (e.g., T-cell proliferation, and production of activation markers) are easier to obtain than SPR-based affinity measurements, we next asked whether HERMES scores, which are more attuned for predicting TCR-pMHC binding affinity, can serve as predictors of T-cell activities.

As the first system, we examined T-cell responses to all single-point mutants of a native peptide, as measured in ref. [52] for different TCR-pMHC systems. Specifically, we make predictions for (i) three TCRs (TCRs 1-3) recognizing variants of a highly immunogenic HLA-A*02:01-restricted human cytomegalovirus (CMV) epitope NLVPMVATV (NLV), (ii) three TCRs (TCRs 4-6) recognizing variants of the weaker HLA-A02:01-restricted melanoma gp100 self-antigen IMDQVPFSV, and (iii) a TCR (TCR 7) recognizing variants of weakly immunogenic HLA-B*27:05-restricted pancreatic cancer neo-peptide GRLKALCQR, believed to have elicited an effective immune response in long-term cancer survivors [52]. T-cell activity was quantified by determining the half-maximal effective concentration (EC_50_) from the dose-response curves of 4-1BB^+^ CD8^+^ T-cells across varying peptide concentrations; see Methods on details of inferring EC_50_.

With the exception of TCR1 (where a TCR-pMHC template with HLA-A*02:01 and one peptide variant was already available), we relied on AlphaFold3 (AF3) [30] to model the remaining TCR-pMHC complexes; see Table S1 and Dataset S1 for template information and AF3 sequence inputs for TCRs 2-7. As shown in Figs. 3A, and S3, HERMES-*fixed* peptide energy predictions (eq. 1) correlate well with experimental EC_50_ measurements for the CMV peptide variants, reaching up to Spearman correlation *ρ* = 0.71 for TCR2, and outperforming the alternative models. For TCR1 and TCR3, the performance of ProteinMPNN [31] is worse but statistically indistinguishable from HERMES (Tables I, S3, and Fig. S1). Moreover, replacing the crystal structure for TCR1 with an AF3 model yields comparable perfor-mance, underscoring HERMES’s robustness to structural inputs so long as the structural models are of high quality (Figs. S5, S4). Correlations are lower for TCRs 4-6 targeting variants of the melanoma gp100 self-antigen, with HERMES achieving Spearman correlation *ρ* = 0.42 for TCR6 as its best prediction in this class, with an accuracy comparable to ProteinMPNN [31] (Fig. 3B and Tables I, S3). HERMES underperforms for TCR7 responding to the pancreatic cancer neo-peptide GRL-KALCQR (Fig. 3C), for which only the ProteinMPNN’s predictions show significant albeit weak correlation with data (Table I). The reduced accuracy of HERMES for cancer epitopes compared to the variants of the viral CMV epitope could reflect the impact of antagonism from self-antigens similar to tumor epitopes, hindering T-cell responses [35, 36]—though additional experiments are needed to confirm this.

**Figure 3.**
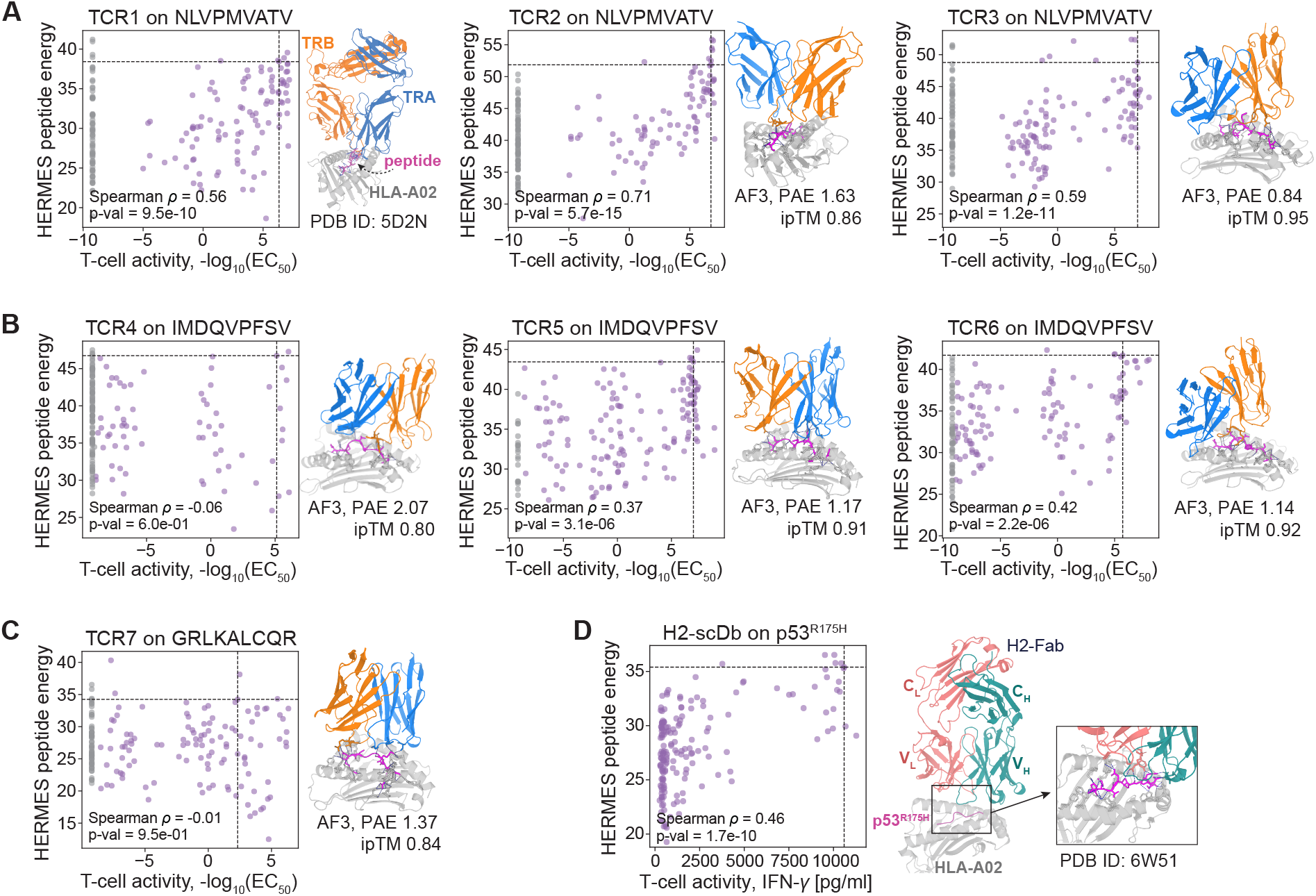
Predicting T-cell activity in response to peptide variants. **(A-C)** The predicted HERMES-*fixed* peptide energies versus experimentally measured T-cell activity are shown for (A) HLA-A*02:01-restricted CMV peptide variants interacting with TCR1-TCR3 (panels), (B) HLA-A*02:01-restricted melanoma gp100 peptide variants interacting with TCR4-TCR6, and (C) HLA-B*27:05-restricted pancreatic cancer peptide variants interacting with TCR7. In each case, T-cell activity in response to all single amino acid mutations from the wild-type is measured. T-cell activity is based on fitted EC_50_ values from dose-response curves of 4-1BB^+^ CD8^+^ T-cells (SI; data from [52]). Gray points indicate peptides with T-cell responses below detection limit (no EC_50_ obtained) and are excluded from the reported Spearman correlations. Except for TCR1 (resolved structure, PDB ID: 5D2N [54]), the other TCR-pMHC complexes were modeled with AF3, and the model’s confidence for each wild-type peptide is denoted. These structures are used as templates to estimate peptide energies with HERMES-*fixed*; see Table S1 and Dataset S1 for details on the templates. In all panels, TCR-*α* (TRA) is shown in blue, TCR-*β* (TRB) in orange, the peptide in magenta, and HLA in gray. **(D)** The predicted HERMES-*fixed* peptide energies are shown against experimentally measured T-cell activity for single point mutants of the HLA-A*02:01-restricted p53^R175H^ neoepitope from ref. [5]. T-cell response (IFN-*γ* production) is mediated through a bispecific antibody binding to the pMHC complex. The co-crystallized structure of the bispecific antibody with the p53^R175H^-HLA-A*02:01 complex (PDB ID: 6W51 [5]) is used as the template for HERMES-*fixed* predictions. For all systems (A-D), the heatmaps for the predicted and the experimentally measured mutational effects are shown in Fig. S3.

Sequence-based models (TAPIR [41], NetTCR [53], and TULIP [16]) are outperformed by structure-based models tested here in all cases except for TCR4, which has the lowest AF3 confidence at the TCR-pMHC interface (i.e., a lower confidence in its fold); see Table S4 and Fig. S4 for the AF3 confidence metrics for all systems. Here, even the basic substitution models (BLO-SUM62 [40] and the TCR-specific matrices from ref. [52]) outperform the structure-based predictors (Tables I, S3). This suggests that structural approaches hinge on a reliable structure as input, though more data is needed to draw definitive conclusions (Fig. S4).

Next, we examined the therapeutic T-cell responses to all single-point mutants of the HLA-A*02:01-restricted p53-derived neoepitope p53^R175H^, HMTEVVRHC, measured in ref. [5]. This T-cell therapy approach used a designed bi-specific Fab antibody that binds both to the TCR-CD3 complex on T-cells and to the variants of the p53^R175H^-MHC complex on tumor cells (Table S1). A strong binding to an epitope activates the engaged T-cell, and the extent of such activity was measured by the production of IFN-*γ*. The resulting HERMES-*fixed* predicted peptide energies (eq. 1) show a Spearman *ρ* = 0.46 with the experimentally measured IFN-*γ* production (Figs. 3D, S3). ProteinMPNN achieves a slightly higher correlation (Spearman *ρ* = 0.55), but this improvement over HERMES is not statistically significant (Fisher’s z-test p-value *>* 0.05); see Table I. Lastly, we found that replacing the crystal structure with the AF3 model substantially degrades HERMES and ProteinMPNN performances (Fig. S4, Tables I, S3), as the AF3 model consistently mis-docks the bispecific antibody on the pMHC complex (Table S4 and Fig. S5).

Across the limited number of systems examined, our analyses indicate that–when a suitable structural template is available–HERMES can estimate epitope mutational effects at the pMHC-TCR interface and, in the H2-scDb case, at a Fab-pMHC interface. In this benchmark, ProteinMPNN performs comparably to HERMES, indicating that inverse-folding algorithms not explicitly trained for this task may nonetheless capture the impact of peptide mutations on T-cell responses. However, definitive conclusions about the generalizability of these methods to neoepitopes will require larger datasets with broader mutational coverage across peptide variants. Importantly, predictive performance for both approaches depends on access to a high-quality structure (experimental or computational), which limits applicability in systems without suitable structural models (Fig. S4); see SI and Tables I and S3 for a detailed benchmark of different approaches, including different HERMES models.

### Design of novel peptides reactive to a TCR-MHC complex

TCR-engineered T-cell therapies are designed to target defined neoantigens, but specificity remains a central challenge. Engineered high-affinity TCRs can cross-react with untested self-peptides, causing severe off-target toxicity; in a MAGE-A3 directed trial, recognition of a Titin-derived peptide led to fatal cardiac events [59]. Therefore, developing approaches to predict the distribution of peptides recognized by engineered TCRs could enable more stringent preclinical screening and help mitigate off-target risk in patients.

To map the peptide-recognition landscape of TCRs, we present a structure-guided pipeline for designing immunogenic peptides predicted to elicit strong TCR recognition when presented by MHC class I. Leveraging HERMES’s ability to predict T-cell responses to peptides presented by MHC-I, our approach begins with an existing template structure–experimentally or computationally resolved—of a TCR-MHC in complex with at least one peptide variant, which we refer to as the wild-type peptide. We then generate candidate peptides using two strategies, with different degrees of complexity; see Fig. 1C for a schematic description of the two pipelines.

In the basic pipeline (HERMES-*fixed*), we sequentially mask the atoms of each amino acid along the peptide, one at a time. Using the HERMES model, we predict the probability of each amino acid type for a given residue, based on the local structural environment within 10 Å of the masked residue. Repeating this procedure for every amino acid along the peptide yields a position weight matrix (PWM) that represents the probabilities of different amino acids at each peptide position. Peptides are then sampled by drawing from this PWM. It should be noted that HERMES-*fixed* retains the structural fingerprints of the original peptide, as it does not account for local structural changes induced by amino acid substitutions; see Fig. 1C and S6A, and SI for more details.

To incorporate structural flexibility and relaxation following amino acid substitutions, we introduce the design pipeline of HERMES-*relaxed*, using simulated annealing and Markov chain Monte Carlo (MCMC) sampling. We begin with the template TCR-MHC structure and a completely random peptide sequence, which is packed *in silico* within the structure using PyRosetta’s packing functionality, followed by the Relax protocol [29]. During relaxation, both side-chain and backbone atoms of the peptide are allowed to move, while only the side-chain atoms of the MHC and TCR chains within 10 Å of the peptide are flexible. With this relaxed structure, we apply the HERMES-*fixed* method to generate a PWM and sample a new peptide, which is then packed and relaxed to form a new structure. We use MCMC to determine whether to accept the newly sampled peptide by comparing its HERMES peptide energy (favorability), evaluated in its relaxed conformation, to that of the original peptide (eq. 1). We then iteratively perform this procedure, incrementally reducing the MCMC temperature at each step, until the results converge; see Fig.s 1C, S6B-C, and SI for more details.

The peptides designed by both approaches are then filtered by TCRdock [18] or AlphaFold3 [30], using the Predicted Alignment Error (PAE) between TCR and pMHC interfaces, to assure folding and binding of the resulting TCR-pMHC structures. For each system, we define a specific PAE threshold close to the TCRDock PAE of the system’s TCR-MHC in complex with the native (wild-type) peptide, which is known a priori to activate T-cells. As a result, the PAE thresholds vary slightly across different systems; see SI for details.

We tested the accuracy of our peptide design pipeline in three systems, with the structural templates taken from (i) the cancer-testis antigen NY-ESO-1 in complex with the TCR 1G4 and HLA-A*02:01, (ii) a peptide derived from the Epstein-Barr virus (EBV) in complex with the TK3 TCR and the HLA-B*35:01, and (iii) the immuno-therapeutic target MAGE-A3 in complex with an engineered TCR and HLA-A*01:01. For brevity, we refer to these three systems as NY-ESO, EBV and MAGE, respectively; See Table S5 for information on the structure templates used in each case. These systems were selected because each one has high-resolution crystal structures of the same TCR-MHC scaffold bound to at least two distinct peptide variants, and that they span a therapeutically relevant spectrum that includes both engineered (for MAGE and NY-ESO) and natural (for EBV) TCRs directed against viral as well as tumor antigens. It should be noted that as we progressed from NY-ESO to EBV and MAGE, we chose to explore the sequence space farther from the wild-type peptides, leading to more challenging designs.

We validated our designs using Jurkat cells expressing a specific TCR and an endogenous NFAT-eGFP reporter to indicate T-cell activation in response to different peptides. We tested the TCR-MHC specificity in all systems–NY-ESO, MAGE, EBV–under 96 different conditions, including the de novo designed peptides, positive controls (wild-type peptides from the template structures), and an unstimulated “no peptide” control. Peptides were presented on artificial APCs (aAPCs), expressing the specified HLA in each system. For NY-ESO designs, we determined the the percentage of GFP-positive cells through both flow cytometry and fluorescence microscopy. Given the consistency between the two exper-imental approaches (Fig. S7) we relied on fluorescence microscopy only to measure GFP levels induced by different peptides for MAGE and EBV designs; see SI for experimental details and Dataset S2 for data on these measurements.

We used two structural templates for the NY-ESO system: one containing the original 9-amino acid peptide SLLMWITQC, and another with the more immunogenic C9V substitution [38, 51]. Our de novo designs differed from the closest wild-type peptide by 2 to 7 amino acids, with a TCRDock PAE *≤* 5.5. For this system only, we selected 35 negative designs (designed by HERMES, but with PAE *>* 5.5) for experimental testing: only one of these was a false negative, which interestingly, had a desirable AF3 PAE score (as low as the wild-type peptides); see Figs. S8, S9. For the 58 positive designed peptides (PAE *≤* 5.5), we achieved an overall design accuracy of 50%, with GFP levels significantly higher than the negative control; see SI for details. The accuracy of predictions decreases as the sequence divergence of the de novo designs from the wild-type peptides increases (Fig. 4A, S9). HERMES-*fixed* achieved higher design accuracy of 70% within its smaller designed sequence subspace of up to 5 substitutions from the wildtypes, whereas HERMES-*relaxed* explored sequences further from the wild-type peptides (4-7 substitutions) but with a reduced accuracy of 24% (Fig. 4A). The main difference between the true and false positive sequences was at position 8–from strongly preferring glutamine within the true positives, to having similar preferences among glutamine, glycine, arginine, or methionine within the false positives (Fig. 4A).

**Figure 4.**
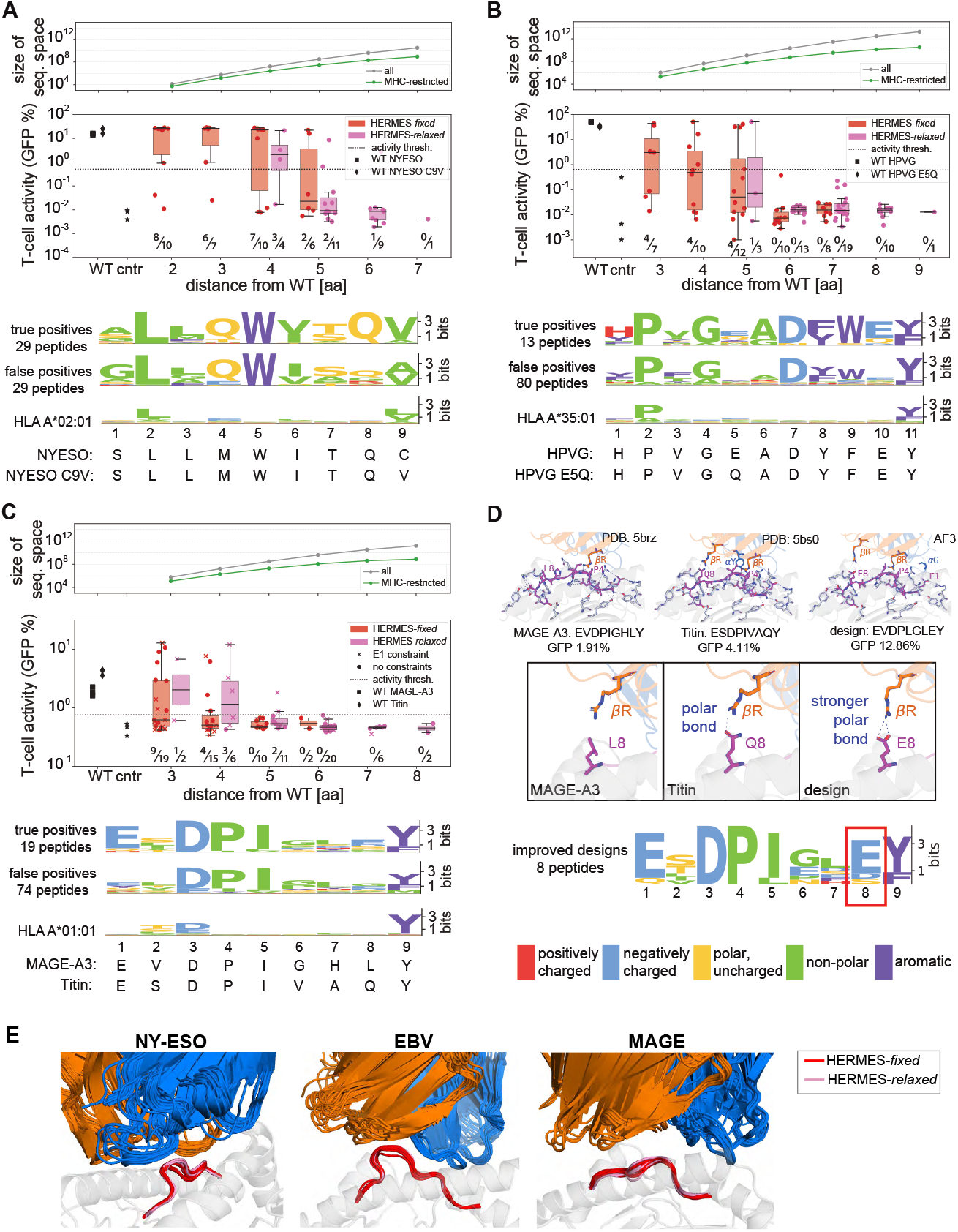
De novo peptide design to elicit a T-cell response. **(A-C)** Validation results are shown for designed peptides built upon the wild-type templates from (A) NY-ESO peptide against TCR 1G4 and HLA-A*02:01 (PDB IDs: 2BNR, 2BNQ [38]), (B) EBV epitopes against the TK3 TCR and the HLA-B*35:01 (PDB IDs: 3MV7 [55], 4PRP [56]), and (C) MAGE-A3 and Titin epitopes against an engineered TCR and HLA-A*01:01 (PDB IDs: 5BRZ, 5BS0 [57]). Designs were validated using Jurkat cells with endogenous NFAT-eGFP reporter expressing the paired TCR, interacting with peptide presented by the MHC in each system. The reported T-cell activities (percentage GFP-positive) are averaged over three replicates. The box plots show T-cell activity for designed peptides at different Hamming distances from the wild-type; the black line is the median, the box spans the 25-75% quantiles, and dots indicate individual data points. Red and pink boxes/markers correspond to designs with HERMES-*fixed* and HERMES-*relaxed* models, respectively; square markers denote wild-type peptides, and stars are negative controls (no peptide). The dashed line marks the activation threshold, and numbers below each bar show the relative number of successful designs out of total designs. The top plot in each panel shows the size of the peptide sequence space at different distances from the wild-types, with the gray line indicating the complete sequence space, and the green indicating the typical number of sequences likely to be presented by the MHC of each system; see SI for details. The bottom shows sequence motifs as PWMs for (i) true-positive designs that induced T-cell activation, (ii) false positives that did not, and (iii) the MHC presentation motif from the MHC Motif Atlas [58]. **(D)** 3D structures of TCR–pMHC complexes focused on the peptide are shown for MAGE-A3 (PDB ID: 5BRZ), Titin (PDB ID: 5BS0), and the highest-activity design in (C) modeled by AF3 [30]. The enhanced activity for Titin versus MAGE-A3, and for the designed peptide versus both wild-types follows from the formation of polar bonds between position 8 of the peptide (magenta) and the CDR3 loop of TCR*β* (orange). The PWM highlights the prominence of E_8_ among the eight designs with improved T-cell activity in (C). **(E)** Structures predicted with AF3 for the activity-inducing designed peptides (above the dashed lines in A-C) are displayed in complex with their cognate TCR-MHC’s. For each system, designs generated by *HERMES-fixed* (red) and *HERMES-relaxed* (pink) are overlaid after aligning the MHC molecule (in gray). Slight shifts are seen in the backbone conformations of the peptides and the TRA (blue) and TRB (orange) loops.

For the EBV system, we used two structural templates: one with the 11-amino acid peptide HPVGEADYFEY— commonly referred to as HPVG—derived from the viral latent nuclear protein EBNA-1, and the other with an epitope from a wide-spread viral variant, with a glutamine at position 5 [56]. Unlike NY-ESO, the HPVG peptide has a more flexible conformation with a helical turn within the MHC binding pocket. Our de novo designs differed from the closest wild-type peptide by 3 to 9 amino acids, with a TCRDock PAE *≤* 5.1. Across the 93 de novo designs, we achieved an overall accuracy of 14%, associated with the peptides that induced significant GFP levels in T-cells. The accuracy of predictions decreases with increasing the sequence divergence from the wild-types (Fig. 4B, S10). Notably, we achieved 40% accuracy among the designs within 3-5 amino acid distance of one of the wild-type templates, while none of the designs with larger than 5 amino acid distance were successful. Lastly, we observe a strong preference for glutamic acid at position 10 of the successful designs relative to the false positives (Fig. 4B).

For the MAGE system, we used two structural templates: one with the 9-amino acid MAGE-A3 peptide EVDPIGHLY and another with the Titin-derived selfpeptide ESDPIVAQY, which is expressed in cardiomyocytes [57]. Our de novo designs differed from the closest wild-type peptide by 3 to 8 amino acids, with a TCRDock PAE *≤* 5.2 (Fig. 4C). From the resulting 93 de novo designs, 20% activated T-cells with significant levels of GFP expression. Similar to the other systems, the accuracy of predictions decreases with increasing the sequence divergence from the wild-types (Fig. 4C). We achieved 30% accuracy among the designs with 3-5 to amino acid distance, while none of the designs with larger than 5 amino acid distance from the wild-types were successful.

Fig. 4C shows differences in amino acid composition between successful and failed designs. Alanine and glycine scanning experiments have previously suggested that MAGE-A3 peptide variants with an E-DPI- - -Y motif activate T-cells in this system [60]. Consistent with these findings, the fraction of T-cell activating variants in our designs increases as their sequences more closely resemble this motif (Fig. S11).

In a subset of designs using the MAGE-A3 template, we fixed the glutamic acid at position 1 (E_1_), which is part of the E-DPI—Y pattern. In both MAGE-A3 and Titin E_1_ is within 4 Å of the MHC and forms strong polar interactions with it (Fig. 4D). Fixing E_1_ improved the design accuracy to 1*/*3, compared to 1*/*9 when this amino acid was not constrained. Even without this constraint in the design protocol, E_1_ appeared in 2*/*3 of successful designs, and in none of the unsuccessful ones (Fig. S11). Notably, among the more distant designs made with HERMES-*relaxed*, only those containing a fixed E_1_ were able to activate T-cells (Fig. 4C). This suggests that constraining essential amino acids can enable broader exploration of sequence space at other positions, possibly due to epistatic interactions. However, further investigation is needed to confirm this.

Several of our designs outperformed the wild-type peptides in activating T-cells: eight surpassed Titin’s GFP levels, and eleven exceeded MAGE-A3’s (Fig. 4C). These improved designs strongly favor glutamic acid (E) at position 8 of the peptide, where MAGE-A3 has leucine (L), and Titin has glutamine (Q) (Fig. 4D). In Titin, the glutamine forms a polar bond with an arginine (R) in the interacting TCR*β*, and it is likely responsible for increasing Titin’s activity relative to MAGE-A3’s. The glutamic acid substitution in our designs further strengthens this polar bond (Fig. 4D), resulting in a stronger T-cell response. For NY-ESO and EBV, the wild-type peptides were strongly immunogenic and induced high levels of GFP, and many of our designs induced comparable T-cell responses, but did not improve upon those (Fig. 4).

To gauge the impact of using AF3 structures as design templates, we reran the HERMES-*fixed* protocol with AF3 predicted structures (with and without template guidance), and obtained peptide PWMs that were almost identical to those derived from using crystal structures (SI and Fig. S12). An interesting exception is the MAGE system, where using the AF3 template introduced a strong preference for glutamic acid at positing 1 of the peptide (Fig. S12C), mirroring the pattern observed among the designed peptides that induced strong T-cell response (Fig. 4D).

Although no direct experimental comparisons were performed, computational evaluations using TCRdock PAE and presentation score by MHC-I alleles indicate that HERMES designs outperform or match those generated by ProteinMPNN [31], the corresponding MHC position weight matrix [58, 61–63], or the BLOSUM substitution matrix [40] (SI and Figs. S13, S14, S15). A notable exception is in the MAGE system, where ProteinMPNN designs generally have more favorable PAE scores, albeit with a lower proportion of designs passing the antigen presentation filter by the MHC molecule (Fig. S15).

In all the three systems we used the PAE values from TCRdock and AF3 as filtering criteria to select designs for experimental validation (see SI). The PAE values reported by the structure prediction algorithms have been shown to be noisy indicators of protein folding fidelity and design quality [64]. While larger experimental libraries are needed to draw definitive conclusions, it appears that PAEs from TCRdock and AF3 have some complementary predictive strengths (Figs. S8, S16), and therefore, combining their information could lead to a more robust design decisions (SI). Both of these scores correlate only weakly with HERMES (Figs. S8, S13, S14, S15), so they can provide complementary design filters. Furthermore, designed peptides with higher (more favorable) HERMES energies more often activated T-cells (Fig. S16), suggesting that a more restrictive sampling (e.g., with a lower MCMC temperature for HERMES-*relaxed* protocol) could yield a higher success rate, albeit at the expense of peptide diversity.

Overall, HERMES demonstrates a potential for designing highly immunogenic peptides up to five amino acid substitutions away from the wild-type–a task that would otherwise require exploring a vast sequence space of roughly 10^8^ to 10^10^ possibilities for a 9-residue to an 11-residue peptide (Fig. 4A-C). This capability raises an important question: to what extent do these substantial changes in sequence alter the binding geometry of the peptide inside the MHC-TCR cleft? To this end, we used AF3 to model the TCR-pMHC interfaces for the experimentally validated immunogenic designs from the three systems. Although we see slight variation in both peptide and TCR backbone conformations, overall, we found that our designed immunogenic peptides adopt conformations very close to their wild-type templates, even when those designs were generated with HERMES-*relaxed* (Fig. 4E). We attribute this structural convergence to two potential factors: (i) residual limitations in HERMES peptide-backbone sampling strategy, and (ii) an inherent AF3 bias toward conformations it encountered during training. Given that TCRs can cross-react with peptides of markedly different conformational poses [65], developing methods that can better explore peptide conformational space within the TCR-MHC groove remains a compelling direction for future research.

### Structural basis for TCR specificity

TCRs exhibit substantial degeneracy in their recognition, with some autoimmune T-cells experimentally shown to recognize over one million distinct peptides [67]. However, the full extent of TCR degeneracy for typical receptors remains unclear, largely due to the limitations in high-throughput experimental assays for TCR recognition of many peptides presented on different MHCs.

Our computational peptide design framework helps address this limitation, at least for the TCR-MHC complexes with known structural templates. Specifically, for a given TCR-MHC pair, we can use HERMES-*fixed* protocol to generate a peptide position weight matrix (PWM), representing the ensemble of peptides presented by the MHC and recognized by the TCR. We use the entropy of this PWM as a proxy for TCR degeneracy.

Fig. 5A shows the entropy of these peptide distributions for 105 TCR-MHC pairs across 10 human MHC-I alleles; see Table S6 for details on these structures. The median peptide entropy across these TCR-MHC pairs is approximately 10 bits, indicating that a typical TCR-MHC pair can recognize on the order of *≃* 10^3^ peptides. In contrast, examining the degeneracy of MHC-I presentation alone in humans–using the peptide PWMs gathered from the MHC Motif Atlas [58]–yields a median entropy of 31 bits, implying that MHC-I molecules can typically recognize *≃* 2 *×* 10^9^ distinct peptides (Fig. 5A). A similar analysis on 36 TCR-MHC pairs spanning three mouse MHC-I alleles shows comparable trends, with a MHC-I degeneracy on the order of *≃* 5 *×* 10^8^ peptides (29 bits), and the TCR-MHC degeneracy of *≃* 10^3^ peptides (10 bits); see Fig. 5B, and Table S6 for details on these structures. This pronounced reduction in entropy— observed by comparing the distribution of peptides recognized by MHC-I molecules alone to that recognized by TCR-MHC-I complexes—quantifies how TCRs constrain the accessible antigenic shape space, as illustrated by the polar interactions that they form with peptides in Fig. 5C.

**Figure 5.**
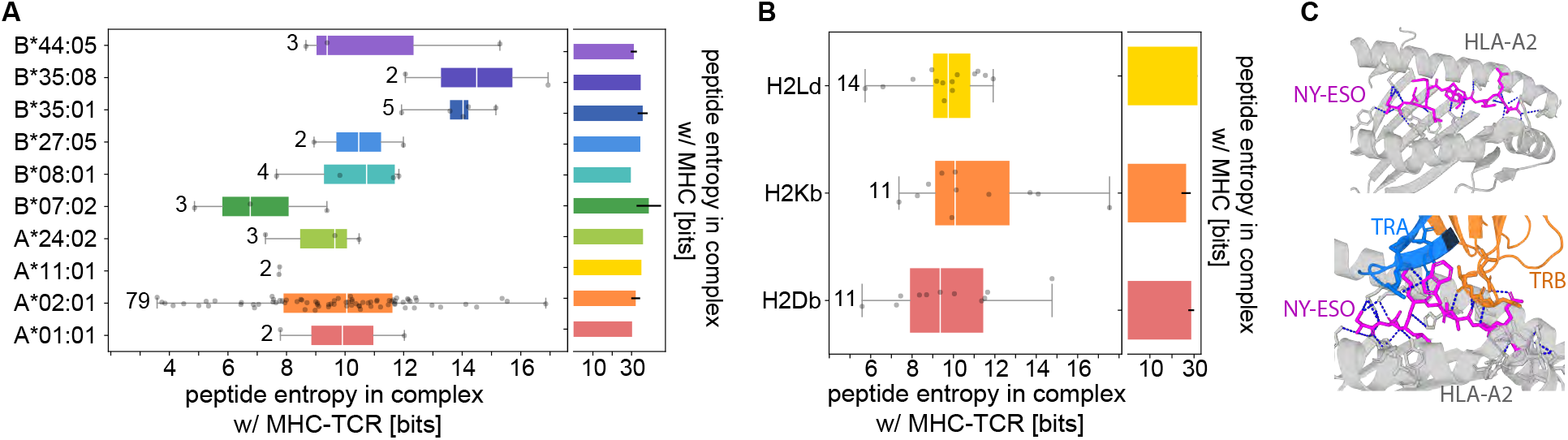
Evaluating T-cell specificity from de novo designed peptides. **(A)** T-cell specificity is measured by peptide entropy, which is computed from the HERMES-*fixed* predicted amino acid likelihood matrix, starting from a TCR-pMHC structural template. Each point on the plot corresponds to the entropy estimate from a distinct TCR-pMHC template, grouped by their MHC allele. The box plots display the distribution of peptide entropies for TCR-pMHC templates sharing the same MHC allele, with a white line marking the median and the box spanning the 25% to 75% quantiles. The number on the side of each box indicates the number of distinct structural templates we used for that MHC class; see Table S6 for details on these structures. The side plot shows peptide degeneracy for MHC presentation alone, evaluated as the entropy of the peptide position weight matrices gathered from the MHC Motif Atlas [58]. The error bars indicate the standard deviation of peptide degeneracies across motifs of different lengths, associated with a given MHC allele. **(B)** Similar to (A) but for degeneracy of peptides interacting with TCRs and MHCs in mice. **(C)** Polar bonds (blue dashed lines) between the NY-ESO peptide (magenta) and HLA-A*02:01 (gray) is shown at the top (PDB ID: 1S9W [66]). The additional polar bonds formed between the peptide and TRA (blue) and TRB (orange) of an interacting TCR are shown at the bottom (PDB ID: 2BNR [38]). These additional polar bonds limit the degeneracy of the peptides that can interact with a TCR-MHC complex compared to those presented by the same MHC.

As an alternative to MHC Motif Atlas [58], we explored the ability of HERMES-*fixed* to model MHC degeneracy. When structural coverage is extensive (roughly *>* 30 structures of distinct peptides for a given MHC allele), the motifs produced by HERMES align closely with those in the MHC Motif Atlas (SI, Fig. S17, and Table S7). With sparser structural coverage, HERMES underestimates the peptide diversity, likely because backbone conformations are insufficiently sampled. Given the imbalanced distribution of structures across MHC alleles, we therefore continue to use MHC Motif Atlas [58] as our reference for computing MHC motif specificity.

Because our analyses focus on interactions involving typical TCR and MHC molecules, our entropy-based degeneracy estimates are smaller than those from previous theoretical studies, which often relied on experimental observations from autoimmune T-cells or limited synthetic peptide libraries [68]. Moreover, the HERMES-*fixed* model generates PWMs based on peptides whose backbone conformations match those of the wild-type template, likely underestimating the true degeneracy of TCR-MHC complexes (Fig. 5). Although HERMES-*relaxed* can partially address this issue, its designs are only reliable within roughly five amino acids of the native peptides with similar backbone conformations (Fig. 4), preventing a full exploration of the possible peptide poses for a given TCR-MHC complex. Nonetheless, our estimates can serve as informative lower bounds on typical MHC-I and TCR degeneracies.

## DISCUSSION

In this work, we introduced a structure-based machine learning model to predict TCR-pMHC interactions and to design novel reactive peptides for TCR-MHC complexes. Our approach builds on HERMES, a physically motivated equivariant neural network model that we previously developed [27, 28]. Trained on the protein universe, HERMES predicts amino acid propensities based on their atomic environments in protein structures, and has been shown to implicitly learn biophysical interatomic interactions in proteins [28], enabling it to make highly generalizable predictions. This generalizability is particularly important for TCR-pMHC complexes, where only a modest number of experimentally resolved structures are available, and computational algorithms often struggle to accurately predict new structures [18, 19].

We found that HERMES-predicted peptide energies closely tracked both TCR-pMHC binding affinities and peptide-induced T-cell activities across diverse systems, including variants from a CMV-derived viral epitope [52], the NY-ESO cancer-testis antigen [48, 49], and a melanoma self-antigen [52]. Notably, HERMES also accurately predicted the activity of p53-derived neoepitope variants in the context of a bispecific antibody-mediated T-cell therapy [5]. This result suggests the algorithm’s potential applicability: given a reliable structural template close to the system of interest, HERMES can predict peptide-induced T-cell responses, whether through direct TCR interactions or antibody-mediated mechanisms. However, drawing more definitive conclusions about the generalizability of HERMES would require examining a larger datasets of TCRs interacting with pMHC variants.

Based on the HERMES predictions for TCR-pMHC reactivities, we presented a computational protocol for de novo design of immunogenic peptides that can elicit robust T-cell responses. Starting from the structural template of a TCR-MHC with a native (wild-type) peptide, we developed an MCMC search guided by HERMES peptide energies to explore the vast sequence space of roughly 10^8^ (10^10^) possibilities for 9-residue (11-residue) peptides, to identify immunogenic candidates. Experimental validation across three systems—with native peptides from NY-ESO and MAGE cancer epitopes, plus a viral epitope from EBV—revealed that our designs reliably activated T-cells, even with up to five substitutions from the native peptide used in the structural template. Notably, several MAGE-derived variants outperformed or matched the native peptide in T-cell activation, demonstrating the method’s potential to enhance peptide potency.

This capability can be relevant to personalized peptide-based cancer vaccines, which aim to induce durable T-cell memory against a patient’s tumor antigens to reduce the risk of rebound [6, 8, 69]. Despite their promise, the success of cancer vaccines has been limited by low peptide immunogenicity and poor uptake by antigen-presenting cells [70]. Recent approaches have employed deep mutational scanning experiments to identify more immunogenic cancer neoantigens against specific TCR candidates [52]. However, these experiments are often limited to scanning single amino acid mutations from the wild-type peptide due to the combinatorial explosion when exploring multiple mutations. Moreover, the precise TCRs responding to peptide vaccines in a patient are often unknown, and many immune responses are induced de novo [69]. With the caveat of access to suitable structural templates, HERMES could inform T-cell vaccine design by computationally generating diverse peptide libraries predicted to engage a broad set of candidate TCRs (identified independently), nominating vaccine epitopes for subsequent safety and efficacy testing.

Accurately modeling T-cell specificity remains challenging, in part due to the limited functional data on TCR-epitope pairs, and the highly cross-reactive nature of TCR-epitope interactions. A design protocol like the one presented here, coupled with experimental validation, offers a promising path forward for future studies. By iteratively generating candidate peptide libraries, testing them experimentally, and refining the model with each iteration, it should be possible to more efficiently explore the high dimensional antigenic sequence space. This active learning strategy has a potential for revealing the “shape space” of TCR-epitope interactions, with broad applicabilities for early disease diagnosis, and for design of targeted immunotherapies. Moreover, this approach can help mitigate immunotherapy-related toxicity by identifying and excluding potential self-reactivities– beyond single-point mutations in the target epitope–such as the off-target Titin reactivity observed in an engineered T-cell therapy against MAGE-A3 that led to a severe cardiac toxicity [59].

Although our findings highlight the promise of structure-based models for immune recognition, a key limitation remains the scarcity of high-quality TCR-pMHC structural templates, despite advances from tools like AlphaFold3 [30] and TCRDock [18]. On the other hand, while TCR sequences are more widely available, sequence-based models often lack generalizability because existing paired TCR-pMHC data are skewed toward just a few peptides [12, 13]. A promising direction for future work lies in developing multi-modal flexi-ble methods that integrate structural and sequence-based models, leveraging both the depth and generalizability offered by structural information and the scalability of sequence data.

TCRs can cross-react with diverse peptides in multiple orientations and conformation [65]. To reflect this flexibility, we sampled peptide conformational space (i) implicitly, by applying noise to the coordinates during training, and (ii) explicitly, by locally relaxing the atomic environment after each amino acid substitution. Even so, AlphaFold3 predicts that our experimentally validated immunogenic designed peptides retain similar structural poses as their wild-type templates. This structural convergence may partly stem from AlphaFold3’s bias toward structures represented in its training set, but it also exposes limitations in our current sampling strategy. Future work should therefore aim to co-design peptide sequence, peptide backbone geometry, together with the CDR loop conformations of the interacting TCR, enabling a more comprehensive exploration of the peptide-TCR conformational landscape.

Although our study primarily examines MHC class I-restricted peptides, the broad applicability of structure-based modeling suggests that similar methods could be adapted for the more complex TCR-peptide-MHCII interactions, thereby offering new insights into CD4+ T-cell responses. Moreover, incorporating additional immunological factors—such as TCR clustering, co-receptor interactions, cross-reactivity with self-antigens, and functional immune profiling—can further enhance the accuracy of T-cell response predictions and improve the resulting de novo peptide desig pipeline.

## DATA AND CODE AVAILABILITY

Data and code for all the analyses can be found in the Github repository: https://github.com/StatPhysBio/tcr_antigen_design. Code for the original HERMES [27] model can be found in the Github repository: https://github.com/StatPhysBio/hermes.

## ACKNOWLEDGMENT

We thank Mikhail V. Pogorelyy for valuable comments and help with CellCyte. This work has been supported by the National Institutes of Health MIRA awards R35 GM142795 (AN, GV, MNP) and R35 GM141457 (PB), the CAREER award from the National Science Foundation grant 2045054 (AN), the National Institutes of Health R01 award AI136514 (PT and PB), the Royalty Research Fund from the University of Washington no. A153352 (AN, MNP), the Allen School Computer Science & Engineering Research Fellowship from the Paul G. Allen School of Computer Science & Engineering at the University of Washington (GV), the Microsoft Azure award from the eScience institute at the University of Washington (AN, GV, MNP), the TIRTL Bluesky Initiative at St. Jude Children’s Research Hospital (PT), and the ALSAC at St. Jude Children’s Research Hospital (PT and AAM). This work is also supported, in part, through the Departments of Physics and Computer Science and Engineering, and the College of Arts and Sciences at the University of Washington (AN, GV, MNP). This work was performed in part during the 2023 summer workshop “Statistical Physics and Adaptive Immunity” at the Aspen Center for Physics, which is supported by National Science Foundation grant PHY2210452. This work also benefited from discussions during the 2024 program “Interactions and Co-evolution between Viruses and Immune Systems” at the Kavli Institute for theoretical physics (KITP), which is supported by National Science Foundation grants PHY-2309135, and PHY-2309135, and the Gordon and Betty Moore Foundation Grant No. 2919.02.

## COMPETING INTEREST STATEMENT

PT is on the Scientific Advisory Board of Immunoscape and Shennon Bio, has received research support and personal fees from Elevate Bio, and consulted for 10X Genomics, Illumina, Pfizer, Cytoagents, Sanofi, Merck, and JNJ.

## Notes

### Summary of Updates

Figures 3-4 updated. Added supplementary figures S13-S17, modified supplementary figures S1,S4, S5, S8, S11. Reordered supplementary figures. Expanded main and supplementary texts.

## References

[1] M. Attaf, E. Huseby, and A. K. Sewell, Cell. Mol. Immunol. 12, 391 (2015).

[2] B. Liu, N. F. Greenwood, J. E. Bonzanini, A. Motmaen, J. Meyerberg, T. Dao, X. Xiang, R. Ault, J. Sharp,C. Wang, G. M. Visani, D. K. Vafeados, N. Roullier, A. Nourmohammad, D. A. Scheinberg, K. C. Garcia, and D. Baker, Science 389, 386 (2025).

[3] R. Leidner, N. Sanjuan Silva, H. Huang, D. Sprott, C. Zheng, Y.-P. Shih, A. Leung, R. Payne, K. Sutcliffe, J. Cramer, S. A. Rosenberg, B. A. Fox, W. J. Urba, and E. Tran, N. Engl. J. Med. 386, 2112 (2022).

[4] J. M. Mirazee, A. Aranganathan, S. Achar, D. Jia, X. Chen, B. P. Vani, C. D. Chien, M. Pouzolles, K. DeDe, P. Youkharibache, K. Walters, P. Tiwary, G. Altan-Bonnet, and N. Taylor, Biophys. J. 123, 552a (2024).

[5] E. H.-C. Hsiue, K. M. Wright, J. Douglass, M. S. Hwang,B. J. Mog, A. H. Pearlman, S. Paul, S. R. DiNapoli, M. F. Konig, Q. Wang, A. Schaefer, M. S. Miller, A. D. Skora, P. A. Azurmendi, M. B. Murphy, Q. Liu, E. Watson, Y. Li, D. M. Pardoll, C. Bettegowda, N. Papadopoulos, K. W. Kinzler, B. Vogelstein, S. B. Gabelli, and S. Zhou, Science 371, eabc8697 (2021).

[6] U. Sahin and Ö. Türeci, Science 359, 1355 (2018).

[7] E. Blass and P. A. Ott, Nat. Rev. Clin. Oncol. 18, 215 (2021).

[8] L. A. Rojas, Z. Sethna, K. C. Soares, C. Olcese, N. Pang, E. Patterson, J. Lihm, N. Ceglia, P. Guasp, A. Chu, R. Yu, A. K. Chandra, T. Waters, J. Ruan, M. Amisaki, A. Zebboudj, Z. Odgerel, G. Payne, E. Derhovanessian, F. Müller, I. Rhee, M. Yadav, A. Dobrin, M. Sadelain, M. Luksza, N. Cohen, L. Tang, O. Basturk, M. Gönen, S. Katz, R. K. Do, A. S. Epstein, P. Momtaz, W. Park, R. Sugarman, A. M. Varghese, E. Won, A. Desai, A. C. Wei, M. I. D’Angelica, T. P. Kingham, I. Mellman, T. Merghoub, J. D. Wolchok, U. Sahin, O. Türeci, B. D. Greenbaum, W. R. Jarnagin, J. Drebin, E. M. O’Reilly, and V. P. Balachandran, Nature 618, 144–150 (2023).

[9] S.-Q. Zhang, K.-Y. Ma, A. A. Schonnesen, M. Zhang, C. He, E. Sun, C. M. Williams, W. Jia, and N. Jiang, Nat. Biotechnol. 36, 1156 (2018).

[10] S. Nolan, M. Vignali, M. Klinger, J. N. Dines, I. M. Kaplan, E. Svejnoha, T. Craft, K. Boland, M. W. Pesesky, R. M. Gittelman, T. M. Snyder, C. J. Gooley, S. Semprini, C. Cerchione, F. Nicolini, M. Mazza, O. M. Delmonte, K. Dobbs, G. Carrenõ Tarragona, S. Barrio, V. Sambri, G. Martinelli, J. D. Goldman, J. Heath, L. D. Notarangelo, J. Martinez-Lopez, B. Howie, J. M. Carlson, and H. S. Robins, Front. Immunol. 16, 1488851 (2025).

[11] P. Dash, A. J. Fiore-Gartland, T. Hertz, G. C. Wang, S. Sharma, A. Souquette, J. C. Crawford, E. B. Clemens, T. H. O. Nguyen, K. Kedzierska, N. L. La Gruta, P. Bradley, and P. G. Thomas, Nature 547, 89 (2017).

[12] R. Vita, S. Mahajan, J. A. Overton, S. K. Dhanda, S. Martini, J. R. Cantrell, D. K. Wheeler, A. Sette, and B. Peters, Nucleic Acids Res. 47, D339 (2019).

[13] M. Shugay, D. V. Bagaev, I. V. Zvyagin, R. M. Vroomans, J. C. Crawford, G. Dolton, E. A. Komech, A. L. Sycheva, A. E. Koneva, E. S. Egorov, A. V. Eliseev, E. Van Dyk, P. Dash, M. Attaf, C. Rius, K. Ladell, J. E. McLaren, K. K. Matthews, E. B. Clemens, D. C. Douek, F. Luciani, D. van Baarle, K. Kedzierska, C. Kesmir, P. G. Thomas, D. A. Price, A. K. Sewell, and D. M. Chudakov, Nucleic Acids Res. 46, D419 (2018).

[14] G. Isacchini, A. M. Walczak, T. Mora, and A. Nourmohammad, Proc. Natl. Acad. Sci. U. S. A. 118, e2023141118 (2021).

[15] A. Weber, J. Born, and M. Rodriguez Martínez, Bioinformatics 37, i237 (2021).

[16] B. Meynard-Piganeau, C. Feinauer, M. Weigt, A. M. Walczak, and T. Mora, Proc. Natl. Acad. Sci. U. S. A. 121, e2316401121 (2024).

[17] K. H. Piepenbrink, B. E. Gloor, K. M. Armstrong, and B. M. Baker, Methods Enzymol. 466, 359 (2009).

[18] P. Bradley, Elife 12, e82813 (2023).

[19] R. Yin, H. V. Ribeiro-Filho, V. Lin, R. Gowthaman, M. Cheung, and B. G. Pierce, Nucleic Acids Res. 51, 569 (2023).

[20] A. Rives, J. Meier, T. Sercu, S. Goyal, Z. Lin, J. Liu, D. Guo, M. Ott, C. L. Zitnick, J. Ma, and R. Fergus, Proc. Natl. Acad. Sci. U. S. A. 118, e2016239118 (2021).

[21] Z. Lin, H. Akin, R. Rao, B. Hie, Z. Zhu, W. Lu, N. Smetanin, R. Verkuil, O. Kabeli, Y. Shmueli, A. Dos Santos Costa, M. Fazel-Zarandi, T. Sercu, S. Candido, and A. Rives, Science 379, 1123 (2023).

[22] A. Madani, B. Krause, E. R. Greene, S. Subramanian, B. P. Mohr, J. M. Holton, J. L. Olmos, Jr, C. Xiong, Z. Z. Sun, R. Socher, J. S. Fraser, and N. Naik, Nat. Biotechnol. 41, 1099–1106 (2023).

[23] R. Schmirler, M. Heinzinger, and B. Rost, Nat. Commun. 15, 7407 (2024).

[24] Y. Wang, H. Lv, Q. W. Teo, R. Lei, A. B. Gopal, W. O. Ouyang, Y.-H. Yeung, T. J. C. Tan, D. Choi, I. R. Shen, X. Chen, C. S. Graham, and N. C. Wu, Immunity 57, 2453 (2024).

[25] B. L. Hie, V. R. Shanker, D. Xu, T. U. J. Bruun, P. A. Weidenbacher, S. Tang, W. Wu, J. E. Pak, and P. S. Kim, Nat. Biotechnol. 42, 275 (2024).

[26] B. Bravi, A. Di Gioacchino, J. Fernandez-de Cossio-Diaz, A. M. Walczak, T. Mora, S. Cocco, and R. Monasson, Elife 12, e85126 (2023).

[27] G. M. Visani, M. N. Pun, W. Galvin, E. Daniel, K. Borisiak, U. Wagura, and A. Nourmohammad, bioRxiv2024.07.09.602403(2024), 10.1101/2024.07.09.602403.

[28] M. N. Pun, A. Ivanov, Q. Bellamy, Z. Montague, C. LaMont, P. Bradley, J. Otwinowski, and A. Nourmohammad, Proc. Natl. Acad. Sci. U. S. A. 121, e2300838121 (2024), https://www.pnas.org/doi/pdf/10.1073/pnas.2300838121.

[29] S. Chaudhury, S. Lyskov, and J. J. Gray, Bioinformatics26, 689 (2010).

[30] J. Abramson, J. Adler, J. Dunger, R. Evans, T. Green,A. Pritzel, O. Ronneberger, L. Willmore, A. J. Ballard, J. Bambrick, S. W. Bodenstein, D. A. Evans, C.-C. Hung, M. O’Neill, D. Reiman, K. Tunyasuvunakool, Z. Wu, A. Zemgulytė, E. Arvaniti, C. Beattie, O. Bertolli, A. Bridgland, A. Cherepanov, M. Congreve, A. I. Cowen-Rivers, A. Cowie, M. Figurnov, F. B. Fuchs, H. Gladman, R. Jain, Y. A. Khan, C. M. R. Low, K. Perlin, A. Potapenko, P. Savy, S. Singh, A. Stecula, A. Thillaisundaram, C. Tong, S. Yakneen, E. D. Zhong, M. Zielinski, A. Zídek, V. Bapst, P. Kohli, M. Jaderberg, D. Hassabis, and J. M. Jumper, Nature 630, 493 (2024).

[31] J. Dauparas, I. Anishchenko, N. Bennett, H. Bai, R. J. Ragotte, L. F. Milles, B. I. M. Wicky, A. Courbet, R. J. de Haas, N. Bethel, P. J. Y. Leung, T. F. Huddy, S. Pellock, D. Tischer, F. Chan, B. Koepnick, H. Nguyen, A. Kang, B. Sankaran, A. K. Bera, N. P. King, and D. Baker, Science 378, 49 (2022).

[32] L. M. Blaabjerg, M. M. Kassem, L. L. Good, N. Jonsson, M. Cagiada, K. E. Johansson, W. Boomsma, A. Stein, and K. Lindorff-Larsen, Elife 12, e82593 (2023).

[33] K. Michalewicz, M. Barahona, and B. Bravi, Structure32, 2422 (2024).

[34] D. J. Diaz, C. Gong, J. Ouyang-Zhang, J. M. Loy, J. Wells, D. Yang, A. D. Ellington, A. G. Dimakis, and A. R. Klivans, Nat. Commun. 15, 6170 (2024).

[35] G. Altan-Bonnet and R. N. Germain, PLoS Biol. 3, e356 (2005).

[36] S. R. Achar, F. X. P. Bourassa, T. J. Rademaker, A. Lee, T. Kondo, E. Salazar-Cavazos, J. S. Davies, N. Taylor, P. François, and G. Altan-Bonnet, Science 376, 880 (2022).

[37] J. D. Stone, A. S. Chervin, and D. M. Kranz, Immunology 126, 165 (2009).

[38] J.-L. Chen, G. Stewart-Jones, G. Bossi, N. M. Lissin, L. Wooldridge, E. M. L. Choi, G. Held, P. R. Dunbar, R. M. Esnouf, M. Sami, J. M. Boulter, P. Rizkallah, C. Renner, A. Sewell, P. A. van der Merwe, B. K. Jakobsen, G. Griffiths, E. Y. Jones, and V. Cerundolo, J. Exp. Med. 201, 1243 (2005).

[39] D. N. Garboczi, P. Ghosh, U. Utz, Q. R. Fan, W. E. Biddison, and D. C. Wiley, Nature 384, 134 (1996).

[40] S. Henikoff and J. G. Henikoff, Proc. Natl. Acad. Sci. U. S. A. 89, 10915 (1992).

[41] E. Fast, M. Dhar, and B. Chen, bioRxiv, 2023.09.12.557285 (2023).

[42] C. Hsu, R. Verkuil, J. Liu, Z. Lin, B. Hie, T. Sercu, A. Lerer, and A. Rives, in Proceedings of the 39th International Conference on Machine Learning, Proceedings of Machine Learning Research, Vol. 162, edited by K. Chaudhuri, S. Jegelka, L. Song, C. Szepesvari, G. Niu, and S. Sabato (PMLR, 2022) pp. 8946–8970.

[43] M. M. Davis, J. J. Boniface, Z. Reich, D. Lyons, J. Hampl, B. Arden, and Y. Chien, Annu. Rev. Immunol.16, 523 (1998).

[44] S. Zhong, K. Malecek, L. A. Johnson, Z. Yu, E. Vega-Saenz de Miera, F. Darvishian, K. McGary, K. Huang,J. Boyer, E. Corse, Y. Shao, S. A. Rosenberg, N. P. Restifo, I. Osman, and M. Krogsgaard, Proc. Natl. Acad. Sci. U. S. A. 110, 6973 (2013).

[45] C. M. Soto, J. D. Stone, A. S. Chervin, B. Engels,H. Schreiber, E. J. Roy, and D. M. Kranz, Cancer Immunol. Immunother. 62, 359 (2013).

[46] Y. Li, R. Moysey, P. E. Molloy, A.-L. Vuidepot, T. Mahon, E. Baston, S. Dunn, N. Liddy, J. Jacob, B. K. Jakobsen, and J. M. Boulter, Nat. Biotechnol. 23, 349 (2005).

[47] S. M. Dunn, P. J. Rizkallah, E. Baston, T. Mahon, B. Cameron, R. Moysey, F. Gao, M. Sami, J. Boulter, Y. Li, and B. K. Jakobsen, Protein Sci. 15, 710 (2006).

[48] M. Aleksic, O. Dushek, H. Zhang, E. Shenderov, J.-L. Chen, V. Cerundolo, D. Coombs, and P. A. van der Merwe, Immunity 32, 163 (2010).

[49] J. Pettmann, A. Huhn, E. Abu Shah, M. A. Kutuzov, D. B. Wilson, M. L. Dustin, S. J. Davis, P. A. van der Merwe, and O. Dushek, Elife 10, e67092 (2021).

[50] A. Esfandiary and S. Ghafouri-Fard, Immunotherapy 7, 411 (2015).

[51] J. L. Chen, P. R. Dunbar, U. Gileadi, E. Jäger, S. Gnjatic, Y. Nagata, E. Stockert, D. L. Panicali, Y. T. Chen, A. Knuth, L. J. Old, and V. Cerundolo, J. Immunol.165, 948 (2000).

[52] M. L uksza, Z. M. Sethna, L. A. Rojas, J. Lihm, B. Bravi, Y. Elhanati, K. Soares, M. Amisaki, A. Dobrin, D. Hoyos, P. Guasp, A. Zebboudj, R. Yu, A. K. Chandra, T. Waters, Z. Odgerel, J. Leung, R. Kappagantula, A. Makohon-Moore, A. Johns, A. Gill, M. Gigoux, J. Wolchok, T. Merghoub, M. Sadelain, E. Patterson, R. Monasson, T. Mora, A. M. Walczak, S. Cocco, C. Iacobuzio-Donahue, B. D. Greenbaum, and V. P. Balachandran, Nature 606, 389 (2022).

[53] M. F. Jensen and M. Nielsen, eLife 12, RP93934 (2024).

[54] X. Yang, M. Gao, G. Chen, B. G. Pierce, J. Lu, N.-P. Weng, and R. A. Mariuzza, J. Biol. Chem. 290, 29106 (2015).

[55] S. Gras, Z. Chen, J. J. Miles, Y. C. Liu, M. J. Bell, L. C. Sullivan, L. Kjer-Nielsen, R. M. Brennan, J. M. Burrows, M. A. Neller, R. Khanna, A. W. Purcell, A. G. Brooks, J. McCluskey, J. Rossjohn, and S. R. Burrows, J. Exp. Med. 207, 1555 (2010).

[56] Y. C. Liu, Z. Chen, M. A. Neller, J. J. Miles, A. W. Purcell, J. McCluskey, S. R. Burrows, J. Rossjohn, and S. Gras, J. Biol. Chem. 289, 16688 (2014).

[57] M. C. C. Raman, P. J. Rizkallah, R. Simmons, Z. Donnellan, J. Dukes, G. Bossi, G. S. Le Provost, P. Todorov, E. Baston, E. Hickman, T. Mahon, N. Hassan, A. Vuidepot, M. Sami, D. K. Cole, and B. K. Jakobsen, Sci. Rep. 6, 18851 (2016).

[58] D. M. Tadros, S. Eggenschwiler, J. Racle, and D. Gfeller, Nucleic Acids Res. 51, D428 (2023).

[59] G. P. Linette, E. A. Stadtmauer, M. V. Maus, A. P. Rapoport, B. L. Levine, L. Emery, L. Litzky, A. Bagg, B. M. Carreno, P. J. Cimino, G. K. Binder-Scholl, D. P. Smethurst, A. B. Gerry, N. J. Pumphrey, A. D. Bennett, J. E. Brewer, J. Dukes, J. Harper, H. K. Tayton-Martin, B. K. Jakobsen, N. J. Hassan, M. Kalos, and C. H. June, Blood 122, 863 (2013).

[60] B. J. Cameron, A. B. Gerry, J. Dukes, J. V. Harper, V. Kannan, F. C. Bianchi, F. Grand, J. E. Brewer, M. Gupta, G. Plesa, G. Bossi, A. Vuidepot, A. S. Powlesland, A. Legg, K. J. Adams, A. D. Bennett, N. J. Pumphrey, D. D. Williams, G. Binder-Scholl, I. Kulikovskaya, B. L. Levine, J. L. Riley, A. Varela-Rohena, E. A. Stadtmauer, A. P. Rapoport, G. P. Linette, C. H. June, N. J. Hassan, M. Kalos, and B. K. Jakobsen, Sci. Transl. Med. 5, 197ra103 (2013).

[61] C. Lundegaard, K. Lamberth, M. Harndahl, S. Buus, O. Lund, and M. Nielsen, Nucleic Acids Res. 36, W509 (2008).

[62] C. Lundegaard, O. Lund, and M. Nielsen, Bioinformatics24, 1397 (2008).

[63] M. Andreatta and M. Nielsen, Bioinformatics 32, 511 (2016).

[64] J. P. Roney and S. Ovchinnikov, Phys. Rev. Lett. 129, 238101 (2022).

[65] T. P. Riley, L. M. Hellman, M. H. Gee, J. L. Mendoza, J. A. Alonso, K. C. Foley, M. I. Nishimura, C. W. Vander Kooi, K. C. Garcia, and B. M. Baker, Nat. Chem. Biol. 14, 934 (2018).

[66] A. I. Webb, M. A. Dunstone, W. Chen, M.-I. Aguilar,Q. Chen, H. Jackson, L. Chang, L. Kjer-Nielsen, T. Beddoe, J. McCluskey, J. Rossjohn, and A. W. Purcell, J. Biol. Chem. 279, 23438 (2004).

[67] L. Wooldridge, J. Ekeruche-Makinde, H. A. van den Berg, A. Skowera, J. J. Miles, M. P. Tan, G. Dolton,M. Clement, S. Llewellyn-Lacey, D. A. Price, M. Peakman, and A. K. Sewell, J. Biol. Chem. 287, 1168 (2012).

[68] A. K. Sewell, Nat. Rev. Immunol. 12, 669 (2012).

[69] M. Vormehr, Türeci, Ö, and U. Sahin, Annu. Rev. Med.70, 395 (2019).

[70] K. Song and S. H. Pun, BME Front. 5, 0038 (2024).

