## Supplementary Text, figures, tables for "T-cell receptor specificity landscape revealed through de novo peptide design"

**Data and code availability.** Data and code for all the analyses can be found in the Github repository: https://github.com/StatPhysBio/tcr_antigen_design. Code for the original HERMES [1] model can be found in the Github repository: https://github.com/StatPhysBio/hermes.

#### I. HERMES NEURAL NETWORK MODEL FOR AMINO ACID PROPENSITIES

HERMES is a self-supervised structure-based machine learning model for proteins [1, 2]. HERMES has a 3D rotationally equivariant architecture and is trained to predict a residue's amino acid identity from its surrounding atomic environment within a 10 Å of the central residue's C-$\alpha$ in the 3D structure. The input to the neural network is a point cloud of atoms from a given structural neighborhood, in which all atoms associated with the central residue are masked. These point clouds are featurized by atom types, including computationally-added hydrogens, partial charge, and Solvent Accessible Surface Area (SASA). The point clouds are first projected onto the orthonormal Zernike Fourier basis, centered at the position of the (masked) central residue's C-$\alpha$ (Fig. 1A). The resulting *holographic* projections are fed to a stack of SO(3)-equivariant layers, the output of which is SO(3)-invariant, and is then passed through a multi-layer perceptron (MLP) to generate the desired predictions. The MLP's outputs are termed pseudo-energies, associated with the 20 different possible amino acids that can be placed at the masked residue, and act as logits within a softmax function to compute probabilities of each amino acid.

Each HERMES model is an ensemble of 10 individually-trained architectures. In this work, we used HERMES models trained on structural data with different levels of added noise: 0.00 noise (the original PDB structure), and 0.50 noise, which is obtained by adding Gaussian noise to the 3D coordinates of the original structure, with standard deviation of 0.50 Å. We only use models that were trained on the wild-type amino acid classification task [1]. Models were pre-trained on structures from 30% similarity split of ProteinNet's CASP12 set [3], resulting in around ∼10k training structures.

We refer the reader to refs. [1, 2] for further details on the architecture, training procedure, training data preparation, and the mathematical introduction to SO(3)-equivariant models in Fourier space.

#### II. TCR-PMHC BINDING AFFINITY AND T-CELL ACTIVITY PREDICTION

**Peptide energy calculation with HERMES.** To predict the affinity of TCR-pMHC complexes or the activity of TCRs induced by their interactions with a pMHC complex, we introduce the predicted energy for a peptide $\sigma$, given its surrounding TCR and MHC,

$$E_{\text{peptide}}(\sigma; \text{TCR, MHC}) = \sum_{i=1}^{\ell} E(\sigma_i; \text{TCR, MHC}, \sigma_{/i}). \tag{S1}$$

We assume that each peptide residue contributes linearly to the total energy by the amount $E(\sigma_i; \text{TCR, MHC}, \sigma_{/i})$, where $\sigma_i$ is the amino acid at position $i$ of the peptide. A residue's energy contribution $E$ is evaluated as its associated logit value from HERMES, taking in as input the atomic composition of the surrounding TCR, MHC and the rest of the peptide $\sigma_{/i}$ (Fig. 1A).

To compute peptide energies, HERMES operates on the structural composition of the protein complex. However, data on the impact of substitutions on the TCR-pMHC structure is limited, and computational models often fail to capture subtle conformational changes in the structure due to amino acid substitutions in a peptide. To address this issue, we adopt two protocols to estimate energy for a diverse array of peptides for a given TCR-MHC complex:

- HERMES-*fixed*: In the simplest approach, we select the TCR-pMHC structure with matching TCR and MHC to our query and with the peptide sequence that most closely matches the peptide of interest and directly input it into HERMES. The peptide energy is subsequently calculated as described in eq. S1 (Fig. 1B). This method does not modify the underlying structure, effectively assuming that any amino acid substitutions do not significantly alter the protein conformation.

- HERMES-*relaxed*: Since amino acid substitutions can locally reshape the protein complex, we introduce a more involved protocol to account for these structural changes. We begin with the available crystal structure of the TCR-pMHC complex that closely matches our query as template. We then mutate the original peptide to the desired state and relax the structure using PyRosetta's cartesian_ddg fastrelax protocol [4]. During relaxation, side chain and backbone atoms of the peptide are allowed to move, while only the side chain atoms of the MHC and TCR chains within a 10Å of the peptide are flexible. We then calculate the peptide energy with HERMES (eq.S1), using the relaxed structure of the mutated variant as input (Fig. 1B). Since PyRosetta's relaxation procedure is stochastic in nature, we run 100 realizations of the relaxation procedure and average the peptide energies across them. For ablation purposes, we also use an alternative protocol, whereby we use the HERMES peptide energy from the conformation with the minimum (most favorable) Rosetta energy computed on the relaxed atoms; we call this protocol *relaxed-min_energy*.

Overall, we use four protocols for scoring peptides that differ in their restrained protocol (HERMES-*fixed* or HERMES-*relaxed*), and the amount of noise injected during training of the underlying HERMES model. Specifically, we train HERMES models with no added noise ($\sigma = 0.00$ $\AA$) and another with moderate Gaussian noise with standard deviation $\sigma = 0.50$ $\AA$ applied to the atomic coordinates of the protein structure data during training. We refer to the resulting peptide scoring protocols as HERMES-*fixed* 0.00, HERMES-*fixed* 0.50, HERMES-*relaxed* 0.00, HERMES-*relaxed* 0.50, specifying both the model and the noise amplitude.

**Evaluation of $\mathbf{EC_{50}}$ from dose response measurements against peptide libraries.** Ref. [5] presents dose response measurements for fraction of 4-1BB$^+$ CD8$^+$ T-cells across varying peptide concentrations. These measurements are done for peptide libraries of single point mutants from a given wild-type peptide, across 7 different TCR-MHC systems: three TCRs recognizing variants of the highly immunogenic HLA-A02:01-restricted human cytomegalovirus (CMV) epitope NLVPMVATV (NLV); three TCRs recognizing variants of the weaker HLA-A02:01-restricted melanoma gp100 self-antigen IMDQVPFSV; and one TCR recognizing variants of the weakly immunogenic HLA-B*27:05-restricted pancreatic cancer neo-peptide GRLKALCQR [5].

The half-maximal effective concentration ($EC_{50}$) from the dose response curves of 4-1BB$^+$ CD8$^+$ T-cell fractions can quantify the level of T-cell activity in response to each peptide. We used our own fits to estimate the $EC_{50}$ associated with T-cell responses. Specifically, we fitted a Hill function with background to the dose response measurements for each peptide from ref. [5],

$$f(x) = \frac{(A - A_0)x^n}{x^n + K^n} + A_0 \tag{S2}$$

where $f(x)$ is the fraction of 4-1BB$^+$ CD8$^+$ T-cells pulsed by a peptide at concentration $x$, $A$ is the response amplitude, $A_0$ is the background activation, $n$ is the Hill coefficient, and $K$ is the half-maximal effective concentration $EC_{50}$. Similar to ref. [5], we regularized our fitted Hill functions, whereby the deviation of the amplitude $A$ and the Hill coefficient $n$ from 1, and the background activation $A_0$ from 0 are penalized. The code to fit these curves is available in our github repository.

### III. DE-NOVO PEPTIDE DESIGN

**Peptide design algorithm.** Given a TCR and an MHC, we use HERMES to design de novo peptides that can form a stable TCR-pMHC complex, and potentially, elicit a T-cell response. Our approach starts with an existing template TCR-pMHC structure. We refer to the peptide in the template as wild-type; see Table S5 for the templates we use in our analyses. Similar to evaluating the peptide energy, we use two pipelines to sample novel peptides:

- HERMES-*fixed*: This is our basic pipeline, in which we generate novel peptide sequences by conditioning on the fixed TCR-pMHC template structure. Specifically, we sequentially mask the atoms of each amino acid along the peptide and then use HERMES to compute the probability $P(\sigma_i|\mathbf{x}_i)$ of different amino acid types $\sigma_i$ at the masked residue $i$, conditioned on the surrounding atomic neighborhood $\mathbf{x}_i$, within a 10 Å radius of the masked residue's C-$\alpha$. The neighborhood $\mathbf{x}_i$ can include atoms from the TCR, the MHC, and the other amino acids in the peptide. Repeating this procedure for every residue along the peptide allows us to construct a Position Weight Matrix (PWM) that represents the probabilities of different amino acids at each peptide position, conditioned on the surrounding structural neighborhood. We generate peptides using multinomial sampling from the PWM; see the schematic in Fig. 1C.

- HERMES-*relaxed*: This pipeline incorporates simulated annealing and Markov chain Monte Carlo (MCMC) sampling to account for structural changes resulting from peptide substitutions. We start with the template TCR-MHC structure and an initial random peptide sequence $\sigma^{(0)}$, which is *in-silico* packed via PyRosetta, followed by the Relax protocol, yielding a relaxed structure $\mathbf{x}^{(0)}$. During relaxation, both side chain and backbone atoms of the peptide are allowed to move, while only the side chain atoms of the MHC and TCR within 10 Å of the peptide are flexible. Following the HERMES-*fixed* protocol, we then evaluate the PWM associated with this relaxed structure and use it to generate a new peptide sequence, $\sigma^{(1)}$. This sequence is again packed and relaxed, producing a new structure, $\mathbf{x}^{(1)}$. Using eq. S1, we evaluate the energies of the peptides in their respective (relaxed) structures as $E(\sigma^{(0)};\mathbf{x}^{(0)})$ and $E(\sigma^{(1)};\mathbf{x}^{(1)})$, and apply the Metropolis-Hastings criterion to accept the transition from $\sigma^{(0)}$ to $\sigma^{(1)}$ with probability:

$$P_{\sigma^{(0)}\to\sigma^{(1)}} = \min\left(1, e^{-\frac{E(\sigma^{(1)};\mathbf{x}^{(1)})-E(\sigma^{(0)};\mathbf{x}^{(0)})}{T}}\right) \tag{S1}$$

where $T$ is the temperature. This process is iterated, incrementally reducing $T$ at each step, until convergence. The peptide obtained at the final iteration represents the protocol's de novo design. Repeating this procedure many times produces a diverse set of peptides spanning various distances from the wild-type sequence; see the schematic in Fig. 1C and Algorithm S1 for more details, and the the convergence curves in Fig. S6C,F,I.

**Input:** TCR-pMHC structure $\mathbf{x}$, maximum number of iterations N;
**Output:** peptide sequence $\sigma^{(N)}$;
$T_0 \leftarrow 10$;
$T_f \leftarrow 0$;
$a \leftarrow 0.95$;
$T^{(0)} \leftarrow T_0$;
$\sigma^{(0)} \leftarrow$ random sequence;
$\mathbf{x}^{(0)} \leftarrow \text{RosettaMutate}(\mathbf{x}^{(0)}, \sigma^{(0)})$;
$\mathbf{x}^{(0)} \leftarrow \text{RosettaFastRelax}(\mathbf{x}^{(0)})$;
$E^{(0)}_{\text{peptide}} \leftarrow \sum_{i\in\mathcal{R}_{\text{peptide}}} E(\sigma^{(0)}_i; \mathbf{x}^{(0)}_i)$;
**for** $k = 1, 2, \ldots, N$ **do**
    **for** $i \in \mathcal{R}_{peptide}$ **do**
        $\sigma^{(\text{temp})}_i \sim \text{Multinomial}(\text{softmax}([E(\text{aa}; \mathbf{x}^{(k-1)}_i)/T^{(k)}]_{\text{aa}\in\text{amino-acids}}))$;
    **end**
    $\mathbf{x}^{(\text{temp})} \leftarrow \text{RosettaMutate}(\mathbf{x}^{(k-1)}, \sigma^{(\text{temp})})$;
    $\mathbf{x}^{(\text{temp})} \leftarrow \text{RosettaFastRelax}(\mathbf{x}^{(\text{temp})})$;
    $E^{(\text{temp})}_{\text{peptide}} \leftarrow \sum_{i\in\mathcal{R}_{\text{peptide}}} E(\sigma^{(\text{temp})}_i; \mathbf{x}^{(\text{temp})}_i)$;
    $p \sim \text{Uniform}(0, 1)$;
    **if** $p < \exp(-(E^{(temp)}_{peptide} - E^{(k-1)}_{peptide})/T^{(k)})$ **then**
        $E^{(k)}_{\text{peptide}} \leftarrow E^{(\text{temp})}_{\text{peptide}}$;
        $\mathbf{x}^{(k)} \leftarrow \mathbf{x}^{(\text{temp})}$;
        $\sigma^{(k)} \leftarrow \sigma^{(\text{temp})}$;
    **else**
        $E^{(k)}_{\text{peptide}} \leftarrow E^{(k-1)}_{\text{peptide}}$;
        $x^{(k)} \leftarrow \mathbf{x}^{(k-1)}$;
        $\sigma^{(k)} \leftarrow \sigma^{(k-1)}$;
    **end**
    $T^{(k+1)} \leftarrow (T_0 - T_f)a^k + T_f$;
**end**

**Algorithm S1:** HERMES-*relaxed* protocol for generating TCR-MHC-specific peptide sequences. For brevity, we define $\mathbf{x}_i \equiv (\text{TCR}, \text{MHC}, \sigma_{/i})$ and drop the arguments of $E_{\text{peptide}}$.

Similar to scoring, we use four protocols for peptide design that differ in their restrained protocol (HERMES-*fixed* or HERMES-*relaxed*), and the amount of noise injected during training of the underlying HERMES model. Specifically, we train HERMES models with no added noise ($\sigma = 0.00$ Å) and another with moderate Gaussian noise with standard deviation $\sigma = 0.50$ Å applied to the atomic coordinates of the protein structure data during training. We refer to the resulting peptide design protocols as HERMES-*fixed* 0.00, HERMES-*fixed* 0.50, HERMES-*relaxed* 0.00, HERMES-*relaxed* 0.50, specifying both the model and the noise amplitude.

**Filtering metrics for peptide design with HERMES and benchmarking of different design approaches.** For each system, we first selected designs predicted by NetMHCPan to be at least weak binders to their target MHCs (EL_rank $\leq 2.0$) [6]. We then used TCRdock [7] to filter peptides for reactivity. TCRdock is a structure prediction algorithm built on top of AlphaFold2 [8], and refined to model TCR-pMHC interactions. Its PAE score for the TCR-pMHC interface can be used to distinguish reactive from decoy peptides [7], and therefore, can provide a metric for filtering our designed peptides; other work has similarly used the PAE of structure-prediction algorithms to filter designed protein binders [9]. Using TCRdock, we retained those peptides whose PAE scores at the TCR-pMHC interface fell below a threshold, as illustrated in Figs. S13, S14, and S15). The distribution of PAE scores is highly system specific [9], so we set each system's threshold near the TCRdock PAE value of its *wild-type* peptides in complex with the respective TCR-MHC, as the wild types are known binders. Among the filtered peptides, we selected 93 for experimental validation, ensuring that designs from both HERMES-*fixed* and HERMES-*relaxed*, as well as from models with 0.0 and 0.5Å noise levels were represented. We also enforced a minimum difference of 2-3 amino acids (depending on the system) between the designs and the wild-type peptides.

It should be noted that the AF3 PAE score can also discriminate between reactive and decoy peptides (Figs. S8, S16), making it also suitable for filtering. However, a full-access implementation of AF3 was unavailable before our experimental validations. Therefore, we used AF3 only as a secondary filter for designs in the EBV and MAGE systems, and for benchmarking against TCRdock's PAE score for the designed peptides (Figs. S8); see details below.

**Design parameters for different TCR-pMHC systems.** Here, we provide the details on the specifics of peptide selection for each TCR-MHC system. It should be noted that as we progressed from NY-ESO to EBV and MAGE, we chose to explore the sequence space farther from the wild-type peptides, leading to more challenging designs.

- *NY-ESO:* The wild-type peptides (NYESO: SLLMWITQC and NYESO V9C: SLLMWITQV) have TCRdock PAEs of 5.17 and 5.18, respectively. As positive designs, we selected 58 peptides with the lowest TCRdock PAE values (all below 5.35) and a Hamming distance of at least two amino acids from their respective wild-type sequences. For negative designs, we uniformly sampled 35 peptides among those designed by HERMES whose TCRdock PAE values fell between 5.5 and 7.0. We did not enforce all HERMES models to be represented equally among these designs, nor did we enforce equal representation of the template structures. However, we note that all models are represented among positive designs (Fig. S9).

- *EBV:* The wild-type peptides (HPVG: HPVGEADYFEY and HPVG E5Q: HPVGQADYFEY) have TCRdock PAEs of 4.90 and 4.82, and AF3 PAEs of 1.15 and 1.13, respectively. We selected 93 positive peptides using a TCRdock's PAE score cutoff of 5.1 and AF3 PAE score cutoff of 2.0. Specifically, we sampled uniformly below these cutoffs with the following design protocols: HERMES-*fixed* at both noise levels using the HPVG structure (3MV7) as template (24 peptide), HERMES-*fixed* at both noise levels, using the HPVG E5Q structure (4PRP) as template (23 designs); HERMES-*relaxed* at both noise levels, using the HPVG structure (3MV7) as template (24 peptides), and HERMES-*relaxed* at both noise levels using the HPVG E5Q structure (4PRP) as template (23 designs); see Fig. S10.

- *MAGE:* The wild-type peptides (MAGE-A3: EVDPIGHLY and Titin: ESDPIVAQY) have TCRdock PAEs of 5.21 and 5.03, and AF3 PAEs of 1.00 and 0.98, respectively. We selected 93 positive peptides using a TCRdock's PAE score cutoff of 5.2; the AF3 PAE scores for all of these designed peptide are comparable to that of the wild-types. For these designs, we sampled uniformly below the PAE cutoffs with the following design protocols: HERMES-*fixed* at both noise levels using MAGE-A3 structure (5BRZ) as template (15 peptides); HERMES-*fixed* at both noise levels using the MAGE-A3 structure (5BRZ) as template with $E_1$ fixed (15 peptides); HERMES-*relaxed* at both noise levels using the MAGE-A3 structure (5BRZ) as template (16 peptides); HERMES-*relaxed* at bothsee Fig. S10 noise levels using the MAGE-A3 structure (5BRZ) as template with $E_1$ fixed (15 peptides); HERMES-*fixed* at both noise levels and using the Titin structure (5BS0) as template (16 peptides); HERMES-*relaxed* at both noise levels using the Titin structure (5BS0) as template (16 peptides); see Fig. S11.

**Characterizing the size of the sequence space for design.** The size of the sequence space to explore grows exponentially with increasing distance from the wild-type peptide. If no restriction is imposed (i.e., all amino acids are permissible with equal probabilities), the size of the sequence space follows,

$$\Omega_0(d) = \binom{\ell}{d} 19^d \tag{S2}$$

where $\ell$ is the length of the peptide, and $d$ is the distance from the wild-type sequence.

MHC presentation imposes constraints on amino acid usages in peptides, especially at the anchor residues. Let $p_i(a)$ denote the probability of amino acid $a$ at position $i$ of the peptide, e.g., set by the sequence-specific preferences of the MHC allele. The probability of introducing a mutation at position $i$ depends on the prominence of the wild-type amino acid, and is equal to $q_i = 1 - p_i(\text{wt})$. The typical number of non-wildtype amino acids realized at position $i$ can be estimated by $n_i = 2^{S_i}$, where $S_i = -\sum_{a \neq \text{wt}} \hat{p}_i(a) \log_2 \hat{p}_i(a)$ is the entropy associated with the non-wildtype amino acid frequencies, and $\hat{p}_i(a) = \frac{p_i(a)}{1-p_i(\text{wt})}$ is the normalized probability of amino acid $a$, obtained by excluding the wild-type from the set. Therefore, the typical size of the sequence space at Hamming distance $d$ from the wild-type peptide—subject to constraints imposed by the amino acid frequencies $p_i(a)$ (e.g., due to MHC restriction)—is given by,

$$\Omega(d) = \sum_{\{i_1,\dots,i_d\}} (q_{i_1} n_{i_1}) \dots (q_{i_d} n_{i_d}) \equiv \sum_{\{i_i,\dots,i_d\}} 2^{\hat{S}_{i_1} + \dots + \hat{S}_{i_d}} \tag{S3}$$

where the summation runs over all $\binom{\ell}{d}$ ways to choose $d$ distinct sites at which mutations may occur, and $q_i n_i$ is the expected number of substitutions associated with site $i$—evaluated as the likelihood of mutating site $i$ times the typical number of viable amino acid replacements at that site—and $\hat{S}_i = S_i + \log_2 q_i$ is the resulting adjusted entropy at site $i$.

In principle, evaluating $\Omega(d)$ in eq. S3 involves summation over a combinatorially large number of terms, which can become computationally prohibitive. In practice, however, many of the sites have the same adjusted entropy, which simplifies the evaluations of $\Omega(d)$. For example, the adjusted entropies for mutations away from the 9 amino acid NY-ESO peptide restricted by HLA-A*02:01 yield the following (rounded) values: $[4, 2, 4, 4, 4, 3, 4, 4, 2]$ in bits, indicating six sites with 4 bits of entropy (or $2^4 = 16$ typical substitutions), one site with 3 bits (8 typical substitutions), and two sites with 2 bits (4 typical substitutions). To demonstrate, in this case, the number of sequences at distance $d = 5$ from the wild-type follows,

$$\begin{aligned}\Omega_{\text{NYESO}}(d=5) &= \binom{6}{5}16^5 + \binom{6}{4}16^4 \times \left[\binom{2}{1}4^1 + 8^1\right] + \binom{6}{3}16^3 \times \left[\binom{2}{1}4^1 \times 8^1 + \binom{2}{2}4^2\right] + \binom{6}{2}16^2 \times \binom{2}{2}4^2 \times 8^1 \\ &= 29{,}065{,}216\end{aligned} \tag{S4}$$

Similar calculations can be done to characterize the size of the sequence space at varying distances from the wild-type, as shown in Fig. 4.

### IV. ALTERNATIVE COMPUTATIONAL METHODS FOR SCORING PEPTIDES AND DESIGN

We compare our approach to alternative methods (listed below), based on which we characterize peptide scores akin to HERMES's peptide energy, and when possible, use them to design novel peptides. A detailed comparison between the accuracy of different methods in predicting the affinity of TCR-pMHC interactions and the peptide-induced activation of T-cells is presented in Tables S2 and S3. Comparison between the fidelity of the designs–only according to computational metrics–is shown in Figures S13, S14, and S15. All the designed peptides as well as the code to generate samples are available in our github repository.

- *BLOSUM62 substitution matrix* [10]: The BLOSUM62 substitution matrix characterizes the favorability of substitutions between all pairs of amino acids, and has been used widely to characterize the mutational effects in proteins. As our first (and simplest) benchmark, we used BLOSUM62 to define a peptide energy (score). Specifically, we score a peptide by summing over the BLOSUM62 scores associated with the amino acid substitutions from the wild-type peptide $\sigma^{\text{wt}}$ (the one present in the template structure) to the query peptide sequence $\sigma$ that we are interested in,

$$E^{(\text{B62})}_{\text{peptide}}(\sigma; \sigma^{\text{wt}}) = \sum_{i=1}^{\ell} B62[\sigma_i^{\text{wt}} \to \sigma_i] \tag{S1}$$

Notably, the BLOSUM62 predicted peptide energy only depends on the substitutions in the peptide relative to the wild-type amino acid, and does not depend on the structural context of these substitutions (Fig. S2).

For designs with BLOSUM62, we independently sample amino acids along the peptide based on their favorability with respect to the wild-type amino acid at each position. Specifically, at a given position $i$ we interpret the substitution score $B62[\sigma_i^{\text{wt}} \to \sigma_i]$ from the wild-type to any of the other 19 amino acids as the energy difference between the two states, and by applying a softmax function with temperature $T$, we evaluate the probability of these possible substitutions. We then use multinomial sampling to draw new amino acids from this probability distribution at each position,

$$\sigma_i \sim \text{Multinomial}\left(\text{softmax}\left[\frac{B62[\sigma_i^{\text{wt}} \to \sigma]}{T}\right]_{\sigma \in \text{amino-acids}}\right) \tag{S2}$$

For our analyses, we use temperatures $T = 1.0, 2.0, 3.0$ to modulate the variability of the designed peptides.

- *Łuksza et al. $M$ and $d \cdot M$ substitution matrices* [5]: Here, $M$ is an amino acid substitution matrix, and $d$ is a vector of position-specific scale factors for 9-mer peptides. Briefly, they were constructed to fit the $\text{EC}_{50}$ measurements on seven saturation mutagenesis experiments of three different wild-type peptides, against a total of seven TCR-MHC pairs–the very same T-cell activity data that we used to test HERMES in Fig. 3. Specifically, $d$ and $M$ were constructed such that:

$$d_i \cdot M[\sigma_i^{\text{wt}} \to \sigma_i^{\text{mt}}] \approx \log \frac{\text{EC}_{50}^{\text{mt}}}{\text{EC}_{50}^{\text{wt}}} + R_{\text{B62}}[\sigma_i^{\text{wt}} \to \sigma_i^{\text{mt}}] \tag{S3}$$

where the mutant and wild-type peptides only differ at position $i$, and $R_{\text{B62}}$ is a regularization term reflecting the BLOSUM62 substitution matrix; we refer the reader to the Supplementary Methods of ref. [5] for details. Since all peptides considered when constructing $d$ and $M$ are 9-mers, the position scale factors can only be used on 9-mer peptides. We use $d$ and $M$ for scoring in a manner analogous to BLOSUM62:

$$E_{\text{peptide}}^{(d\cdot M)}(\sigma; \sigma^{\text{wt}}) = \sum_{i=1}^{\ell} d_i \cdot M[\sigma_i^{\text{wt}} \to \sigma_i] \tag{S4}$$

and

$$E_{\text{peptide}}^{(M)}(\sigma; \sigma^{\text{wt}}) = \sum_{i=1}^{\ell} M[\sigma_i^{\text{wt}} \to \sigma_i] \tag{S5}$$

Similar to BLOSUM62, the predicted peptide energies $E_{\text{peptide}}^{(d\cdot M)}$ and $E_{\text{peptide}}^{(M)}$ only depend on the type substitutions, and not on their structural contexts.

- *MHC position weight matrices* [11]: MHC alleles exhibit distinct sequence-specific preferences for peptide presentation, most notably in the peptide's amino acid composition at anchor positions. Computational models have characterized these preferences with position weight matrices (PWMs) that reflect the the probability of different amino acids at each position for peptides presented by a given MHC allele [11–14]. As a benchmark for our designs, we use the PWMs from the MHC Motif Atlas [11] to independently sample amino acids at each site along the peptide. The resulting peptide sequences should reflect biases imposed by MHC presentation but not interactions with TCRs.

- *ProteinMPNN* [15]: ProteinMPNN is a commonly used tool for inverse-folding (i.e., designing protein sequences consistent with the backbone of a structure). It is primarily used to sample amino acid sequences conditioned on a protein's backbone structure, and optionally with partial sequence information (i.e., amino acids specified at some of the residues). As ProteinMPNN also outputs probability distributions of amino acids at different sites of the protein, it can also be used to score peptides (i.e., evaluate peptide energy) in a manner similar to HERMES, as

$$E_{\text{pep 1-by-1}}^{(\text{MPNN})}(\sigma; TCR, MHC) = \sum_{i=1}^{\ell} \log P^{(\text{MPNN})}(\sigma_i; \text{backbone}, \sigma_{\text{TCR}}, \sigma_{\text{MHC}}, \sigma_{/i}) \tag{S6}$$

where $P^{(\mathrm{MPNN})}(\sigma_i; \mathrm{backbone}, \mathrm{TCR}, \mathrm{MHC}, \sigma_{/i})$ is the ProteinMPNN estimate for the probability of amino acid $\sigma_i$ at position $i$ of the peptide, given the backbone structure of the whole protein complex, and the amino acid sequence of the MHC $\sigma_{\mathrm{MHC}}$, TCR $\sigma_{\mathrm{TCR}}$, and the rest of the peptide $\sigma_{/i}$. We term this scoring scheme as peptide 1-by-1 since we mask the peptide's residues one at a time.

We explored two additional ProteinMPNN scoring schemes that differ in the residues they mask and score. As our second scoring scheme, we simultaneously mask the entire peptide's sequence $\sigma$, instead of one amino-acid at a time, and evaluate the score for the whole peptide $E^{(\mathrm{MPNN})}_{\mathrm{pep}}(\sigma; TCR, MHC)$:

$$E^{(\mathrm{MPNN})}_{\mathrm{pep}}(\sigma; TCR, MHC) = \sum_{i=1}^{\ell} \log P^{(\mathrm{MPNN})}(\sigma_i; \mathrm{backbone}, \sigma_{\mathrm{TCR}}, \sigma_{\mathrm{MHC}}) \tag{S7}$$

For the third scoring scheme, we consider masking and scoring the TCR residues that make contact with the peptide–defined as those whose C-$\alpha$ lie within 12 Å of any peptide residue's C-$\alpha$. We call this set **TCR-int** and its corresponding amino acid composition as $\sigma_{\text{TCR-int}}$. In this scheme, both the peptide and the interacting TCR residues are masked simultaneously and scored together:

$$\begin{aligned} E^{(\mathrm{MPNN})}_{\text{pep and tcr}}(\sigma; TCR, MHC) &= \sum_{i=1}^{\ell} \log P^{(\mathrm{MPNN})}(\sigma_i; \mathrm{backbone}, \sigma_{\text{TCR / TCR-int}}, \sigma_{\mathrm{MHC}}) \\ &+ \sum_{j=1}^{n} \log P^{(\mathrm{MPNN})}(\sigma_{\text{TCR-int}_j}; \mathrm{backbone}, \sigma_{\text{TCR / TCR-int}}, \sigma_{\mathrm{MHC}}) \end{aligned} \tag{S8}$$

Similar to HERMES, we considered ProteinMPNN models trained with two noise levels: 0.02 Å (very little noise) and 0.20 Å.

For designs with ProteinMPNN, we use the algorithm's native functionalities, keeping the TCR and MHC structures fixed (from the design template) and prompting the model to design only the peptide. We use two ProteinMPNN models (trained with 0.02 Å and 0.20 Å noise), and use a temperature of 0.7 for all systems.

- *ESM-IF1* [16]: ESM-IF1, like ProteinMPNN, is an inverse-folding model, but it lacks native support for scoring or designing proteins when only partial sequence information is available.

  To access the ESM-IF1 model, we used the code in the esm repository at `https://github.com/facebookresearch/esm/tree/main/examples/inverse_folding`.

  For scoring a peptide sequence given a TCR-MHC context, we used the script `score_log_likelihoods.py` with the `--multichain-backbone` flag, with the same TCR-pMHC structure inputs as for HERMES and ProteinMPNN. This script computes a score for the peptide sequence conditioned on the backbone coordinates of the entire TCR-pMHC complex, but *not* on the sequence of the TCR and the MHC:

$$E^{(\mathrm{ESM-IF1})}_{\mathrm{peptide}}(\sigma; TCR, MHC) = \log P^{(\mathrm{ESM-IF1})}_{\mathrm{peptide}}(\sigma; \mathrm{backbone})$$

  To design peptide sequences given a TCR-MHC context, we used the script `sample_sequences.py` with the `--multichain-backbone` flag, and a temperature of 0.7 for all systems, with the same TCR-pMHC structure inputs as for HERMES and ProteinMPNN. Similar to scoring, this script samples peptide sequences conditioned on the backbone coordinates of the entire TCR-pMHC complex, but *not* on the sequences of the TCR and MHC.

- *TCRdock* [7]: TCRdock is a structure prediction algorithm built on top of AlphaFold2 [8], and refined to predict the structure of TCR-pMHC complexes. The algorithm model the structure of TCR-pMHC complexes by leveraging structural templates provided by its curated database, as well as the knowledge of TCR-pMHC canonical binding poses. Crucially, TCRdock's Predicted Alignment Error (PAE) for TCR-pMHC interfaces was found to well discriminate between true peptide binders and random decoys [7]. Thus, to benchmark against TCRdock, we interpret this PAE score as peptide energy, and use it to predict TCR-pMHC binding affinities and peptide-induced T-cell activities.

  TCRdock takes as input sequences of a TCR, MHC, and peptide, and identifies TCR-pMHC structures in its database with similar amino acid sequences to the query to serve as templates to model the structure of the input TCR-pMHC complex. To examine how template-query sequence similarity influences TCRdock's predictive accuracy, we ran the algorithm under two conditions: the default mode, which uses any closest-matching templates available, and the *benchmark* mode, which restricts the maximum sequence similarity of templates with the input TCR-pMHC.

We use TCRdock to characterize peptide energies, and to filter the sequences designed by HERMES. However, TCRdock does not provide a natural way to sample peptides for design, and because of its high computational cost, we did not include TCRdock PAE calculations in our design pipeline (e.g., for MCMC sampling).

- *TULIP* [17]: TULIP is a transformer-based model trained to generate either TCR or peptide sequences conditioned on the other binding partners in a TCR-pMHC complex. The conditional likelihood function learned by TULIP can be used to score peptides within a TCR-pMHC context. Specifically, this scoring scheme was used in ref. [17] to predict the T-cell activity measurements from ref. [5], and we benchmarked our predictions against TULIP's for this data. In Table S3, we present the Spearman correlations between TULIP predictions and the fitted $EC_{50}$ values as reported in ref. [5], which are the same correlations reported in the TULIP paper [17]. Additionally, we provide correlations with the fitted $EC_{50}$ values obtained using the procedure outlined in Section II and eq. S2, consistent with correlations reported in Fig. 3.

- *NetTCR2.2* [18]: NetTCR2.2 consists of a Convolutional Neural Network classifier to predict TCR-peptide binding from the the amino acid sequences of the peptide and of all six CDRs from the $\alpha$ and the $\beta$ chains of a TCR. Because the training set contains many examples of different TCRs binding to the same peptide, but far fewer cases of multiple peptides tested against a given TCR, the model performs best when ranking candidate TCRs for a fixed peptide, as reported in ref. [18]. By contrast, we aim to address the complementary task for ranking peptides for a fixed TCR.

  We query NetTCR2.2 through the authors' web server (`https://services.healthtech.dtu.dk/services/NetTCR-2.2/`). CDR boundaries are assigned by aligning each TCR sequence to the IMGT reference sequences, using code from the TCRdock repository [7]. Of the TCR-pMHC pairs examined in this work, only one system (TCR1, Table S1) appears in the NetTCR2.2 training set, and even then with just a single annotated positive peptide. Therefore, this small overlap is unlikely to bias our predictions.

- *TAPIR* [19]: Similar to NetTCR-2.2, TAPIR consists of a Convolutional Neural Network trained to predict TCR-peptide-MHC binding. As input, TAPIR considers the CDR3's amino acid sequences, the V and J genes, the peptide's amino acid sequence, and the MHC allele. We use TAPIR for scoring via the web interface hosted by the authors `https://vcreate.io/tapir/`. We extract V/J genes and CDR3 from a TCR sequence by aligning it to the IMGT reference sequences, using code from the TCRdock repository [7]. We find that five out of nine of the TCR-pMHC systems in this study are present in the training data of TAPIR, but each only with a single annotated positive peptide (1G4 TCR, TCR1, TCR4, TCR5, TCR6, Table S1), and therefore, this small overlap is unlikely to bias our predictions.

**Computational modeling of TCR-pMHC templates with AlphaFold3.** For six out of seven systems with T-cell activity measurements from ref. [5] (TCRs 2-7), no experimentally-determined structure is available. For these, we generated structures using the AlphaFold3 (AF3) model [20]. Furthermore, to probe how sensitive HERMES is to AF3 modeled inputs, we produced AF3 predictions for every TCR-pMHC complex we studies in this work–both for scoring and design design targets–using the wild-type sequences of all the chains as inputs; see Dataset S1 for these sequences. Each target was folded using both the AF3 server's option with template search (with default cutoff date of September 9th 2021), and without template, the latter approximating the scenario in which no homologous structures exist. We note, however, that several of TCR-pMHC complexes we analyzed here were likely present in the AF3 training data, so the "template-free" predictions can only serve as an approximate test of true de-novo performance.

We fold TCR-pMHC systems for which we have a structure using restricted amino-acid sequences of the TCR and MHC chains that were extracted with functionalities in the TCRdock codebase, and are equivalent to the sequences that TCRdock would use to predict the structure [7]. Specifically, we first extract the amino-acid sequences from the experimental structure. Then, we extract V/J genes, CDR3 and MHC allele from them using 'parse_tcr_pmhc_pdbfile.py', and input them to 'setup_for_alphafold.py' to get the restricted amino-acid sequences. For the bispecific Fab antibody-mediated T-cell therapy from ref. [21], we used AF3 to fold and dock the Fab and the pMHC by extracting the amino-acid sequences from the experimental structure. We found that AF3 displayed high uncertainty (as measured by plDDt) only at the Fab-pMHC interface, and not in the folds of individual chains. The sequences we used as input to AF3 are listed in Dataset S1.

**Benchmarking HERMES predictions for T-cell affinity and activity.** We compared the performance of HERMES models at predicting the binding affinity and activity of T-cells against substitution matrices (BLOSUM62 [10], and Łuksza *et al*'s TCR-specific $d \cdot M$ and $M$ matrices [5]), the TCR-pMHC structure prediction algorithm TCRdock [7], the sequence-based machine learning algorithms for TCRs (TAPIR [19], NetTCR [18], and TULIP [17]), and the structure-based inverse folding models (ESM-IF1 [16] and ProteinMPNN [15]). Since all of these models are unsupervised in nature (except for Łuksza *et al* activity predictions for TCRs 1-7), we do not expect predictions to be in the same units as the affinity and activity values, and thus, we use Spearman correlation between data and predictions as a metric for model performance.

For predicting TCR-pMHC binding affinities (Table S2), BLOSUM62 leads for the 1G4 TCR system, while template-guided TCRdock tops on the A6 TCR systems. HERMES-*relaxed* with no noise (0.00) achieves slightly lower Spearman correlation but is statistically indistinguishable from these best performers (p-value $> 0.05$ associated with the difference in the Fisher's z-transformed Spearman correlations). For 1G4, ESM-IF1 and the Łuksza *et al*'s TCR-specific $d \cdot M$ and $M$ substitution matrices are similarly competitive. HERMES-*relaxed* 0.00 consistently outperforms the sequence-based models (TAPIR, NetTCR-2.2), ProteinMPNN, and TCRdock-*no template* (the TCRdock model without using structural templates similar to the query data). Lastly, choosing the minimum-Rosetta-energy conformation for HERMES-*relaxed* yields correlations with the data that are comparable to the baseline HERMES-*relaxed* model that averages over the energies of 100 different realizations of relaxed peptides (Fisher's z-test p-value $> 0.05$). Table S2 shows these benchmarking results in detail.

For predicting T-cell activity (Table S3), HERMES-*fixed* without noise (0.00) achieves the best performance for TCR1, TCR2, and TCR3, and proteinMPNN-0.02 shows competitive performances for TCR1 and TCR3. ProteinMPNN-0.02 leads for TCR6 and H2-scDb, while HERMES-*fixed* 0.00 achieves slightly lower but statistically indistinguishable Spearman correlations (Fisher's z-test p-value $> 0.05$). All models struggle to provide reliable predictions for TCR7, with the exception of ProteinMPNN 0.02, which attains a moderate Spearman correlation $\rho = 0.31$ when scoring one peptide amino acid at a time (eq. S6), and $\rho = 0.29$, when scoring the peptide and the TCR together (eq. S8). For TCR4, which has a poor structural model (AF3 PAE = 2.07), all structure-based models fail to produce reliable predictions, whereas BLOSUM62 achieves a moderate Spearman correlation $\rho = 0.33$. The Łuksza *et al*'s $d \cdot M$ substitution matrix overall performs well on the TCR 1-7 benchmark; however, because it was obtained by fitting on these same data, its predictions are not strictly zero-shot. The structure-based models HERMES and ProteinMPNN consistently outperform the sequence-based models (TAPIR, NetTCR-2.2, and TULIP), as well as the structure-based ESM-IF1. Lastly, choosing the minimum-Rosetta-energy conformation for HERMES-*relaxed* model yields a similar performance to the baseline HERMES-*relaxed* model that averages over the energies of 100 different realizations of relaxed peptides. Table S3 shows these benchmarking results in detail.

Our results do not indicate a clear preference between HERMES-*fixed* and HERMES-*relaxed*: HERMES-*fixed* predicted T-cell activity better, while HERMES-*relaxed* predicted binding affinity between TCR and pMHC better. However, it is important to note that in all systems tested here, the scored peptides are highly similar to the wild-type peptides, with a maximum Hamming distance of just one amino acid from a wild-type peptide found in one of the available structural templates. As a result, relaxation may not be essential for achieving high performance. Interestingly, noised HERMES models underperform compared to the un-noised counterparts. This is in contrast to prior observations, where noised models achieve higher performances in predicting the stability effect of mutations in proteins [1]. Moreover, we observe that ProteinMPNN predictions are robust to the choice of scoring scheme, i.e., scoring peptide amino acids one at a time $E^{(\mathrm{MPNN})}_{\mathrm{pep\ 1\text{-}by\text{-}1}}$ (eq. S6), the whole peptide simultaneously $E^{(\mathrm{MPNN})}_{\mathrm{pep}}$ (eq. S7), or peptide and the TCR interface together $E^{(\mathrm{MPNN})}_{\mathrm{pep\ and\ tcr}}$ (eq. S8); see Fig. S1.

Lastly, we benchmarked all four HERMES models and two ProteinMPNN models (using peptide-scoring schemes eq. S7 with different noise levels) on AF3-predicted structures, for both activity and affinity predictions (Fig. S4). We generated the AF3 structures both with and without template guidance, and the resulting predictions made with these AF3 structures were compared to performances using the corresponding experimental structures. AF3 produces accurate models for all complexes except the H2-scDb Fab bound to pMHC, for which the docking orientation and conformation were incorrect (Table S4). Expectantly, we saw a significant drop in model performances for both HERMES and ProteinMPNN when using the inaccurate AF3 predictions for H2-scDb (Fig. S4). For both HERMES and ProteinMPNN, replacing experimental structures with AF3 predictions altered predictions for the 1G4 and A6 TCRs binding to pMHC variants, but the changes were not statistically significant (p-value$> 0.05$ associated with the difference in the Fisher's z-transformed Spearman correlations); see Fig. S4. Finally, for TCR1, we observe no differences in performance at predicting T-cell activity between using the AF3 structures vs. the experimental structures. Comparisons for TCRs 2-7 show insignificant differences between predictions using AF3 models with and without templates.

In Table S4 we report, where available, the $C_\alpha$ RMSD between AF3-predicted and experimentally determined structures, together with AF3 confidence metrics computed with and without template guidance. The confidence metrics include the TCR–pMHC interface predicted alignment error (PAE), the interchain predicted TM-score (ipTM), and the predicted TM-score (pTM; not interface-specific). We found ipTM and pTM to be closely related (Table S4). Because structure-based models such as HERMES and ProteinMPNN rely on accurate input structures, Fig. S4B plots their best-performing variants against AF3 confidence. The AF3 model for TCR4—on which both HERMES and ProteinMPNN perform poorly—exhibits the second-lowest ipTM confidence (after H2-scDb). However, the limited range of available AF3 confidence values prevents definitive conclusions about how these metrics impact the accuracy of predictions for peptide mutation-effects on T-cell responses.

**Benchmarking HERMES designs against alternative models.** Although we did not experimentally test the designs by the different models, we used both peptide presentation by MHC predicted by NetMHCPan and TCRdock's PAE as benchmarking metrics across different design methods. We note that these are only computational metrics and should not be taken as ground truth; this is particularly true for results using TCRdock PAE as a proxy for binding, which are much more likely farther from the ground truth than the easier-to-predict MHC presentation.

Figs. S13, S14, and S15 show the fraction of peptides meeting each criterion for designs produced by various HERMES models, ProteinMPNN, BLOSUM62, and the system's MHC peptide preferences. Consistently, across the three systems, and for all models, the fraction of designs passing the PAE threshold decreases with increasing Hamming distance from the wild-types' sequences (Figs. S13, S14, and S15; panels E, and G). Also consistently, but more surprisingly, models trained with Gaussian noise applied to the structure—for both HERMES and ProteinMPNN—perform worst than their noise-less counterparts (Figs. S13, S14, and S15; panels E, and G).

HERMES models are consistently the best at designing peptides that are scored well for presentation by the respective MHC alleles (panel C of Figs. S13, S14, and S15). No single model stands out as being consistently best across the three systems at designing peptides above the TCRdock PAE threshold: HERMES outperforms ProteinMPNN in the NY-ESO system (Fig. S13), performs similarly in the EBV system (Fig. S14), and performs worse in the MAGE system (Fig. S15). BLOSUM62 makes worse designs compared to HERMES and ProteinMPNN based on TCRdock PAE scores. Surprisingly, ESM-IF1 designs have the lowest success in MHC presentation, but show better TCRdock PAE scores compared to HERMES-*relaxed* for the MAGE system (Figs. S13, S14, and S15). Designs sampled from the MHC motif, while scoring well on MHC presentation, consistently score poorly for TCRdock PAE. We expected HERMES-*relaxed* to design better binders than HERMES-*fixed* at greater sequence distances from the wild-types. However, this holds only for the NY-ESO system (Fig. S13) and not for EBV or MAGE (Figs. S14, S15). Overall, a broader benchmarking study, ideally with experimental validation, is needed to rigorously evaluate the effectiveness of different peptide design methods.

To assess how using AF3 structures as templates influence design, we ran the HERMES-*fixed* design protocol using experimental and AF3 structures as templates and compared the resulting peptide likelihood matrices (as position weight matrices or PWMs) for the three systems (Fig. S12). Across NY-ESO, EBV, and MAGE, the resulting PWMs were largely consistent across the two template choices: most peptide positions showed the same preferences, and when variability did appear it was typically confined to chemically similar residues. There are two noteworthy exceptions: in NY-ESO, the top choice in position 7 of the peptide was flipped from Threonine and Aspartic Acid when AF3 template was used (Fig. S12A). In MAGE, using the AF3 model introduced a pronounced preference for glutamic acid at position 1 (Fig. S12C), a residue that we have already shown to be important for eliciting T-cell activity (Fig. 4). Aside from these cases, AF3-derived structures preserve the peptide specificity landscape learned by HERMES.

### V. EXPERIMENTAL PROTOCOL TO MEASURE PEPTIDE ACTIVITIES

**Artificial antigen-presenting cell lines.** We obtained the full-length coding sequences of HLA-A*02:01, HLA-A*01:01, and HLA-B*35:01 from the IMGT/HLA database [22]. Gene fragments were synthesized by Genscript and were cloned into the lentiviral pLVX-EF1$\alpha$-IRES-Puro (Clontech) vector. We used Lipofectamine 3000 Transfection Reagent (Invitrogen) to transduce the 293T packaging cell line (ATCC CRL-3216) with the psPAX2 packaging plasmid (Addgene plasmid #12260), the pMD2.G envelope plasmid (Addgene plasmid #12259), and the HLA-encoding lentiviral vector. The lentivirus-containing supernatant was collected after 24 and 48 hours post-transfection and was filtered through a 0.45um SFCA filter (Thermo Scientific). The lentivirus was concentrated using Lenti-X Concentrator (Takara Bio) according to the manufacturer's protocol, resuspended in DPBS (Gibco), and stored at -80C for future use. To generate monoallelic HLA cell lines for HLA-A*02:01 and HLA-B*35:01, HLA-deficient K562 cells (ATCC CCL-243) were transduced with lentivirus. Since K562 cells have a high expression of both MAGE-A3 and Titin antigens, we transduced murine EL4 cell lines (ATCC TIB-39) with HLA-A*01:01 lentivirus to produce HLA-A*01:01 presenters lacking the natural expression of the tested antigens. At 48 hours post-transduction, K562 cells were transferred to Iscove's Modified Dulbecco's Medium containing 10% FBS, 2mM L-glutamine, 100 U/mL penicillin/streptomycin, and 2 $\mu$g/mL puromycin (Gibco). EL4 cells were transferred to RPMI media containing 10% FBS, 2mM L-glutamine, 100 U/mL penicillin/streptomycin, and 2 $\mu$g/mL puromycin (Gibco). The antibiotic selection continued for one week before transferring cells to the antibiotic-free media.

**TCR-expressing Jurkat 76.7 cell lines.** Nucleotide sequences of TCRa and TCRb chains were generated using stitchr [23]. Both chains were modified to incorporate murine constant segments (TRAC*01 and TRBC2*01) to enhance TCR surface expression. For NYESO and EBV-specific TCRs, the TCRa and TCRb chains were connected through a P2A ribosomal skip motif, synthesized as a single bicistronic gene fragment by Genscript, and cloned into the lentiviral pLVX-EF1a-P2A-mCherry-IRES-Puro (Clontech). For MAGE-specific TCR, gene fragments for the TCRa and TCRb chains were separately synthesized by Genscript and cloned into either the lentiviral pLVX-EF1a-P2A-mCherry-IRES-Puro (Clontech) or pLVX-EF1$\alpha$-IRES-G418 (modified in-house from pLVX-EF1$\alpha$-IRES-Puro). The 293T packaging cell line (ATCC CRL-3216) was transfected with TCR-encoding lentiviral vectors, psPAX2 packaging plasmid (Addgene plasmid #12260), and pMD2.G envelope plasmid (Addgene plasmid #12259) using Lipofectamine 3000 Transfection Reagent (Invitrogen). The lentivirus-containing supernatant was collected after 24 and 48-hours after transfection, filtered through a 0.45$\mu$m SFCA filter (Thermo Scientific), concentrated using Lenti-X Concentrator (Takara Bio) according to the manufacturer's protocol, resuspended in DPBS (Gibco), and stored at -80C for future use. To generate Jurkat cells expressing the paired TCR, TCR-null CD8-positive Jurkat 76.7 cells with NFAT-eGFP reporter were transduced with lentivirus. At 48 hours post-transduction, cells were transferred to RPMI media containing 10% FBS, 2mM L-glutamine, 100 U/mL penicillin/streptomycin, and 1 $\mu$g/mL puromycin (Gibco). For MAGE-specific cell line generation Jurkat cells were co-transduced with lentiviruses encoding TCRa and TCRb chains and cultured with both 1 $\mu$g/mL puromycin and 500 $\mu$g/mL Genectin (Gibco). The antibiotic selection continued for one week. The abTCR expression on the surface was confirmed by mCherry expression and staining with a 1:100 dilution of anti-mouse TCRb APC-Fire750 (clone H57-597, 109246, Biolegend).

**Specificity validation of TCR-expressing Jurkat.** Original cognate peptides and their variants were synthesized as a crude peptide library with by Genscript, and then diluted to a 4mM stock solution in DMSO (Sigma). To examine the specificity of the TCR-expressing Jurkat cell lines, we co-cultured 100,000 Jurkat cells with equal amounts of artificial APCs expressing the corresponding HLA allele in a 96-well round-bottom tissue culture plate. We tested each specificity—MAGE, EBV, NYESO—under 96 different conditions. These included one or two positive control original peptides and an unstimulated "no peptide" control. The cells were cultured in RPMI media supplemented with 10% FBS, 2mM L-glutamine, 100 U/mL penicillin/streptomycin, and 1 $\mu$g/ml each of anti-human CD28 (BD Biosciences, 555725) and CD49d (BD Biosciences, 555501). The co-cultures were pulsed with 1$\mu$M peptide in triplicate. We added DMSO to unstimulated control wells (no peptide) at concentrations equivalent to those in the peptide-stimulated wells. After incubating the cells for 24 hours at 37°C and 5% CO2, we washed them once in DPBS (Gibco) and resuspended them in 150$\mu$l DPBS for flow cytometry analysis using a custom-configured BD Fortessa with FACSDiva software (Beckton Dickenson). We determined GFP percentages using FlowJo version 10.8.1 software (BD Biosciences). Alternatively, we measured the GFP percentage in wells of a 96-well round-bottom plate using Cellcyte X (Echo), using 4x objective in spheroid mode, acquisition in green channel (800 ms exposure 3dB gain). Image analysis was done with CellCyte Studio software with total spheroid area recipe and standard defaults (50 a.u. Contrast Sensitivity, 2 a.u Smoothing, 100 $\mu$m$^2$ filled hole size 100 $\mu$m Min. object size), relative GFP+ spheroid area was used as output. For NY-ESO, we compared the T-cell activity level measured using flow cytometry with fluorescence microscopy, and show consistency between the two (Fig. S7). Given the consistency between the two experimental approaches, we relied on fluorescence microscopy only to measure GFP levels induced by different peptides for MAGE and EBV.

**Selecting the GFP expression threshold for T-cell activity.** Experimental measurements are done across three replicates, and the reported GFP levels in Figs. 4, S8 for each condition are averaged over the three replicates. We consider a peptide to have activated T-cells if its induced replicate-averaged GFP level met two criteria: 1) it exceeded 0.5%, and 2) it surpassed the mean plus three standard deviations of the GFP level negative controls (no-peptide). According to these criteria, the GFP activation cutoffs were 0.5% for the NY-ESO system, 0.64% for the EBV system, and 0.76% for the MAGE system.

### VI. EXPERIMENTAL ACTIVITIES OF THE DESIGNED PEPTIDES

**Comparing design success rates across the three TCR-pMHC systems.** Among the designed peptides, 50% in the NY-ESO system (Fig. 4A), 16% in the EBV system (Fig. 4B), and 26% in the MAGE system induce significant

T-cell activities. However, it should be noted that the designed NY-ESO peptides were closer in sequence to their wild-type templates (2-7 amino acid differences) than the peptides designed for the EBV (3-9 differences) or MAGE (3-8 differences) systems. We find that the activity of designs decays with increasing distance from the respective wild-type, with almost no designs beyond 5 amino-acid distance from the wild-types activating their respective T-cells (Fig. 4, S9, S10, S11). This underscores the difficulty of exploring peptide variants far from the wild-type sequences. If any positive designs do exist at these greater distances, they represent an exponentially smaller fraction of the sequence space, and when in complex with the TCR-MHC, are likely to adopt a different binding mode compared to that of the wild-type.

Minor adjustments to the design protocols, aimed at better exploring the design space, may have also contributed to the observed differences in performance across the systems. For NY-ESO, which achieved the highest design success rate, peptides with highest PAE scores were chosen for experimental validation, whereas in EBV and MAGE, peptides were sampled uniformly above the set PAE thresholds. In MAGE, fixing the amino-acid at position 1 to a Glutamic Acid (E) for the one-third of the designs led to an elevated success rate in this subset (Fig. S11D); $E_1$ is shared between the two wild-type peptides and this design choice allowed us to assess the importance of this amino acid for function.

**The design potential of different HERMES models.** In panel A of Figures S9 (NY-ESO), S10 (EBV), and S11 (MAGE), we show the fidelity of designs using the four HERMES models: HERMES-*fixed* with no noise, HERMES-*fixed* with 0.5 Å noise, HERMES-*relaxed* with no noise, and HERMES-*relaxed* with 0.5 Å noise. Although PyRosetta relaxations allows HERMES-*relaxed* to explore the sequence space more broadly, the resulting designs that are made at greater distances from the wild-types seldom succeed in activating T-cells. This pattern is also observed with using models trained with 0.5 Å Gaussian noise. Indeed, successful designs lie within five amino acid substitutions from the wild-types, and for these designs, the un-noised HERMES-*fixed* model demonstrates the highest overall fidelity. To better explore the peptide energy landscape and to design sequences further from the wild-types, it may be necessary to adopt a more specialized relaxation model that accounts for the intricacies of TCR-pMHC poses, while being consistent with the neural network's model of protein environments.

**Comparing TCRdock and AF3 as design filters.** Using our T-cell activity measurements as the ground truth, we sought to quantify how well the TCRdock and AF3 PAE scores can be used as predictors of T-cell activation and filtering mechanisms for peptide design. For this purpose, we focus primarily on the NY-ESO system, for which we validated a balanced mix of positive and negative designs, chosen based on their TCRdock PAE scores. Here, we found that TCRdock PAE could moderately distinguish true positive from true negative designs (AUROCs = 0.91 and 0.88, Fig. S16A). Notably, using TCRdock PAE resulted in a very low false negative rate (0.03%) but a considerably higher false positive rate (50%); see Fig. S8A top. AF3 PAE scores performed equivalently to TCRdock PAE in differentiating positives from negatives (AUROCs = 0.89 and 0.94, Fig. S16A), though they correlated only moderately with TCRdock's PAE values predictions (Spearman correlation $\rho = 0.5$, Fig. S8C). Indeed, some peptides deemed positive by TCRdock were classified as negative by AF3, and vice versa (Fig. S8C).

When restricting the analysis to peptides that were deemed as positive based on their TCRdock and AF3 PAE scores, we did not expect either TCRdock or AF3 PAE to effectively discriminate between peptides that did or did not elicit a T-cell response in our experimental assay (i.e., true vs. false positives). However, we found that both TCRdock PAE and AF3 PAE scores retained significant predictive power in the NY-ESO system (AUROCs = 0.81 and 0.77, and AUROCs = 0.80 and 0.89, Fig. S16A). In the EBV and MAGE systems, the TCRdock and AF3 PAE values for peptides that we classified positive showed lower predictive power in distinguishing true from false positives compared to the NY-ESO system (Fig. S16B-C). The PAE scores of TCRdock and AF3 differed most markedly in the MAGE system (Spearman correlation $\rho = 0.09$ for MAGE), with TCRdock assigning relatively unfavorable PAE scores to the wild-type peptides, whereas AF3 ranked those same peptides near the top (Fig. S8I).

The PAE values reported by the structure prediction algorithms have been shown to be noisy indicators of protein folding fidelity and design quality [24]. While larger experimental libraries will be needed to draw definitive conclusions, it appears that PAEs from TCRdock and AF3 have some complementary predictive strengths, and therefore, combining their information could lead to more robust design decisions. Moreover, both of these scores appear to be uncorrelated or only weakly correlated with HERMES scores across systems (Figs. S8, S13H, S14H, S15H), suggesting that AF3 and TCRdock PAE's can serve as complementary filters to HERMES during design.

**HERMES energy as design filter.** We further sought to assess whether HERMES energy score itself could serve as a predictor of T-cell activation and as a filtering criterion for design. For each of the three systems, we scored all peptides we designed with all four combinations of HERMES protocols, regardless of which protocol was originially used for design. For each system, we then computed two sets of ROC curves, separately for peptides designed with HERMES-*fixed* and those designed with HERMES-*relaxed*. We found that scores from the HERMES-*relaxed* models consistently performed poorly in predicting T-cell activation, with only one out of six cases achieving an AUROC better than random; HERMES-*relaxed* 0.00 as scoring criterion on the peptides designed with HERMES-*fixed* in the EBV system showing AUROC = 0.66 (Fig. S16B). In contrast, HERMES-*fixed* 0.00 was consistently the best filtering criterion, outperforming even TCRdock and AF3 PAE score in several cases: in the EBV system, HERMES-*fixed* 0.00 scores on peptides designed by HERMES-*fixed* reached AUROC = 0.74; in the MAGE system, AUROC were 0.65 and 0.89 for and peptides designed with HERMES-*fixed* and HERMES-*relaxed*, respectively (Fig. S16). These results highlight two main takeaways: first, that HERMES-*fixed* 0.00 is the most reliable model for both scoring and designing peptides to elicit T-cell response; second, selecting high-scoring peptides using this model can improve hit rates, though potentially at the cost of reduced diversity in the peptide pool.

### VII. T-CELL AND MHC SPECIFICITY AS THE ENTROPY OF THE ASSOCIATED PEPTIDE DISTRIBUTION

We characterize the specificity of a TCR-MHC complex and that of the MHC by the diversity of peptides that they can recognize. For TCR-MHC specificity, we starting with a TCR-pMHC structural template, and use HERMES-*fixed* model to generates a peptide position weight matrix (PWM) that captures the ensemble of peptides recognized by the TCR and presented by the MHC. In Figure 5, we analyzed 105 human and 36 mouse MHC class I TCR-pMHC structures—curated by the TCRdock template database [7]—which included only MHC alleles represented in at least two structures. For each MHC allele in this set, we then characterized the distribution of peptides that the MHC can present (independently of the TCR) using PWMs inferred by the MHC Motif Atlas [11]. In both cases, we use the entropy of these PWMs, $S = -\sum_{\alpha,i} p_i(\alpha) \log_2 p_i(\alpha)$, as a proxy for degeneracy of the target (TCR-MHC or MHC), where $\alpha \in \{1, \ldots 20\}$ denotes the amino acid types, and $i \in \{1, \ldots, \ell\}$ are positions along the peptide.

### VIII. PREDICTING MHC-I PEPTIDE PRESENTATION PREFERENCES WITH HERMES.

HERMES is designed to be broadly applicable to diverse protein-interaction contexts. As such, we attempted to use HERMES-*fixed* 0.00 model to recapitulate the ensemble of peptides presented by an MHC-I allele. To do so, we gathered 416 peptide-MHC-I crystal structures from the TCR3D database [25] (Table S7). For each structure, we computed a peptide position position weight matrix (PWM) with the HERMES-*fixed* 0.00 design protocol, conditioning on the surrounding MHC structure and the masked wild-type peptide as the template. The resulting PWMs were then averaged across complexes that shared the same MHC allele and peptide length to produce an allele-specific peptide motif (for a given peptide length).

We compared these HERMES-generated PWMs with the corresponding PWMs from the MHC Motif Atlas [11], which we consider as ground truth; we used relative entropy (KL-Divergence), $D_{KL} = \sum_{\alpha,i} p_i^{\text{motif}}(\alpha) \log_2 \frac{p_i^{\text{motif}}(\alpha)}{p_i^{\text{HERMES}}(\alpha)}$ as well as the entropy difference between the two PWMs as metrics; here, $p_i^{(\cdot)}(\alpha)$ is the probability of amino acid $\alpha$ at position $i$ of a peptide, given the morel specified in the superscript. When structural coverage is extensive (roughly more than 30 structures of distinct peptides with a given MHC allele), HERMES-derived motifs align closely with those from MHC Motif Atlas, as quantified by the low KL-divergence and comparable entropy values (Fig. S17). Under sparse coverage, HERMES motifs begin to underestimate the peptide diversity that can be presented by an MHC allele (measured by motif entropy), a limitation we attribute to incomplete sampling of peptide backbone conformations by HERMES-*fixed* (Fig. S17C). Overall, we see a clear pattern that the KL-Divergence increases with decreasing number of structural templates for a given MHC allele (Fig. S17B). Importantly, even with only a handful of templates, HERMES still captures substantially more diversity than a naïve baseline PWM generated by simply aligning the template peptides themselves (Fig. S17A). Moreover, in almost all cases, HERMES generated PWMs are more similar (lower KL-Divergence) to that of the MHC Motif Atlas than the naïve baseline PWM are (Fig. S17A).

These results suggest that accurate modeling of MHC presentation with HERMES-*fixed* design protocol requires both sufficient structure templates and peptide diversity. It should be noted that MHC Motif Atlas's allele-specific motifs–derived from the large body of peptide binding data curated for individual MHC-I alleles–are very reliable, so we continue to use them as the benchmark for assessing motif specificity. Developing design protocols that better sample peptide backbone conformations remains an important avenue for future work.

| **TCR / Ab** | **(PDBid, WT peptide)** | **MHC allele** |
|---|---|---|
| 1G4 TCR | (2BNR [26], SLLMWITQC); (2BNQ [26], SLLMWITQV) | HLA-A*02:01 |
| A6 TCR | (1AO7 [27], LLFGYPVYV); (1QSF [28], LLFGYPVAV) | HLA-A*02:01 |
| TCR1 | (5D2N [29], NLVPMVATV) | HLA-A*02:01 |
| TCR2 | (AF3 model, NLVPMVATV) | HLA-A*02:01 |
| TCR3 | (AF3 model, NLVPMVATV) | HLA-A*02:01 |
| TCR4 | (AF3 model, IMDQVPFSV) | HLA-A*02:01 |
| TCR5 | (AF3 model, IMDQVPFSV) | HLA-A*02:01 |
| TCR6 | (AF3 model, IMDQVPFSV) | HLA-A*02:01 |
| TCR7 | (AF3 model, GRLKALCQR) | HLA-B*27:05 |
| H2-scDb (Fab) | (6W51 [21], HMTEVVRHC) | HLA-A*02:01 |

Table S1. **Structure templates used for the binding affinity and T-cell activity analyses.** The first two rows contain information related to the analyses on binding affinity data, the rest are related to T-cell activity data. Every structure identified by a 4-letter PDBid was downloaded from the Protein Data Bank. The raw PDB file for 5D2N [29] has two copies of the TCR-pMHC complex, so we deleted one before running any analysis. Similarly, the raw PDB file for 6W51 [21] has the Antibody and pMHC coordinates of two different copies within the crystal, so we expanded the crystal using PyMOL's `symexp` command and kept a single bound Antibody-pMHC complex. The sequence inputs used to fold TCR-pMHC complexes with AlphaFold3 [20] can be found in Dataset S1.

| Model type | Model | 1G4 TCR | A6 TCR |
|---|---|---|---|
| Substitution Matrix | BLOSUM62 | **0.83** (0.00) | 0.56* (0.02) |
| | Luksza et al. $M$ | 0.44* (0.12) | 0.24 (0.38) |
| | Luksza et al. $d \cdot M$ | 0.48* (0.08) | 0.29 (0.27) |
| Structure Predictor | TCRdock | 0.62* (0.02) | **0.88** (0.00) |
| | TCRdock-*no template* | 0.25 (0.39) | -0.14 (0.62) |
| Sequence-based | TAPIR | 0.29 (0.32) | 0.35 (0.19) |
| | NetTCR-2.2 | -0.25 (0.40) | -0.27 (0.36) |
| Structure-based | ESM-IF1 | 0.38* (0.17) | 0.17 (0.53) |
| | ProteinMPNN 0.02 Pep 1-by-1 | -0.01 (0.97) | 0.49 (0.05) |
| | ProteinMPNN 0.20 Pep 1-by-1 | 0.12 (0.69) | 0.29 (0.27) |
| | ProteinMPNN 0.02 Pep | -0.01 (0.97) | 0.49 (0.06) |
| | ProteinMPNN 0.20 Pep | 0.14 (0.63) | 0.29 (0.27) |
| | ProteinMPNN 0.02 Pep and TCR | 0.07 (0.81) | 0.54 (0.03) |
| | ProteinMPNN 0.20 Pep and TCR | -0.08 (0.79) | 0.26 (0.33) |
| Structure-based | HERMES-*fixed* 0.00 | 0.29 (0.31) | 0.63* (0.01) |
| | HERMES-*fixed* 0.50 | 0.19 (0.51) | -0.28 (0.29) |
| | HERMES-*relaxed* 0.00 | 0.72* (0.00) | 0.56* (0.03) |
| | HERMES-*relaxed* 0.50 | 0.65* (0.01) | 0.06 (0.83) |
| | HERMES-*relaxed*-min_energy 0.00 | 0.58* (0.03) | 0.71* (0.00) |
| | HERMES-*relaxed*-min_energy 0.50 | 0.52* (0.06) | 0.51 (0.04) |

Table S2. **Benchmarking of models for predicting TCR-pMHC binding affinities.** The table lists the Spearman correlation $\rho\,(\mathrm{p}-\mathrm{value})$ of models' predictions against experimentally measured binding affinities for (i) 1G4 TCR in complex with HLA-A*02:01 binding to the variants of the NY-ESO-1 peptide, and (ii) the A6 TCR in complex with HLA-A*02:01 binding to the variants of the Tax peptide. The HERMES model names specify the scoring scheme (fixed, relaxed, and relaxed with peptide energy computed from the relaxed conformation with the minimum Rosetta energy), and the noise amplitude (in Å) injected to the coordinates of the structure data during training. Similarly, the ProteinMPNN model names capture scoring scheme (1-by-1 (eq. S6), the whole peptide (eq. S7), and the peptide and TCR interface together (eq. S8)), and the noise amplitude injected to the training data. We show in **bold** the best model for each dataset, and any model whose performance is not significantly different from the best model (p-value$>$ 0.05 for the difference in the Fisher's z-transformed Spearman correlations) is indicated with an asterisk (*).

| Model type | Model | H2-scDb (Fab) | TCR1 | TCR2 | TCR3 | TCR4 | TCR5 | TCR6 | TCR7 |
|---|---|---|---|---|---|---|---|---|---|
| Sub. Matrix | BLOSUM62 | -0.05 (0.52) | -0.07 (0.47) | 0.14 (0.20) | 0.38 (0.00) | 0.33* (0.01) | 0.42* (0.00) | 0.55* (0.00) | -0.01 (0.89) |
| | Luksza et al. $M$ | 0.19 (0.02) | $^\dagger$0.25 (0.01) | $^\dagger$0.39 (0.00) | $^\dagger$0.53* (0.00) | $^\dagger$0.25* (0.03) | $^\dagger$0.27 (0.00) | $^\dagger$0.36* (0.00) | $^\dagger$0.01 (0.94) |
| | Luksza et al. $d \cdot M$ | 0.46* (0.00) | $^\dagger$0.50* (0.00) | $^\dagger$0.66* (0.00) | $^\dagger$0.40 (0.00) | $^\dagger$**0.38** (0.00) | $^\dagger$ **0.55** (0.00) | $^\dagger$0.42* (0.00) | $^\dagger$0.27* (0.00) |
| Struc. Pred. | TCRdock | N/A | 0.02 (0.83) | 0.13 (0.24) | 0.21 (0.03) | 0.18 (0.15) | -0.02 (0.83) | 0.19 (0.04) | 0.06 (0.50) |
| | TCRdock-*no template* | N/A | -0.03 (0.77) | 0.44 (0.00) | 0.11 (0.27) | -0.22 (0.08) | 0.28 (0.00) | 0.14 (0.14) | 0.06 (0.50) |
| Seq.-based | TAPIR | N/A | 0.36 (0.00) | 0.36 (0.00) | -0.00 (0.97) | 0.12 (0.32) | 0.39* (0.00) | 0.08 (0.42) | -0.01 (0.94) |
| | NetTCR-2.2 | N/A | -0.05 (0.59) | 0.20 (0.06) | 0.36 (0.00) | -0.21 (0.09) | 0.23 (0.01) | 0.20 (0.03) | -0.22 (0.02) |
| | TULIP | N/A | 0.12 (0.24) | 0.18 (0.09) | -0.12 (0.20) | 0.04 (0.73) | 0.24 (0.00) | 0.29 (0.00) | N/A |
| | TULIP* (from [17]) | N/A | 0.47* (0.00) | 0.44 (0.00) | 0.14 (0.07) | 0.23* (0.00) | 0.20 (0.01) | 0.09 (0.25) | N/A |
| Struc.-based | ESM-IF1 | 0.12 (0.13) | 0.43* (0.00) | 0.41 (0.00) | 0.19 (0.05) | -0.15 (0.22) | 0.28 (0.00) | 0.25 (0.01) | 0.08 (0.40) |
| | ProteinMPNN 0.02 1-by-1 | 0.53* (0.00) | 0.46* (0.00) | 0.53 (0.00) | 0.49* (0.00) | -0.23 (0.06) | 0.45* (0.00) | 0.42* (0.00) | **0.31** (0.00) |
| | ProteinMPNN 0.20 1-by-1 | 0.03 (0.65) | 0.27 (0.01) | 0.46 (0.00) | 0.40 (0.00) | -0.23 (0.06) | 0.26 (0.00) | **0.49** (0.00) | 0.02 (0.79) |
| | ProteinMPNN 0.02 Pep | **0.55** (0.00) | 0.46* (0.00) | 0.52 (0.00) | 0.49* (0.00) | -0.28 (0.02) | 0.45* (0.00) | 0.41* (0.00) | 0.31 (0.00) |
| | ProteinMPNN 0.20 Pep | 0.04 (0.64) | 0.27 (0.01) | 0.44 (0.00) | 0.40 (0.00) | -0.23 (0.05) | 0.25 (0.00) | 0.48* (0.00) | 0.02 (0.83) |
| | ProteinMPNN 0.02 Pep and TCR | 0.50* (0.00) | 0.44* (0.00) | 0.53 (0.00) | 0.34 (0.00) | -0.24 (0.05) | 0.51* (0.00) | 0.46* (0.00) | 0.29* (0.00) |
| | ProteinMPNN 0.20 Pep and TCR | -0.02 (0.84) | 0.20 (0.04) | 0.47 (0.00) | 0.30 (0.00) | -0.23 (0.06) | 0.32 (0.00) | 0.48* (0.00) | -0.02 (0.81) |
| Struc.-based | HERMES-*fixed* 0.00 | 0.46* (0.00) | **0.56** (0.00) | **0.71** (0.00) | **0.59** (0.00) | -0.06 (0.60) | 0.37 (0.00) | 0.42* (0.00) | -0.01 (0.95) |
| | HERMES-*fixed* 0.50 | 0.26 (0.00) | 0.42* (0.00) | 0.61* (0.00) | 0.54* (0.00) | 0.03 (0.83) | 0.26 (0.00) | 0.31 (0.00) | 0.02 (0.82) |
| | HERMES-*relaxed* 0.00 | 0.29 (0.00) | 0.31 (0.00) | 0.57 (0.00) | 0.55* (0.00) | -0.20 (0.11) | 0.37 (0.00) | 0.42* (0.00) | 0.14 (0.15) |
| | HERMES-*relaxed* 0.50 | 0.16 (0.04) | 0.37 (0.00) | 0.50 (0.00) | 0.45* (0.00) | -0.16 (0.20) | 0.23 (0.00) | 0.06 (0.49) | 0.08 (0.37) |
| | HERMES-*relaxed*-min_energy 0.00 | 0.18 (0.02) | 0.32 (0.00) | 0.56* (0.00) | 0.50* (0.00) | -0.10 (0.39) | 0.37 (0.00) | 0.45* (0.00) | 0.04 (0.65) |
| | HERMES-*relaxed*-min_energy 0.50 | 0.13 (0.09) | 0.39 (0.00) | 0.51 (0.00) | 0.46* (0.00) | -0.12 (0.34) | 0.15 (0.07) | 0.17 (0.07) | 0.08 (0.38) |

Table S3. **Benchmarking of models for predicting peptide-induced T-cell activities.** The table lists the Spearman correlation $\rho\,(\mathrm{p-value})$ of models' predictions against experimentally measured T-cell activities. The correlations for TCR1-7 are reported based on the $EC_{50}$ values obtained using the procedure outlined in Section II and Eq. S2, consistent with correlations reported in Fig. 3. In addition, we present the Spearman correlations between TULIP predictions and the fitted $EC_{50}$ values as reported in ref. [5], which are the same correlations reported in the TULIP paper [17] (TULIP* row). The TULIP* correlations also include datapoints for which T-cell responses were below the detection limit, and which we exclude from our analysis (gray points in Fig. 3). We note that the substitution matrix - with $(C)$ and without $(C/d)$ the position-based scaling $d$ - by Luksza et al. [5] was constructed precisely using experimental activity data from TCR1-7; we note predictions made on these measurements with a dagger$^\dagger$ to indicate that they are not representative of the matrix' ability to predict T-cell activity in new settings. The HERMES model names specify the scoring scheme (fixed, relaxed, and relaxed with peptide energy computed from the relaxed conformation with the minimum Rosetta energy), and the noise amplitude (in Å) injected to the coordinates of the structure data during training. Similarly, the ProteinMPNN model names capture scoring scheme (1-by-1 (eq. S6), the whole peptide (eq. S7), and the peptide and TCR interface together (eq. S8)), and the noise amplitude injected to the training data. We show in **bold** the best model for each dataset, and any model whose performance is not significantly different from the best model (p-value$>0.05$ for the difference in the Fisher's z-transformed Spearman correlations) is indicated with an asterisk (*).

| TCR / Ab | with templates RMSD | PAE | ipTM | pTM | without templates RMSD | PAE | ipTM | pTM |
|---|---|---|---|---|---|---|---|---|
| 1G4 TCR | 0.72 | 1.06 | 0.94 | 0.94 | 0.82 | 1.08 | 0.93 | 0.94 |
| 1G4 TCR w/ 9V on peptide | 0.72 | 1.07 | 0.94 | 0.94 | 0.82 | 1.08 | 0.93 | 0.94 |
| A6 TCR | 0.87 | 0.91 | 0.95 | 0.95 | 0.83 | 0.91 | 0.95 | 0.95 |
| A6 TCR w/ 8A on peptide | 0.67 | 1.00 | 0.94 | 0.94 | 0.73 | 1.03 | 0.94 | 0.95 |
| TCR1 | 0.92 | 1.52 | 0.88 | 0.91 | 1.12 | 2.46 | 0.82 | 0.86 |
| TCR2 | N/A | 1.63 | 0.86 | 0.89 | N/A | 1.86 | 0.84 | 0.87 |
| TCR3 | N/A | 0.84 | 0.95 | 0.95 | N/A | 0.86 | 0.95 | 0.95 |
| TCR4 | N/A | 2.07 | 0.80 | 0.83 | N/A | 2.01 | 0.84 | 0.86 |
| TCR5 | N/A | 1.17 | 0.91 | 0.92 | N/A | 1.19 | 0.91 | 0.93 |
| TCR6 | N/A | 1.14 | 0.92 | 0.94 | N/A | 1.16 | 0.91 | 0.93 |
| TCR7 | N/A | 1.37 | 0.84 | 0.87 | N/A | 1.42 | 0.89 | 0.91 |
| H2-scDb (Fab) | 13.89 | 19.44 | 0.48 | 0.54 | 11.57 | 19.2 | 0.47 | 0.55 |
| eng. TCR w/ MageA3 peptide | 0.62 | 1.03 | 0.92 | 0.94 | 0.69 | 1.03 | 0.92 | 0.94 |
| eng. TCR w/ Titin peptide | 0.64 | 1.05 | 0.92 | 0.94 | 0.65 | 1.03 | 0.92 | 0.94 |
| TK3 TCR w/ HPVG peptide | 1.05 | 1.11 | 0.92 | 0.94 | 1.05 | 1.05 | 0.93 | 0.94 |
| TK3 TCR w/ HPVG EQ5 peptide | 1.01 | 1.07 | 0.92 | 0.94 | 1.06 | 1.07 | 0.92 | 0.94 |

Table S4. **AlphaFold3 folding metrics for structures used in our analyses.** Root-mean-square deviation (RMSD) is computed between the $\alpha$-C's of AF3 predicted structures and the corresponding experimentally determined structure. The reported PAE values are averaged across the interfaces of TCR ($\alpha$ and $\beta$ chains) and pMHC. ipTM and pTM values are reported from the AF3's confidence output. For template-guided predictions, we used the default template cutoff in the AlphaFold Server (September 9th 2021). If an AF3 prediction incorrectly docked the TCR on the opposite side of the pMHC, the run was discarded and repeated. In the case of the H2-scDb bi-specific Fab-pMHC complex, although AF3 placed the Fab on the correct side of the pMHC, the docking pose was consistently inaccurate, resulting in high RMSD and PAE values. This was the only system for which AF3 failed to produce a realistic structure model. Additional details on experimental structures, peptide, TCR, and MHC sequences used in AF3 runs can be found in Tables S1, S5, Dataset S1. Example AF3-generated structures are shown in Figure S5.

| System | TCR | (PDBid, WT peptide) | MHC allele |
|---|---|---|---|
| NY-ESO | 1G4 TCR | (2BNR [26], SLLMWITQC); (2BNQ [26], SLLMWITQV) | HLA-A*02:01 |
| EBV | TK3 TCR | (3MV7* [30], HPVGEADYFEY); (4PRP* [31], HPVGQADYFEY) | HLA-B*35:01 |
| MAGE | eng. TCR | (5BRZ* [32], EVDPIGHLY); (5BS0* [32], ESDPIVAQY) | HLA-A*01:01 |

Table S5. **Structure templates used for peptide design.** For the PDB entries marked with an asterisk (*), we used the curated structures from the TCRdock database [7] instead of the PDB entries.

| MHC allele | (PDBid, WT peptide) |
|---|---|
| HLA-A*01:01 | (5BS0, ESDPIVAQY); (5BRZ, EVDPIGHLY) |
| HLA-A*02:01 | (4FTV, LLFGYPVYV); (5D2L, NLVPMVATV); (2F53, SLLMWITQC); (5TEZ, GILGFVFTL); (6VQO, HMTEVVRHC); (3HG1, ELAGIGILTV); (2P5E, SLLMWITQC); (7N6E, YLQPRTFLL); (6RP9, SLLMWITQV); (5C08, RQWGPDPAAV); (3H9S, MLWGYLQYV); (5C0B, RQFGPDFPTI); (6EQA, AAGIGILTV); (5EUO, GILGFVFTL); (3D39, LLFGPVYV); (6VRN, HMTEVVRHC); (5YXU, KLVALGINAV); (5EU6, YLEPGPVTV); (1BD2, LLFGYPVYV); (1OGA, GILGFVFTL); (2VLK, GILGFVFTL); (5MEN, ILAKFLHWL); (3D3V, LLFGPVYV); (1AO7, LLFGYPVYV); (5ISZ, GILGFVFTL); (5JHD, GILGFVFTL); (6DKP, ELAGIGILTV); (1QSE, LLFGYPRYV); (3QDM, ELAGIGILTV); (5JZI, KLVALGINAV); (2P5W, SLLMWITQC); (5C0C, RQFGPDWIVA); (7N1E, RLQSLQTYV); (4QOK, EAAGIGILTV); (5E6I, GILGFVFTL); (2VLR, GILGFVFTL); (6RSY, RMFPNAPYL); (3QEQ, AAGIGILTV); (3O4L, GLCTLVAML); (5C0A, MVWGPDPLYV); (5C07, YQFGPDFPIA); (5D2N, NLVPMVATV); (2F54, SLLMWITQC); (2VLJ, GILGFVFTL); (5NHT, ELAGIGILTV); (6TRO, GVYDGREHTV); (7N1F, YLQPRTFLL); (3GSN, NLVPMVATV); (6D78, AAGIGILTV); (3QDG, ELAGIGILTV); (5NQK, ELAGIGILTV); (5HHO, GILEFVFTL); (2PYE, SLLMWITQC); (5HHM, GILGLVFTL); (2GJ6, LLFGKPVYV); (5NMF, SLYNTIATL); (3UTS, ALWGPDPAAA); (5E9D, ELAGIGILTV); (6RPA, SLLMWITQV); (1QSF, LLFGYPVAV); (4EUP, ALGIGILTV); (5YXN, KLVALGINAV); (2BNQ, SLLMWITQV); (6RPB, SLLMWITQV); (5HYJ, AQWGPDPAAA); (3QFJ, LLFGFPVYV); (3UTT, ALWGPDPAAA); (3QDJ, AAGIGILTV); (4L3E, ELAGIGILTV); (5C09, YLGGPDFPTI); (6VRM, HMTEVVRHC); (1QRN, LLFGYAVYV); (2BNR, SLLMWITQC); (3PWP, LGYGFVNYI); (6AMU, MMWDRGLGMM); (6AM5, SMLGIGIVPV); (4MNQ, ILAKFLHWL); (5NME, SLYNTVATL); (5NMG, SLFNTIAVL) |
| HLA-A*11:01 | (5WKH, GTSGSPIINR); (5WKF, GTSGSPIVNR) |
| HLA-A*24:02 | (3VXR, RYPLTFGWCF); (3VXM, RFPLTFGWCF); (3VXS, RYPLTLGWCF) |
| HLA-B*07:02 | (6VMX, RPPIFIRRL); (6AVF, APRGPHGGAASGL); (6AVG, APRGPHGGAASGL) |
| HLA-B*08:01 | (1MI5, FLRGRAYGL); (3FFC, FLRGRAYGL); (4QRP, HSKKKCDEL); (3SJV, FLRGRAYGL) |
| HLA-B*27:05 | (4G8G, KRWIILGLNK); (4G9F, KRWIIMGLNK) |
| HLA-B*35:01 | (6BJ2, IPLTEEAEL); (4PRP, HPVGQADYFEY); (3MV7, HPVGEADYFEY); (3MV9, HPVGEADYFEY); (3MV8, HPVGEADYFEY) |
| HLA-B*35:08 | (4PRH, HPVGDADYFEY); (4PRI, HPVGEADYFEY) |
| HLA-B*44:05 | (3DXA, EENLLDFVRF); (3KPR, EEYLKAWTF); (3KPS, EEYLQAFTY) |
| H2Db | (3PQY, SSLENFRAYV); (5WLG, SQLLNAKYL); (5TJE, KAVYNFATM); (7JWI, ASNENMETM); (5SWS, ASNENMETM); (5TIL, KAPYNFATM); (7JWJ, ASNENMETM); (6G9Q, KAPYDYAPI); (5M02, KAPFNFATM); (5M00, KAVANFATM); (5M01, KAPANFATM) |
| H2Kb | (3PQY, SSLENFRAYV); (5WLG, SQLLNAKYL); (5TJE, KAVYNFATM); (7JWI, ASNENMETM); (5SWS, ASNENMETM); (5TIL, KAPYNFATM); (7JWJ, ASNENMETM); (6G9Q, KAPYDYAPI); (5M02, KAPFNFATM); (5M00, KAVANFATM); (5M01, KAPANFATM) |
| H2Ld | (2OI9, QLSPFPFDL); (4NHU, GGGAPWNPAMMI); (4MVB, QPAEGGFQL); (4MXQ, SPAPRPLDL); (2E7L, QLSPFPFDL); (4N5E, VPYMAEFGM); (6L9L, SPSYAYHQF); (3E3Q, QLSPFPFDL); (3E2H, QLSPFPFDL); (3TPU, FLSPFWFDI); (3TFK, QLSDVPMDL); (4MS8, SPAEAGFFL); (4N0C, MPAGRPWDL); (3TJH, SPLDSLWWI) |

Table S6. **Structure templates to compute TCR-MHC specificities.** The table list the structures used to compute TCR-MHC specificities in Fig. 5 for different MHC class I alleles in humans and mice. Structures were collected from the TCRdock curated database [7].

| MHC allele | Peptide Length | PDBid |
|---|---|---|
| HLA-A*01:01 | 9 | 3BO8; 4NQV; 4NQX |
| HLA-A*01:01 | 10 | 6AT9; 6MPP |
| HLA-A*02:01 | 9 | 1B0G; 1DUZ; 1EEY; 1EEZ; 1HHG; 1HHI; 1HHJ; 1HHK; 1I1F; 1I1Y; 1I7R; 1I7T; 1I7U; 1JHT; 1QEW; 1QR1; 1S9W; 1S9X; 1S9Y; 1TVB; 2C7U; 2GTW; 2GTZ; 2GUO; 2V2W; 2V2X; 2VLL; 2X4O; 2X4S; 2X4U; 3BGM; 3FT2; 3FT3; 3FT4; 3GSO; 3GSQ; 3GSR; 3GSU; 3GSV; 3GSW; 3GSX; 3H7B; 3H9H; 3HPJ; 3I6G; 3IXA; 3KLA; 3MR9; 3MRB; 3MRC; 3MRD; 3MRE; 3MRF; 3MRG; 3MRH; 3MRI; 3MRJ; 3MRK; 3MRL; 3MYJ; 3PWJ; 3PWL; 3PWN; 3QFD; 3TO2; 3V5D; 3V5H; 3V5K; 4I4W; 4K7F; 4L29; 5E00; 5EU3; 5HHN; 5HHP; 5HHQ; 5MEO; 5MEP; 5N6B; 5WSH; 6SS7; 6SS8; 6SS9; 6SSA; 6VR1; 6VR5; 6Z9W; 7EU2; 7LG2; 7LG3; 7MJ6; 7MJ9; 7MKB; 7N1A; 7N1B; 7RTD; 7SA2; 7SIS; 7U1R; 7U21; 7UM2; 7ZUC; 8FU4; 8RBV |
| HLA-A*02:01 | 10 | 1HHH; 1I4F; 1JF1; 2CLR; 2GT9; 3BH8; 3GIV; 3I6K; 3MGO; 3MGT; 3MRM; 3MRN; 3MRO; 3MRP; 3MRQ; 3MRR; 3UTQ; 4GKN; 4GKS; 4JFO; 4JFP; 4JFQ; 5C0D; 5C0G; 5C0I; 5FDW; 5N1Y; 6AMT; 6G3J; 6G3K; 6TRN; 7M8S; 7Q98; 7UR1 |
| HLA-A*02:01 | 11 | 5D9S; 5EOT; 7MJ7 |
| HLA-A*02:06 | 9 | 6UJO; 6UJQ |
| HLA-A*03:01 | 9 | 3RL1; 7L1B; 7L1C; 7MLE; 7UC5 |
| HLA-A*03:271 | 10 | 8JHV; 8JHW |
| HLA-A*11:01 | 9 | 1Q94; 7S8R; 8I5E |
| HLA-A*11:01 | 10 | 1QVO; 5WJN; 7OW3; 7S8Q; 7S8S; 8RBU; 8RH6; 8RHQ |
| HLA-A*11:01 | 11 | 4MJ5; 4MJ6 |
| HLA-A*24:02 | 8 | 4F7T; 4WU5; 5HGA; 5HGB |
| HLA-A*24:02 | 9 | 2BCK; 3I6L; 6XQA; 7F4W; 7JYV; 7JYW; 7MJA; 7NMD |
| HLA-A*24:02 | 10 | 3NFN; 3VXN; 3VXO; 3VXP; 3WL9; 3WLB; 4F7M; 5HGD; 5HGH; 7JYU |
| HLA-A*26:01 | 9 | 8XES; 8XFZ; 8XKC |
| HLA-A*68:01 | 10 | 1TMC; 6EI2 |
| HLA-B*07:02 | 9 | 4U1K; 5EO0; 5EO1; 5WMN; 7LFZ; 7LG0; 7LGD; 7RZD; 7RZJ; 7S7D; 7S7E; 7S7F; 7S8A; 7S8E; 7S8F |
| HLA-B*08:01 | 8 | 8E13; 8E8I |
| HLA-B*08:01 | 9 | 3SKM; 3SKO; 4QRQ; 4QRT; 5WMQ; 5WMR; 7NUI |
| HLA-B*15:01 | 9 | 5TXS; 6UZS; 6VB3; 7XF3; 8ELG; 8ELH |
| HLA-B*15:02 | 9 | 6UZM; 6UZN; 6UZO; 6VB0; 6VB7; 6VIU |
| HLA-B*18:01 | 8 | 4XXC; 8RNG; 8ROO; 8ROP |
| HLA-B*18:01 | 9 | 6MT3; 8RNH |
| HLA-B*27:05 | 9 | 1JGE; 1UXS; 1W0V; 2A83; 2BSR; 2BST; 3B6S; 3BP4; 3DTX; 3LV3; 5IB1; 5IB2; 6PYW; 6Y2A; 7CIS |
| HLA-B*27:05 | 10 | 2BSS; 7CIQ; 7CIR; 7DYN |
| HLA-B*27:09 | 9 | 1K5N; 1W0W; 3B3I; 3BP7; 3CZF; 3HCV; 6Y27; 6Y2B |
| HLA-B*35:01 | 9 | 1A9B; 1A9E; 1CG9; 3LKN; 3LKO; 3LKP; 3LKQ; 3LKR; 3LKS; 7SIG; 8EMF; 8EMG; 8EMI; 8EMJ |
| HLA-B*35:08 | 13 | 3VFM; 3VFR; 3VFT; 3VFU; 3VFV |
| HLA-B*37:01 | 9 | 6MT4; 6MT5; 6MT6 |
| HLA-B*44:03 | 9 | 1SYS; 3KPO |
| HLA-B*44:05 | 9 | 1SYV; 7TUC |
| HLA-B*53:01 | 9 | 7R7V; 7R7W |
| HLA-B*57:01 | 9 | 6D2T; 7R7X |
| HLA-B*57:01 | 10 | 3VRI; 3VRJ; 6D29; 6V2O; 8F7M |
| HLA-B*57:03 | 9 | 2BVP; 6V2Q |
| HLA-B*58:01 | 8 | 7X1B; 7X1C |
| HLA-B*58:01 | 9 | 5VWH; 7WZZ; 7X00 |
| HLA-B*81:01 | 9 | 4U1I; 4U1L; 4U1S |
| H2-Db | 9 | 1BZ9; 1FFN; 1FFO; 1FFP; 1FG2; 1HOC; 1INQ; 1JPG; 1JUF; 1LD9; 1N5A; 1S7U; 1S7V; 1S7W; 1S7X; 1ZHB; 2F74; 2ZOL; 3BUY; 3CC5; 3CCH; 3CH1; 3CPL; 3FTG; 3QUK; 3QUL; 3TBS; 3TBT; 3TBV; 3TBW; 3TBY; 3WS3; 3WS6; 4HUU; 4HUV; 4HUW; 4HUX; 4HV8; 4IHO; 4L8B; 4L8C; 4L8D; 4NSK; 5E8N; 5E8O; 5E8P; 5JWD; 5MZM; 5WLI; 6G9R; 6L9M; 6L9N; 7N9J; 7P0A; 7P0T |
| H2-Db | 10 | 1N3N; 1WBX; 1WBY; 1YN6; 1YN7; 3L3H; 5JWE; 6H6D; 6H6H; 7N5Q |
| H2-Db | 11 | 1JPF; 6X00 |
| H2-Dd | 9 | 5KD7; 5T7G; 8D5E; 8D5F; 8FHL; 8FHU |
| H2-Dd | 10 | 1BII; 3E6H; 3ECB; 5KD4; 5WEU; 6NPR |
| H2-Kb | 8 | 1FZJ; 1G7Q; 1KBG; 1KJ3; 1KPU; 1LEG; 1LEK; 1LK2; 1NAN; 1OSZ; 1RK0; 1RK1; 1T0M; 1VAC; 2FO4; 2MHA; 2VAA; 2ZSV; 2ZSW; 3P4M; 3P4N; 3P4O; 3P9L; 3P9M; 3PAB; 3ROL; 3ROO; 4HS3; 4PV8; 4PV9; 6JQ2; 6JQ3 |
| H2-Kb | 9 | 1FZK; 1G7P; 1KPV; 1S7R; 1VAD; 1WBZ; 2VAB; 4PGB; 4PGD; 7WCY; 8P43 |
| H2-Kd | 9 | 2FWO; 4WDI; 4Z76; 5TRZ; 5TS1; 8D5J; 8D5K |

Table S7. **Structure templates to compute PWMs of peptide presentation on MHC.** The table lists the structures used to compute peptide PWMs in Fig. S17 for different MHC class I alleles in humans and mice. Structures were collected from the Protein Data Bank.

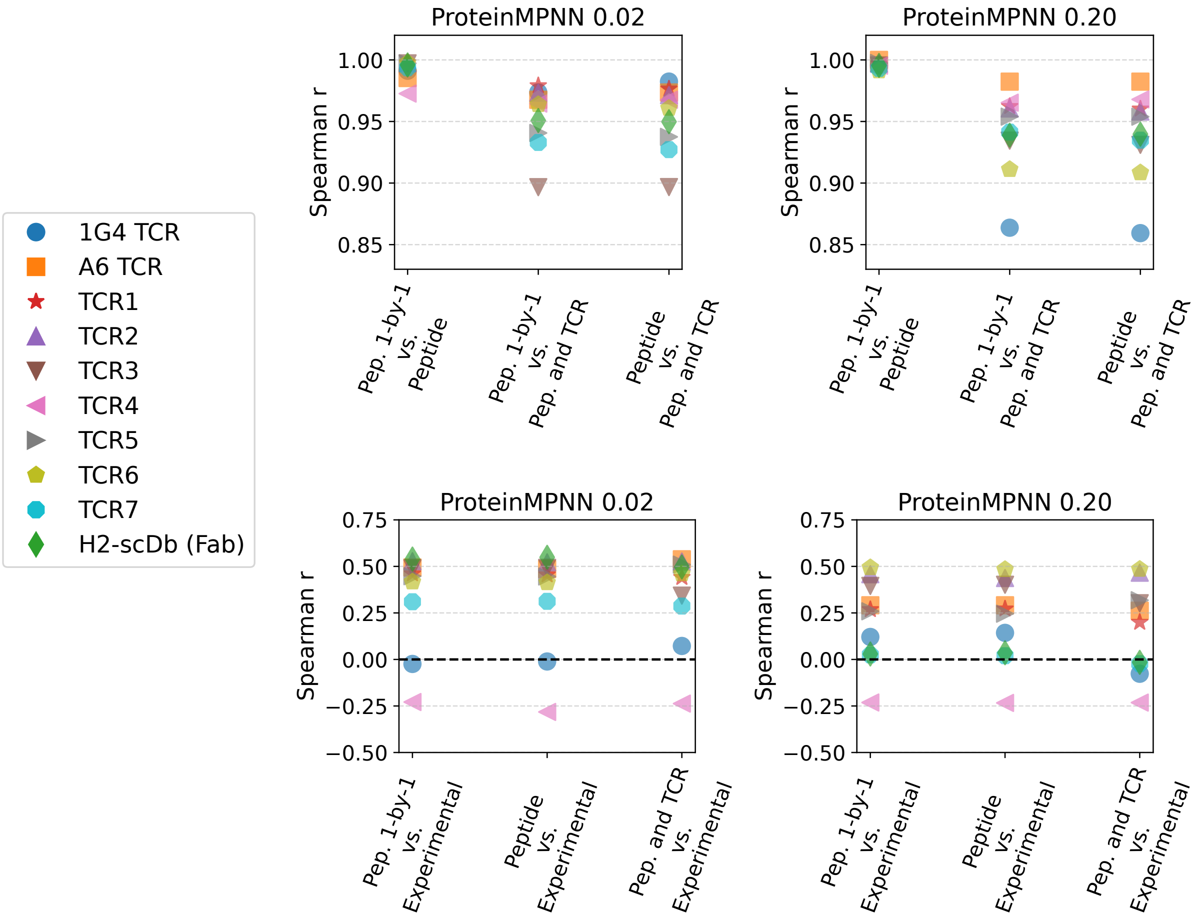


Figure S1. **Robustness of ProteinMPNN performance across scoring schemes .** The performance of ProteinMPNN across different systems (legend) is shown for models trained with varying noise amplitudes (specified above each panel), and using different scoring schemes. We consider three scoring schemes: (i): "Peptide 1-by-1" in which residues within the peptide are masked and scored one at a time, using the remainder of the TCR-pMHC complex as structural context (eq. S6), (ii) "Peptide" scheme, in which the entire peptide is masked and scored simultaneously (eq. S7), and (iii) "Pep. and TCR" scheme in which both the peptide and the nearby TCR residues are masked and scored simultaneously (eq. S8). Top panels show the Spearman correlation between predictions from different scoring schemes, while bottom panels show correlations between each scheme and experimental data. Results demonstrate that ProteinMPNN predictions are robust to the choice of scoring scheme.

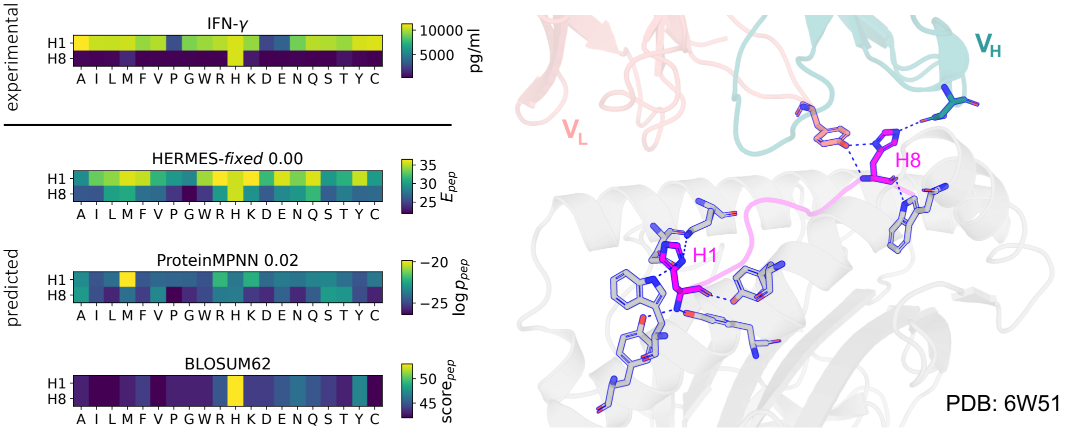


Figure S2. **Context-dependent mutational effects on T-cell activity.** The heatmaps show both experimental measurements and model predictions of T-cell activities of amino acid variants at positions 1 and 8 of the p53$^{R}157H$ peptide, which we study in complex with the designed bi-specific antibody H2-scDB and HLA-A*02:01 [21]. The wild-type peptide has a histidine at both sites. A standard substitution matrix (such as BLOSUM62) would assign identical scores to mutations at these two sites, because they share the same wild-type residue. However, peptide binding and the subsequent T-cell activity depends on the structural context of each substitution. Here, H1 forms polar contacts only with the MHC (from the sides), while H8 is more tightly constrained by polar contacts with the TCR (right panel). HERMES-*fixed* correctly leverages structural context to predict T-cell activity as a result of substitutions away from histidines at both positions. Although ProteinMPNN also incorporates structural information, its predictions are not as accurate in this instance.

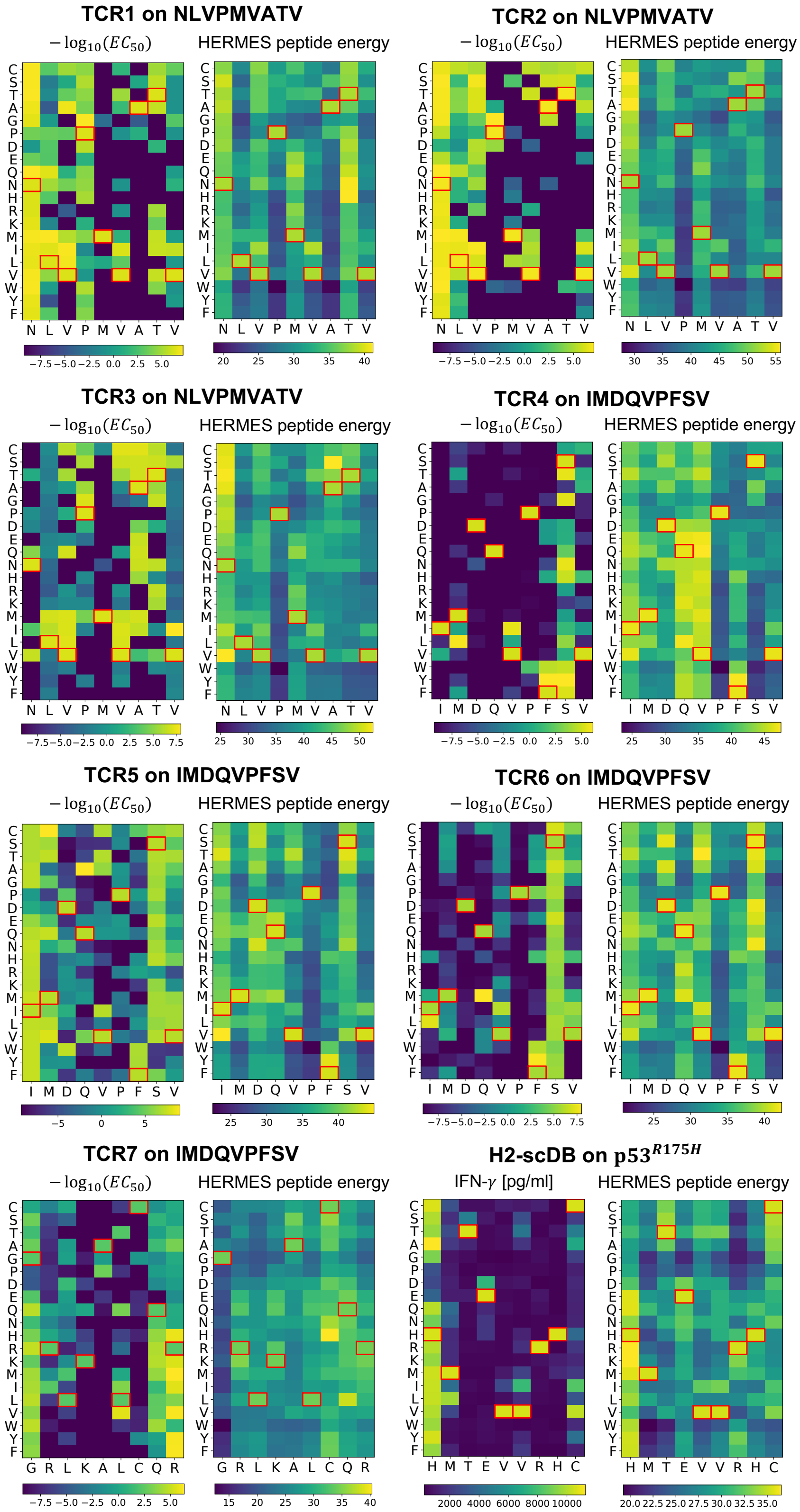


Figure S3. **Mutational effect on T-cell activities**. The heatmaps show the experimental measurements and the predictions from the HERMES-*fixed* model for the effect of all single point mutations on peptides interacting with TCR1-7 [5], and the designed bi-specific antibody Fab in the H2-scDB system [21] (similar to the data presented in scatter plots in Fig. 3). In the heatmaps, each column corresponds to a position in the peptide and is labeled with its wild-type amino acid (indicated by a red square on the heatmaps), and the rows show the amino acid substitutions.

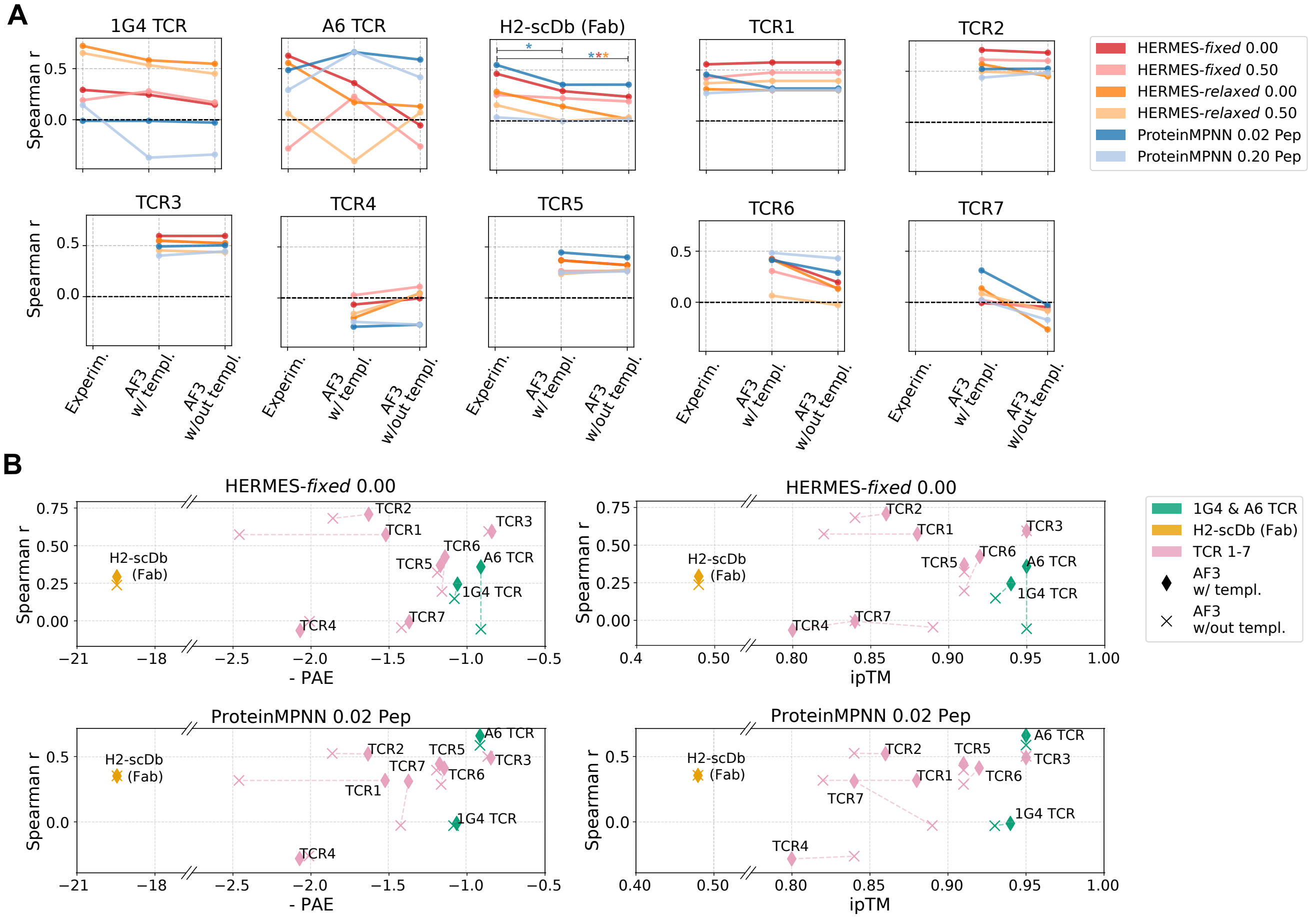


Figure S4. **HERMES and ProteinMPNN prediction accuracies for AlphaFold3-predicted structures. (A)** For all systems analyzed, we used the AlphaFold Server to model the structures of complexes with the wild-type peptides, both with and without template guidance; see Table S4 for details on the confidence of these AF3 models. For each system, the Spearman correlation ($r$) between experimental affinity or activity data and predictions from HERMES and ProteinMPNN models (colors) is shown for three structure inputs: experimental structures (when available), AF3 model with template, and AF3 model without template. Statistically significant drop in accuracy between models using experimental vs. AF3 structures–defined as cases with p-value$<$ 0.05 for difference in the Fisher's $z$-transformed Spearman correlations–are indicated with an asterisk (*) in the same color as that of the model (from the legend). Such drops are only observed for the H2-scDb Fab-pMHC system, where AF3 failed to generate a realistic structure. **(B)** Spearman correlation ($r$) between experimental affinity/activity data and predictions from the overall best HERMES-*fixed* (top) and ProteinMPNN (bottom) models from (A), plotted against AF3 confidence metrics: negative predicted alignment error ($-$PAE; left) and interchain predicted TM-score (ipTM; right). Results are shown for AF3 structures generated with and without template guidance. The confidence metrics are reported in Table S4. The HERMES model names specify the scoring scheme (fixed, relaxed), and the noise amplitude (in Å) injected to the coordinates of the structure data during training. Similarly, the ProteinMPNN model names capture scoring scheme, i.e., the whole peptide (eq. S7), and the noise amplitude injected to the training data.

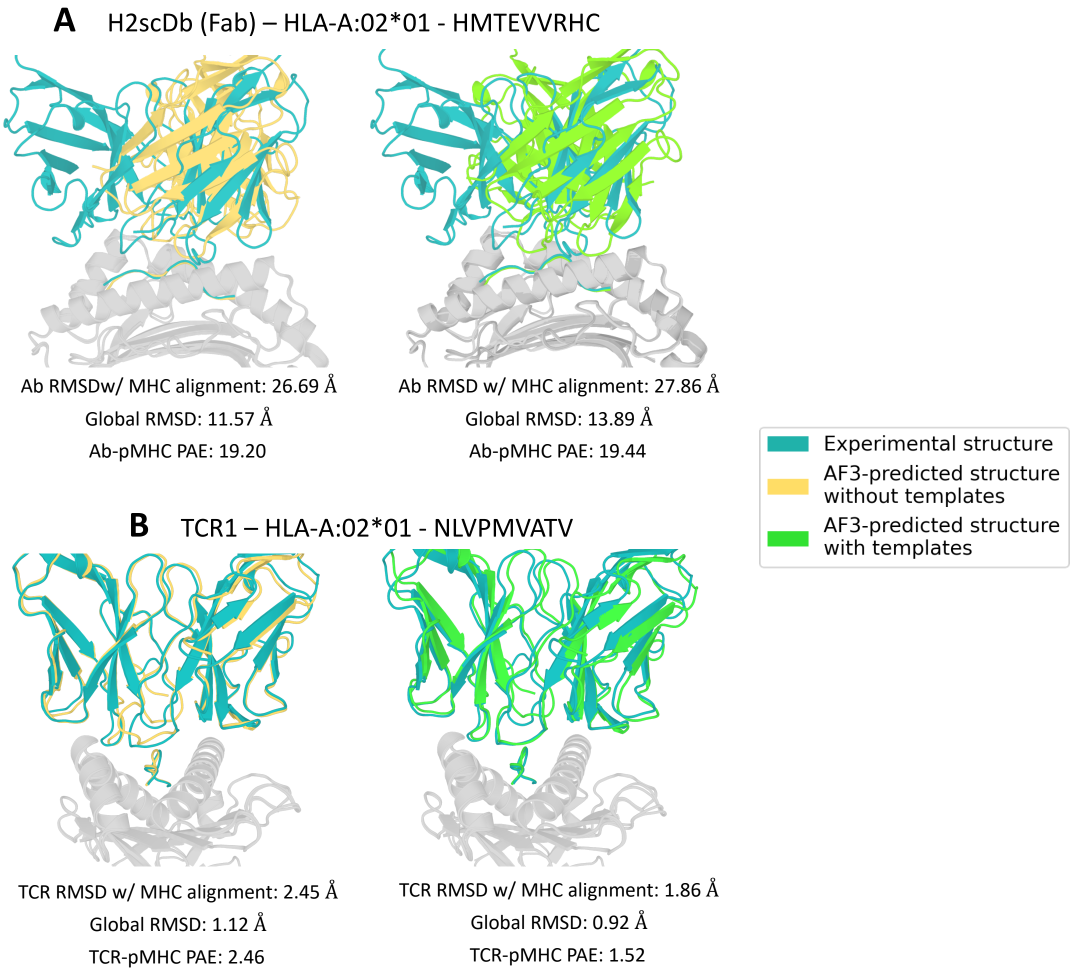


Figure S5. **Comparison of experimentally determined structures with AlphaFold3 predicted structures.** The experimentally determined structure of **(A)** H2-scDb Fab-pMHC complex, and **(B)** TCR1-pMHC complex are shown against the AF3 predicted structures generated without (left) and with (right) template guidance. All structures were aligned on the MHC using PyMOL to enable consistent visualiziation of TCR/Fab docking geometries. AF3 failed to produce a realistic structure model for the H2-scDb system, as evidenced by the high RMSD and interface PAE values reported in Table S4.

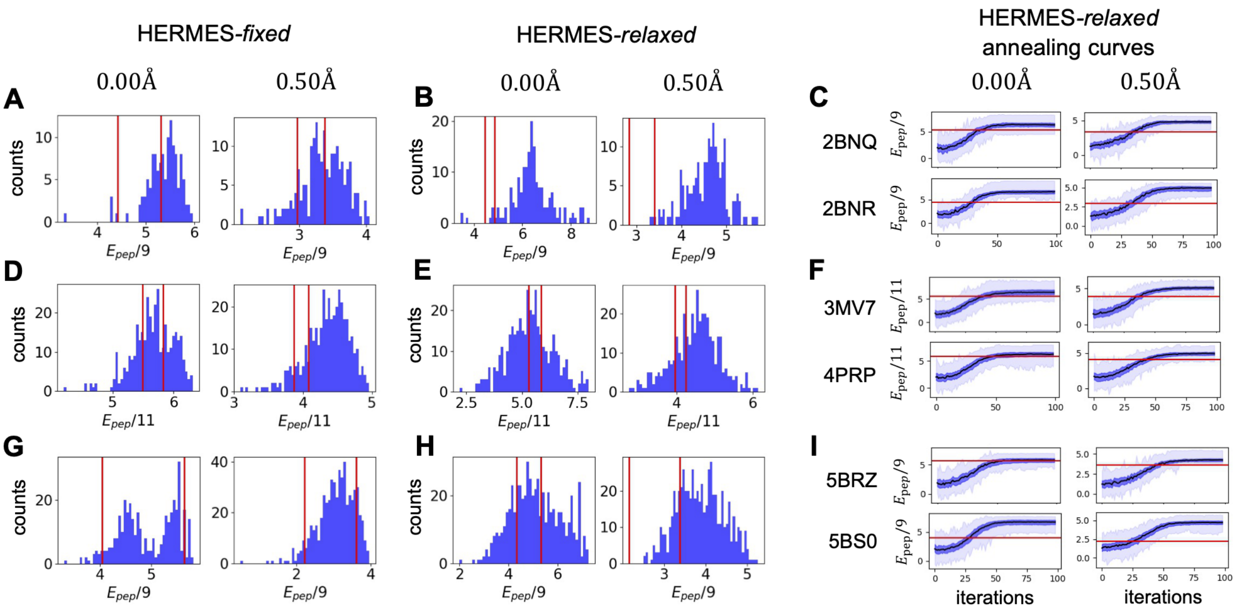


Figure S6. **Peptide design energy profiles across systems and HERMES models.** For the NY-ESO system, panels **(A)** and **(B)** show the distribution of peptide energies (eq. S1) divided by the length, for peptides designed with HERMES-*fixed* and HERMES-*relaxed* protocols, respectively. Designs were pooled across the two structure templates used (2BNQ and 2BNR), with the peptide energies computed using their respective template structures. In all panels, the red lines indicates the energies of the wild-type peptides in their corresponding structures. **(C)** The scaled energy of the designed peptides are shown as a function of the MCMC iterations during the HERMES-*relaxed* protocol, using the two different structural templates for the NY-ESO system: 2BNQ (top) and 2BNR (bottom). The red line indicates the energy of the wild-type peptide in its respective structure. In panels (A-C), designs are grouped by the noise amplitude applied during HERMES training (no noise: 0.00Å, and moderate noise: 0.50Å), as indicated above each panel. The HERMES-*relaxed* protocol consistently produces peptides with more favorable (higher) peptide energies than that of the wild-type, indicating that the protocol can successfully optimize within the peptide energy landscapes; models trained with noise are more successful in finding peptides with higher energy values. Panels **(D-F)** and **(G-I)** present analogous design statistics for the EBV and the MAGE systems, respectively.

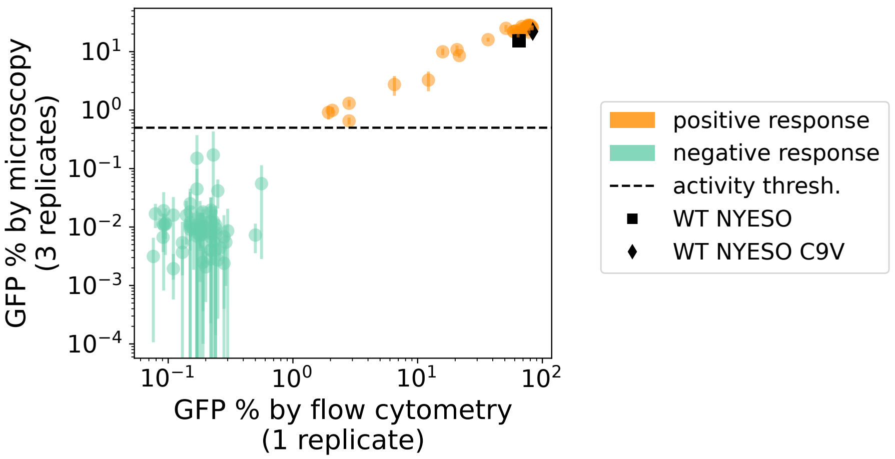


Figure S7. **Measurements of GFP levels by flow cytometry and fluorescence microscopy.** For the NY-ESO system, we performed measurements of GFP levels from peptide-induced T-cell activities, using both flow cytometry and fluorescence microscopy. Fluorescence microscopy measurements were performed in three independent replicates; markers denote the mean GFP level, and error bars show the standard deviation across the three replicates. Flow cytometry was done only on a single replicate. The two measurement methods demonstrate good agreement, with a clear separation between designs that induced significant T-cell activation (orange) and those that did not (green). The black markers indicate the activities induced by the wild-type peptides.

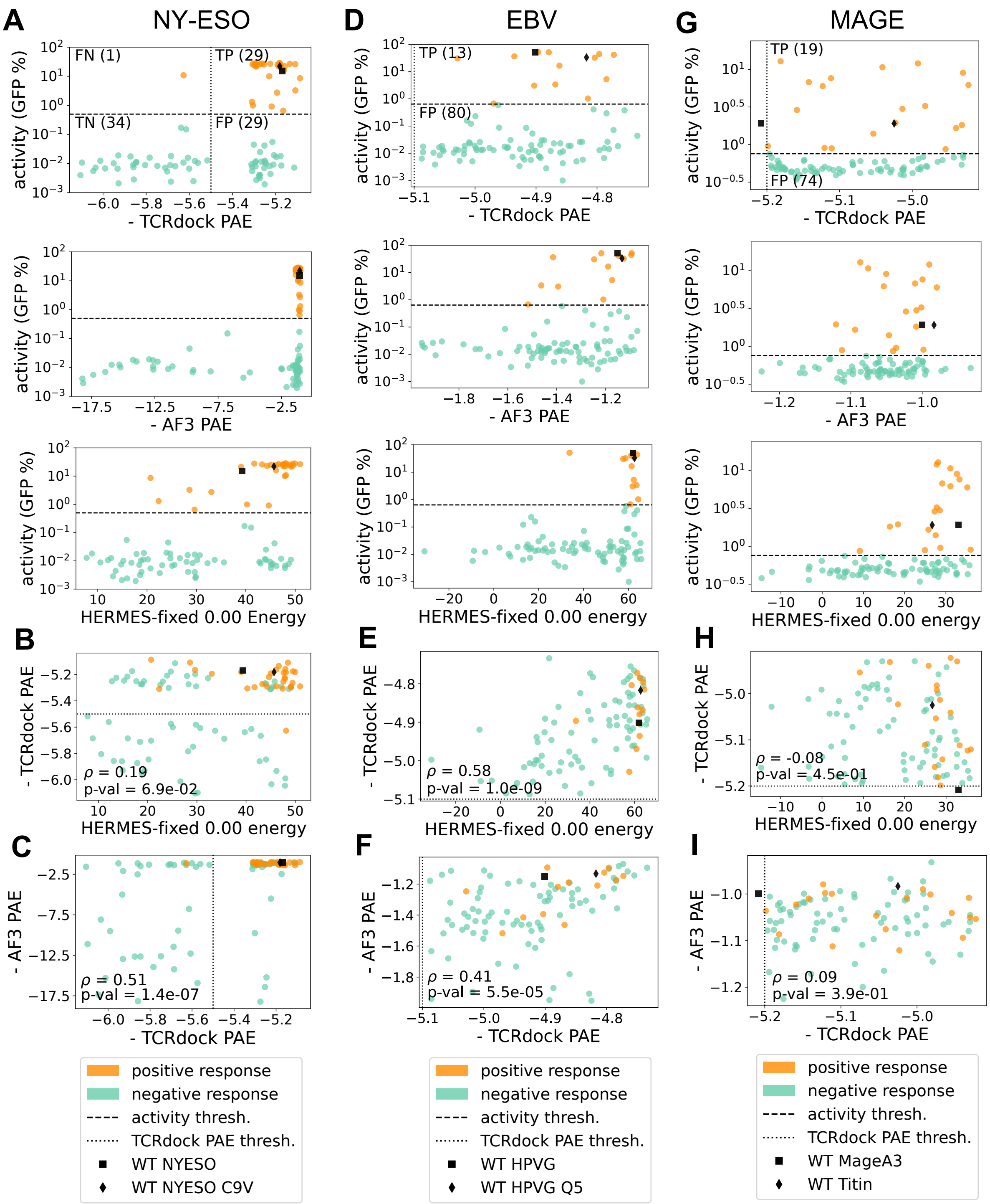


Figure S8. **TCRdock and AlphaFold3 PAEs for HERMES-designed peptides**. **(A)** For the NY-ESO system, the induced T-cell activity by the designed peptides is shown against the computed TCRdock negative-PAE (first row), the AlphaFold3 negative-PAE (second row), and the energy score predicted with HERMES-*fixed* 0.00 (non noise during training) (third row); higher negative-PAE's indicate better designs. For the designed peptides in the NY-ESO system **(B)** shows the relationship between negative TCRdock PAE values and the peptide energies from the HERMES-*fixed* 0.00. model, and **(C)** shows the negative AlphaFold3 PAE versus TCRdock PAE values. Peptides that induced significant T-cell responses are shown in orange, and non-responders are shown in green. The vertical dashed line indicate the filtering threshold for TCRdock PAE, with points to the right representing the negative design set. Panels **(D-F)** and **(G-I)** show analogous statistics for the EBV and the MAGE system, respectively. No negative designs (i.e., designs below the respective TCRdock negative-PAE thresholds) were experimentally validated for these systems. We observe that the HERMES scores and TCRdock PAE values are uncorrelated or only weakly correlated across systems, and so are TCRdock and AF3 PAE, suggesting that AF3 and TCRdock PAE's can serve as complementary filters to HERMES during design.

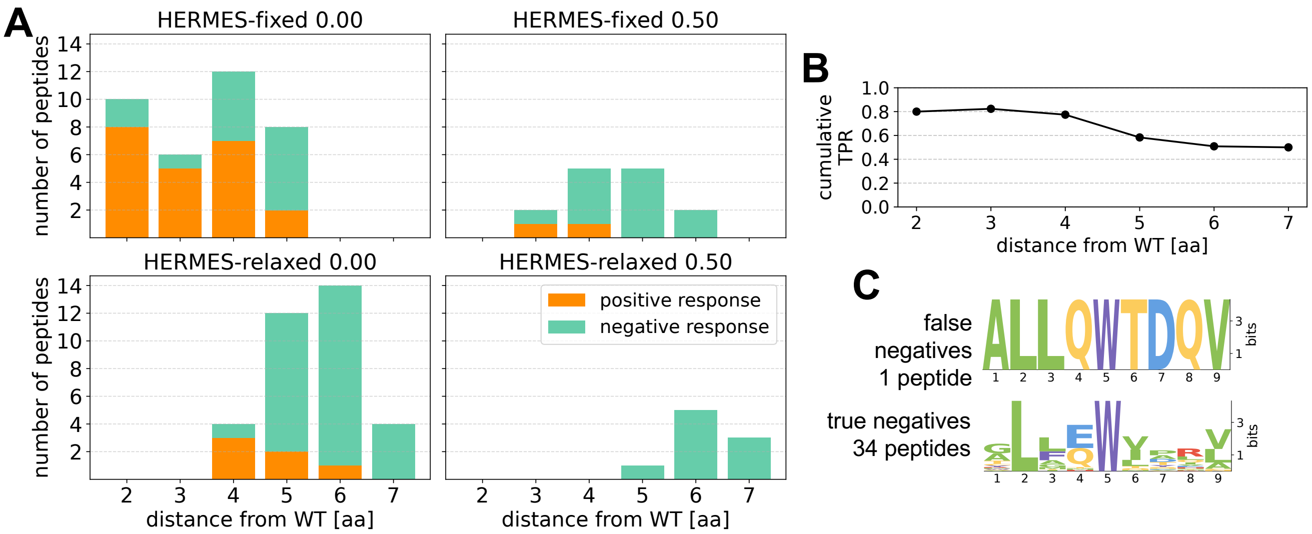


Figure S9. **Experimental T-cell activation by HERMES-designed peptides in the NY-ESO system**. **(A)** The number of peptides designed by different HERMES models (panels) that induced GFP expression beyond the T-cell activation threshold (positives; shown in orange) versus those that did not (negatives; shown in green) is shown. The designed peptides are grouped according to their minimum Hamming distance from any of the wild-type sequences used as design templates (Table S5). The HERMES model names specify the scoring scheme (fixed, relaxed), and the noise amplitude (in Å) injected to the coordinates of the structure data during training. **(B)** The cumulative True Positive Rate (TPR) for designed peptides is shown as a function of their Hamming distance from any of the wild-types. **(C)** Logo plots show the sequence composition of the False Negative and the True Negative designs.

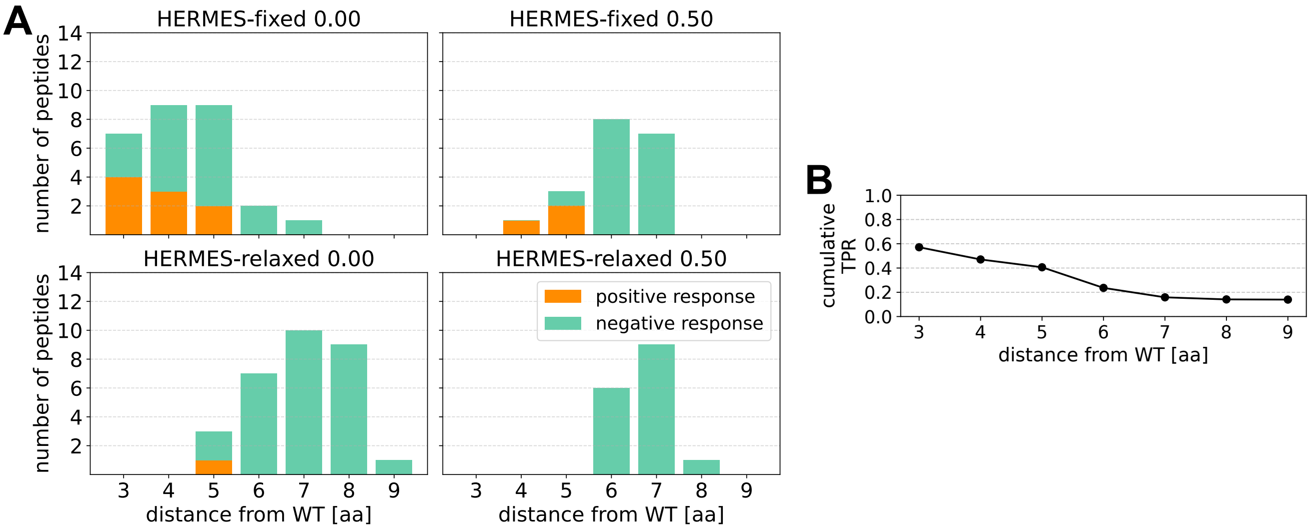


Figure S10. **Experimental T-cell activation by HERMES-designed peptides in the EBV system. (A-B)** Similar to (A-B) of Fig. S9 but for the EBV system.

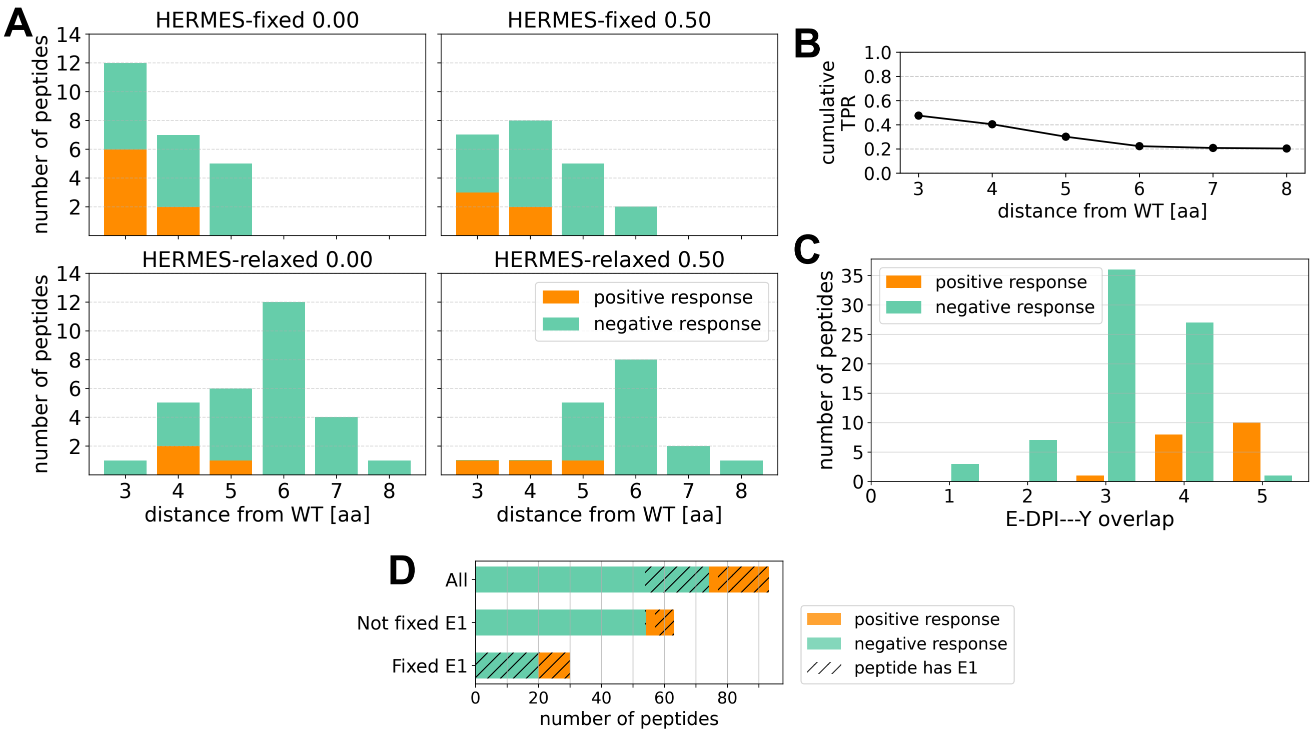


Figure S11. **Experimental T-cell activation by HERMES-designed peptides in the MAGE system. (A-B)** Similar to (A-B) of Fig. S9 but for the MAGE system. **(C)** The number of designs that induced GFP expression beyond the T-cell activation threshold (positive response; in orange) versus those that did not (negative response; in green) is shown as a function of the designs' overlap with the E-DPI- - -Y motif; this motif has been shown to enhance T-cell activation in this system [33]. The overlap is the number of amino acids that a design shares with the E-DPI- - -Y motif (maximum of 5). The fraction of positive designs increases with greater similarity to the motif. **(D)** Shown are the numbers of True Positives (orange) and False Positives (green) among designs generated under two protocols: one that enforced glutamic acid at position one ($E_1$)—labeled "fixed $E_1$"—-and one that did not enforce $E_1$—labeled "not fixed $E_1$". For each protocol, the dashed segments indicate the number of designs that contain $E_1$. The "All" bar presents the aggregate data from both protocols.

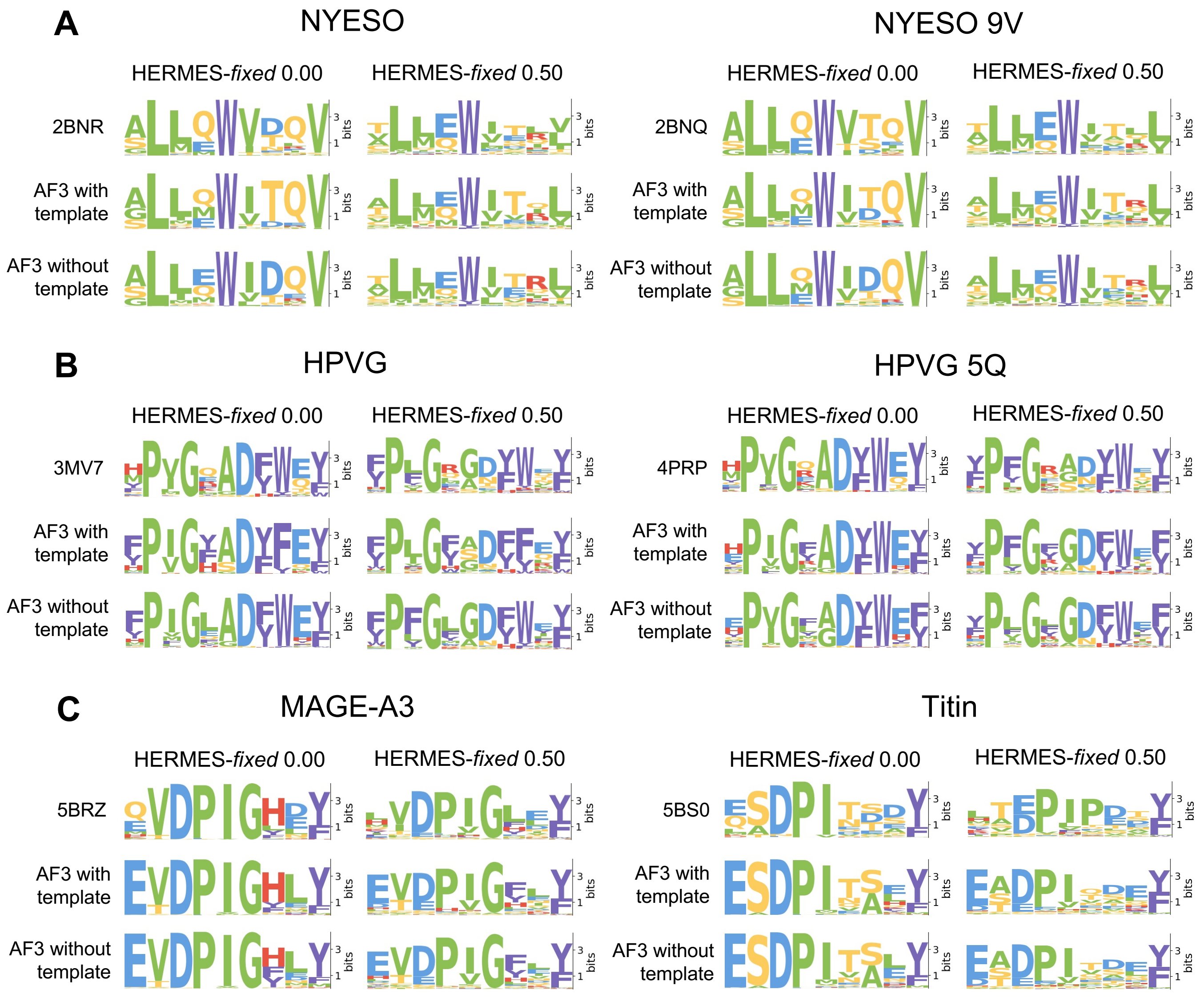


Figure S12. **Composition of designed peptides using experimental versus AlphaFold3 structures as design templates.** For the **(A)** NY-ESO, **(B)** EBV, and **(C)** MAGE, each panel displays the position weight matrices (PWMs) for peptides designed using the HERMES-*fixed* protocol. Designs were generated using three types of structural templates as indicated in each panel: the experimentally determined wild-type structure (first row), and the AlphaFold3-predicted models of that structure generated with (middle) and without (bottom) template guidance. PWMs are shown separately for the HERMES-*fixed* models trained with different noise amplitudes (as indicated above the panels). For each system, results are provided for the two wild-type peptides used as input structural templates (shown in the left and right columns of each panel).

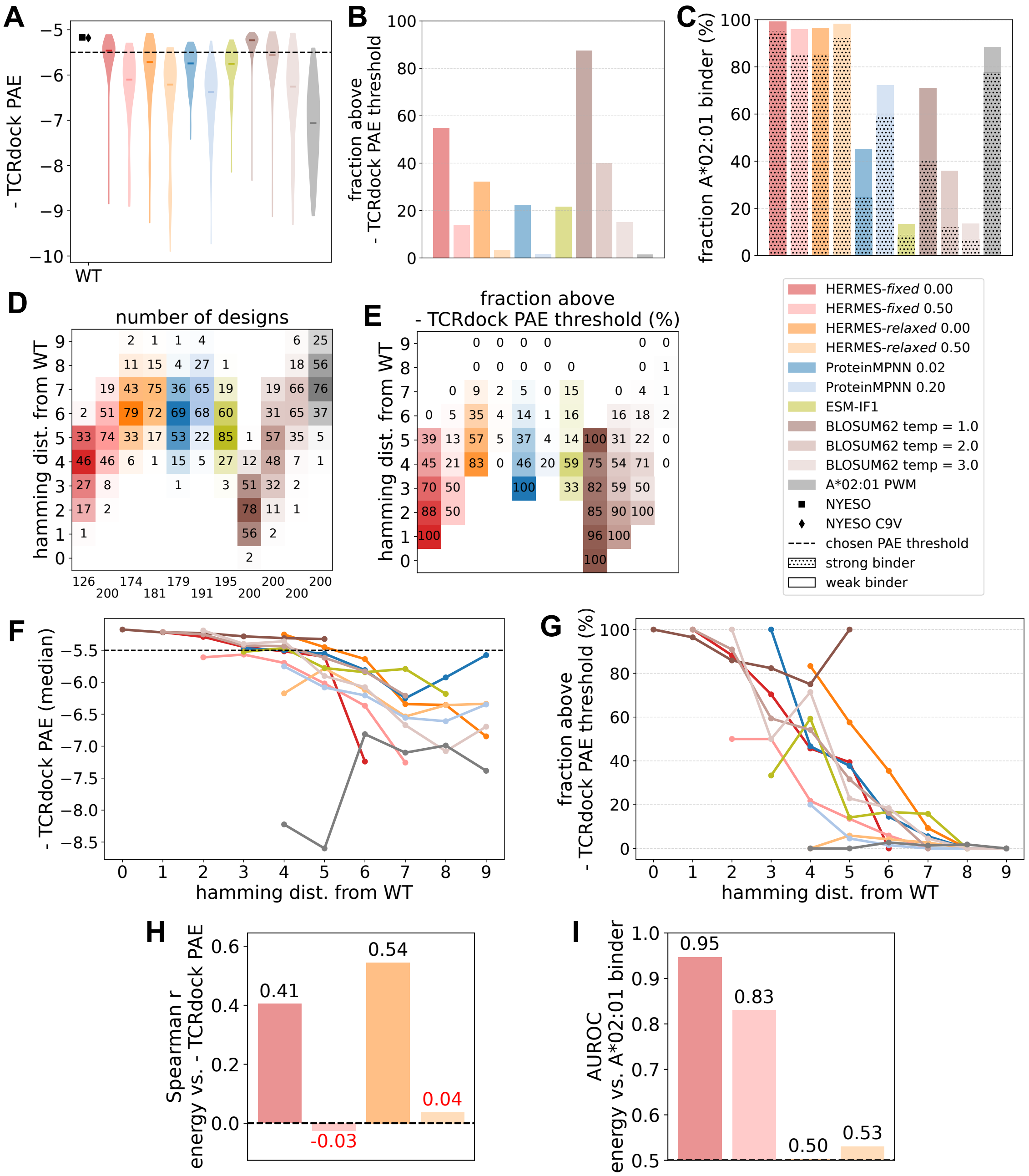

Figure S13. **Benchmarking of models for peptide design in the NY-ESO system. (A)** The distribution of the TCRdock negative-PAE's for peptides designed by different computational protocols (colors) are shown; in each case the score distribution for 100 independently designed peptides is shown. The horizontal dashed line indicates the PAE threshold chosen to classify positive designs. **(B)** The fractions of designed peptides that pass the TCRdock PAE threshold is shown for each method. **(C)** The fraction of designs that are predicted by NetMHCPan [6] to be presented by the template's MHC allele are shown; dotted regions correspond to strong binders (EL_rank $\leq 0.5$), while the bar indicates all weak and strong binders (EL_Rank $\leq 2.0$). HERMES designs are most consistently predicted to be presented by the MHC. **(D** The number of designs, and **(E)** the fraction of designs that pass the TCRdock PAE threshold are shown for each model (colors) at different Hamming distances from the wild-types; The numbers at the bottom of each column in (D) indicate the total number of unique designs. **(F)** The median of TCRdock negative-PAE across designs, and **(G)** the fraction of designs that pass the TCRdock PAE are shown for each model (colors) as a function of the Hamming distance from the wild-types. **(H)** Spearman correlation between HERMES-predicted peptide energy and TCRdock PAE is shown, with data pooled across design models. In each case, HERMES energies are computed using the same protocol applied during design. Correlations that are not statistically significant (p-value $> 0.05$) are highlighted in red. **(I)** Shows the AUROC for classifying whether a designed peptide is presented by the template's MHC allele (i.e., predicted to be a strong binder according to NetMHCPan [6]), using HERMES peptide energy as the classifier. As in (H), for each peptide, HERMES energies are computed via the same protocol used for their deign. The HERMES model names specify the design scheme (fixed, relaxed), and the noise amplitude (in Å) injected to the coordinates of the structure data during training. The ProteinMPNN design uses baseline scoring scheme with peptide 1-by-1 (eq. S6), with the training noise amplitude specified in the model name.

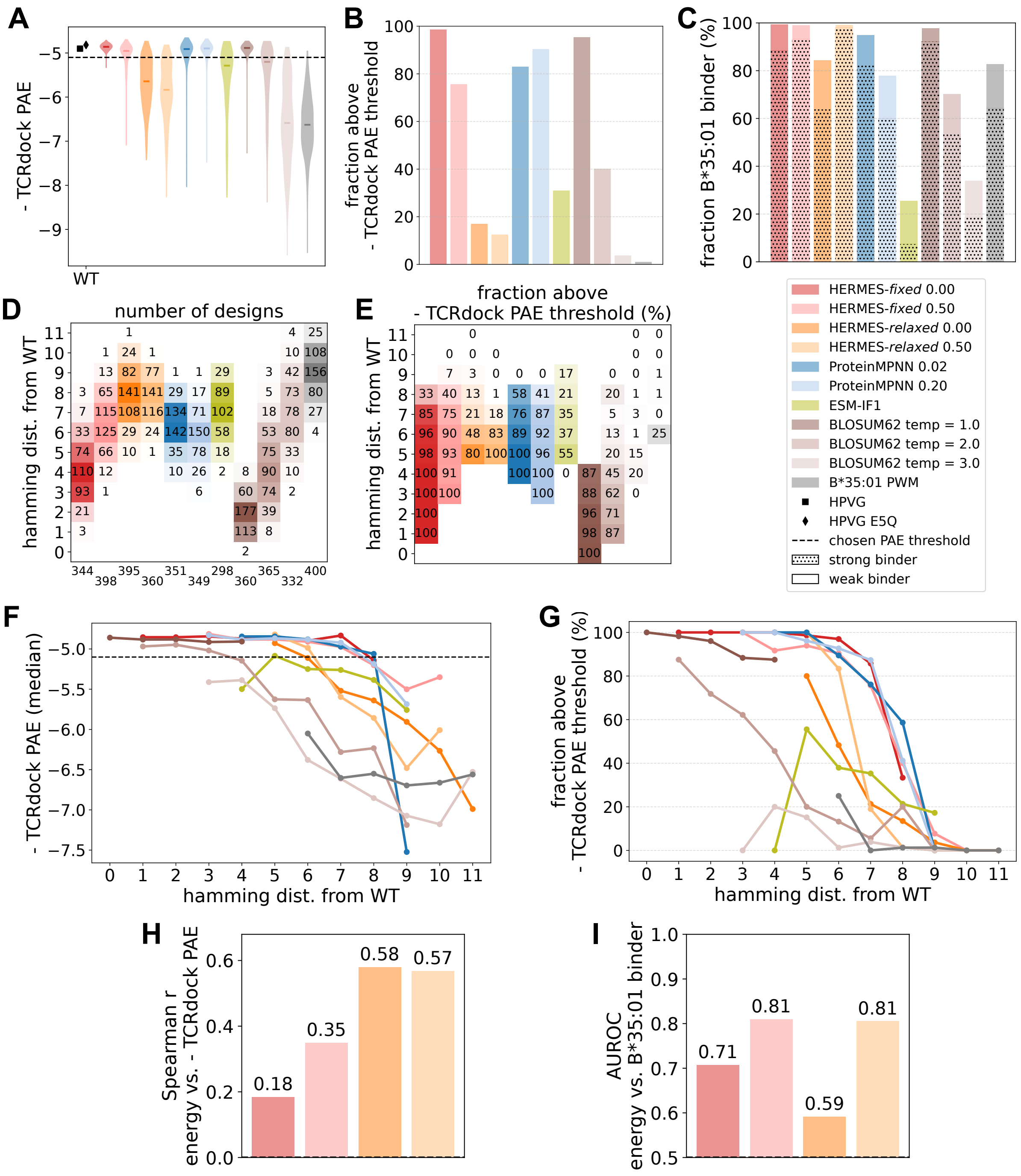


Figure S14. **Benchmarking of models for peptide design in the EBV system.** Similar to Fig. S13 but for the EBV system.

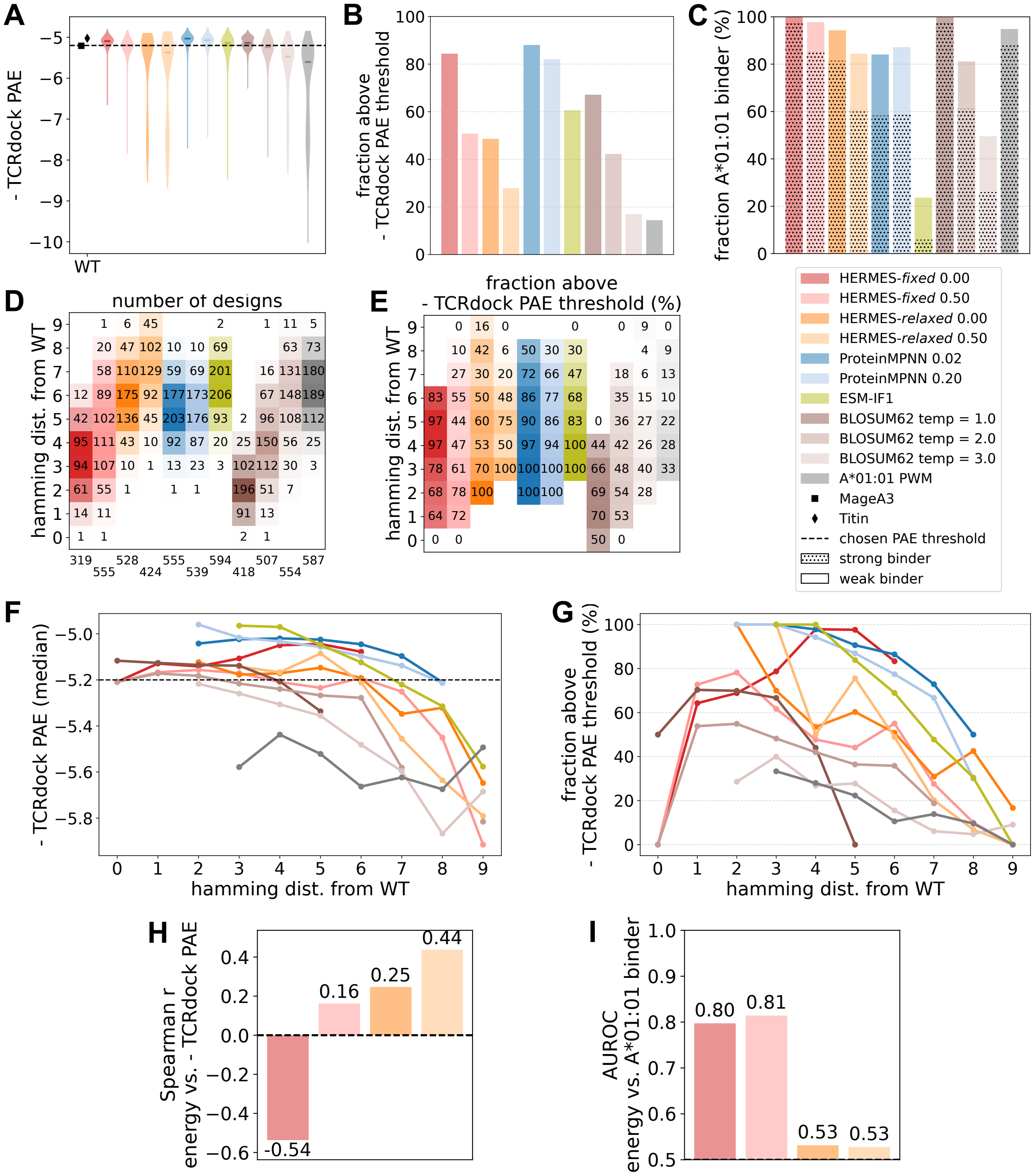


Figure S15. **Benchmarking of models for peptide design in the MAGE system.** Similar to Fig. S13 but for the MAGE system.

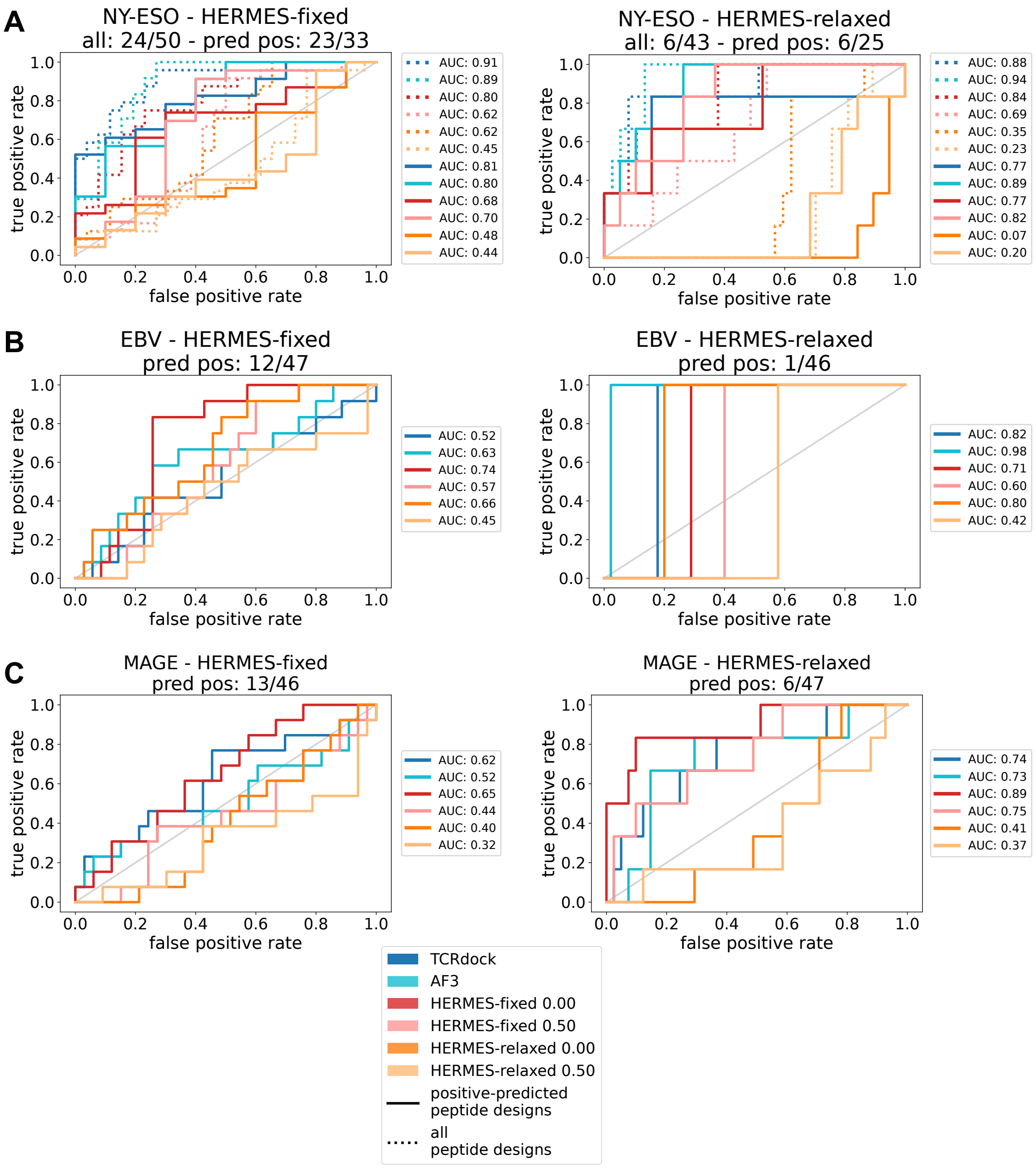


Figure S16. **Discrimination ability of different scoring schemes as design filters.** The ROC curves show true positive versus false positive rates when using scores from different models–HERMES, AlphaFold3, and TCRdock (color coded as in the legend)–as scoring criteria for peptide design. Results are shown separately for designs generated using the HERMES-*fixed* (left) and the HERMES-*relaxed* (right) protocols, for the **(A)** NY-ESO, **(B)** EBV, and **(C)** MAGE systems. Experimentally measured peptide-induced T-cell activity (GFP%) is used as the ground truth for evaluating the predictive performance in each case. For the NY-ESO system only (A) we show ROC curves computed on both peptides that were predicted to activate T-cells (positives) based on their TCRdock PAE scores (full lines), and the mixture of positive and negative peptides (dotted lines). For EBV and MAGE (B-C), the scoring criterion is only tested for the peptides that we deemed to be activating (positive) based on their TCRdock PAE scores (full lines), as these were the ones that we tested experimentally. The HERMES model names specify the design scheme (fixed, relaxed), and the noise amplitude (in Å) injected to the coordinates of the structure data during training.

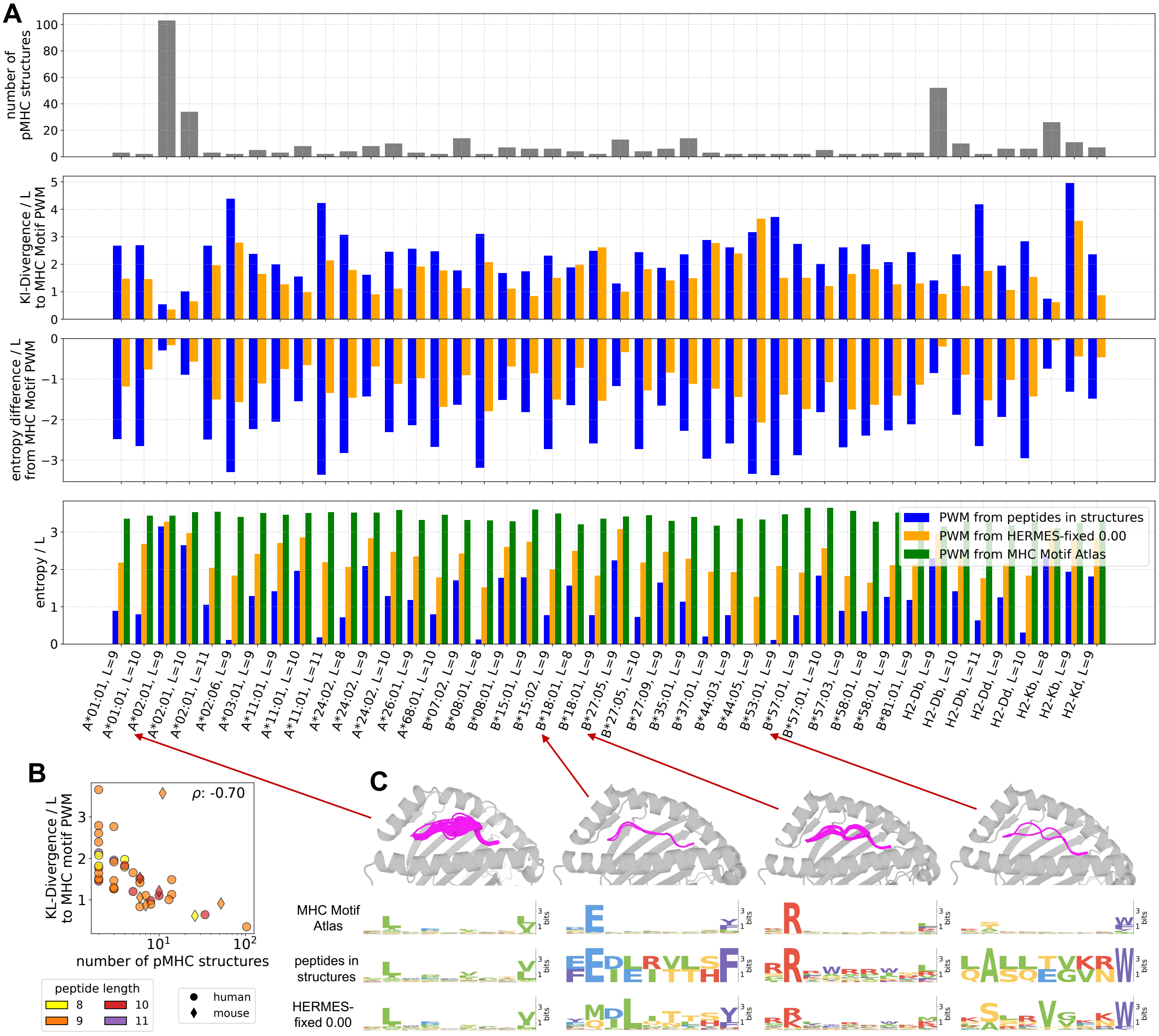


Figure S17. **Predicting MHC-I peptide presentation preferences with HERMES.** We evaluated HERMES's ability to recover MHC class I peptide presentation preferences by analyzing pMHC structures from the TCR3d database [25], grouped by MHC allele and peptide length (L), retaining only those combinations with at least two structures. As ground truth, we used position weight matrices (PWMs) from the MHC Motif Atlas [11], derived from large datasets of peptide sequences known to be presented by each allele. HERMES-*fixed* was used to design peptides for each structure in a given allele-length group, and the resulting PWMs were averaged across structures to generate a motif reflecting the peptide preferences for that MHC I allele (for a given length). As the baseline, we used PWMs generated directly from the peptides found in the structural dataset. PWM quality was evaluated using the Kullback-Leibler divergence, and the entropy difference from the ground-truth PWM. **(A)** Shown are the summaries of data and performances across all MHC I allele-length combinations: number of available structures (first row), KL-divergence between the HERMES-derived PWMs (or the baseline) to the ground truth (second row), the entropy difference to the ground truth (third row), and the entropy of the PWMs based on the three approaches (fourth row). **(B)** Shown is the KL-divergence of HERMES-derived PWMs versus the number of available pMHC structures for across different allele-length classes. **(C)** Examples of pMHC structures and the corresponding PWMs are shown for four allele-length combinations, spanning a range of structure counts. In each case, structures are aligned along the MHC chain to show the diversity of peptide poses captured by the ensemble of available structures, used as design templates.

Dataset S1. Amino-acid sequences used as input to AF3 within the AlphaFold Server to fold the TCR-pMHC and Fab-pMHC systems considered in this study.

Dataset S2. Measured T-cell activity levels and metadata for the peptide designs reported in this study. The excel file contains three sheets, one for each system considered in this study: *NYESO-designs*, *EBV-designs* and *MAGE-designs*. Three replicates of activity levels (GFP %), measured by fluorescence microscopy are reported under columns *r1*, *r2*, and *r3*; their mean values are reported under *r_mean* and the binary response label is under *resp*. For the NYESO system only, we also report the measured GFP levels through flow cytometry (one replicate). Notable metadata includes: peptide sequence, HERMES model and protocol used for the design, TCRdock PAE score, AF3 PAE score, hamming distance from the wild-types. Activity levels induced by the wild-type peptides as well as under unstimulated (negative control) condition are provided.

[1] G. M. Visani, M. N. Pun, W. Galvin, E. Daniel, K. Borisiak, U. Wagura, and A. Nourmohammad, bioRxiv **2024.07.09.602403** (2024), 10.1101/2024.07.09.602403.
[2] M. N. Pun, A. Ivanov, Q. Bellamy, Z. Montague, C. LaMont, P. Bradley, J. Otwinowski, and A. Nourmohammad, Proc. Natl. Acad. Sci. U. S. A. **121**, e2300838121 (2024), https://www.pnas.org/doi/pdf/10.1073/pnas.2300838121.
[3] M. AlQuraishi, BMC Bioinformatics **20**, 311 (2019).
[4] S. Chaudhury, S. Lyskov, and J. J. Gray, Bioinformatics **26**, 689 (2010).
[5] M. Łuksza, Z. M. Sethna, L. A. Rojas, J. Lihm, B. Bravi, Y. Elhanati, K. Soares, M. Amisaki, A. Dobrin, D. Hoyos, P. Guasp, A. Zebboudj, R. Yu, A. K. Chandra, T. Waters, Z. Odgerel, J. Leung, R. Kappagantula, A. Makohon-Moore, A. Johns, A. Gill, M. Gigoux, J. Wolchok, T. Merghoub, M. Sadelain, E. Patterson, R. Monasson, T. Mora, A. M. Walczak, S. Cocco, C. Iacobuzio-Donahue, B. D. Greenbaum, and V. P. Balachandran, Nature **606**, 389 (2022).
[6] B. Reynisson, B. Alvarez, S. Paul, B. Peters, and M. Nielsen, Nucleic Acids Research **48**, W449 (2020).
[7] P. Bradley, Elife **12**, e82813 (2023).
[8] J. Jumper, R. Evans, A. Pritzel, T. Green, M. Figurnov, O. Ronneberger, K. Tunyasuvunakool, R. Bates, A. Zídek, A. Potapenko, A. Bridgland, C. Meyer, S. A. A. Kohl, A. J. Ballard, A. Cowie, B. Romera-Paredes, S. Nikolov, R. Jain, J. Adler, T. Back, S. Petersen, D. Reiman, E. Clancy, M. Zielinski, M. Steinegger, M. Pacholska, T. Berghammer, S. Bodenstein, D. Silver, O. Vinyals, A. W. Senior, K. Kavukcuoglu, P. Kohli, and D. Hassabis, Nature **596**, 583 (2021).
[9] B. Liu, N. F. Greenwood, J. E. Bonzanini, A. Motmaen, J. Sharp, C. Wang, G. M. Visani, D. K. Vafeados, N. Roullier, A. Nourmohammad, K. C. Garcia, and D. Baker, bioRxiv **2024.11.28.625793** (2024), 10.1101/2024.11.28.625793.
[10] S. Henikoff and J. G. Henikoff, Proc. Natl. Acad. Sci. U. S. A. **89**, 10915 (1992).
[11] D. M. Tadros, S. Eggenschwiler, J. Racle, and D. Gfeller, Nucleic Acids Res. **51**, D428 (2023).
[12] C. Lundegaard, K. Lamberth, M. Harndahl, S. Buus, O. Lund, and M. Nielsen, Nucleic Acids Res. **36**, W509 (2008).
[13] C. Lundegaard, O. Lund, and M. Nielsen, Bioinformatics **24**, 1397 (2008).
[14] M. Andreatta and M. Nielsen, Bioinformatics **32**, 511 (2016).
[15] J. Dauparas, I. Anishchenko, N. Bennett, H. Bai, R. J. Ragotte, L. F. Milles, B. I. M. Wicky, A. Courbet, R. J. de Haas, N. Bethel, P. J. Y. Leung, T. F. Huddy, S. Pellock, D. Tischer, F. Chan, B. Koepnick, H. Nguyen, A. Kang, B. Sankaran, A. K. Bera, N. P. King, and D. Baker, Science **378**, 49 (2022).
[16] C. Hsu, R. Verkuil, J. Liu, Z. Lin, B. Hie, T. Sercu, A. Lerer, and A. Rives, in *Proceedings of the 39th International Conference on Machine Learning*, Proceedings of Machine Learning Research, Vol. 162, edited by K. Chaudhuri, S. Jegelka, L. Song, C. Szepesvari, G. Niu, and S. Sabato (PMLR, 2022) pp. 8946–8970.
[17] B. Meynard-Piganeau, C. Feinauer, M. Weigt, A. M. Walczak, and T. Mora, Proc. Natl. Acad. Sci. U. S. A. **121**, e2316401121 (2024).
[18] M. F. Jensen and M. Nielsen, eLife **12**, RP93934 (2024).
[19] E. Fast, M. Dhar, and B. Chen, bioRxiv , 2023.09.12.557285 (2023).
[20] J. Abramson, J. Adler, J. Dunger, R. Evans, T. Green, A. Pritzel, O. Ronneberger, L. Willmore, A. J. Ballard, J. Bambrick, S. W. Bodenstein, D. A. Evans, C.-C. Hung, M. O'Neill, D. Reiman, K. Tunyasuvunakool, Z. Wu, A. Zemgulytė, E. Arvaniti, C. Beattie, O. Bertolli, A. Bridgland, A. Cherepanov, M. Congreve, A. I. Cowen-Rivers, A. Cowie, M. Figurnov, F. B. Fuchs, H. Gladman, R. Jain, Y. A. Khan, C. M. R. Low, K. Perlin, A. Potapenko, P. Savy, S. Singh, A. Stecula, A. Thillaisundaram, C. Tong, S. Yakneen, E. D. Zhong, M. Zielinski, A. Zídek, V. Bapst, P. Kohli, M. Jaderberg, D. Hassabis, and J. M. Jumper, Nature **630**, 493 (2024).
[21] E. H.-C. Hsiue, K. M. Wright, J. Douglass, M. S. Hwang, B. J. Mog, A. H. Pearlman, S. Paul, S. R. DiNapoli, M. F. Konig, Q. Wang, A. Schaefer, M. S. Miller, A. D. Skora, P. A. Azurmendi, M. B. Murphy, Q. Liu, E. Watson, Y. Li, D. M. Pardoll, C. Bettegowda, N. Papadopoulos, K. W. Kinzler, B. Vogelstein, S. B. Gabelli, and S. Zhou, Science **371**, eabc8697 (2021).
[22] J. Robinson, M. J. Waller, P. Parham, N. de Groot, R. Bontrop, L. J. Kennedy, P. Stoehr, and S. G. E. Marsh, Nucleic Acids Res. **31**, 311 (2003).
[23] J. M. Heather, M. J. Spindler, M. H. Alonso, Y. I. Shui, D. G. Millar, D. S. Johnson, M. Cobbold, and A. N. Hata, Nucleic Acids Res. **50**, e68 (2022).
[24] J. P. Roney and S. Ovchinnikov, Phys. Rev. Lett. **129**, 238101 (2022).
[25] R. Gowthaman and B. G. Pierce, Bioinformatics **35**, 5323 (2019).
[26] J.-L. Chen, G. Stewart-Jones, G. Bossi, N. M. Lissin, L. Wooldridge, E. M. L. Choi, G. Held, P. R. Dunbar, R. M. Esnouf, M. Sami, J. M. Boulter, P. Rizkallah, C. Renner, A. Sewell, P. A. van der Merwe, B. K. Jakobsen, G. Griffiths, E. Y. Jones, and V. Cerundolo, Journal of Experimental Medicine **201**, 1243 (2005).
[27] D. N. Garboczi, P. Ghosh, U. Utz, Q. R. Fan, W. E. Biddison, and D. C. Wiley, Nature **384**, 134 (1996).
[28] Y.-H. Ding, B. M. Baker, D. N. Garboczi, W. E. Biddison, and D. C. Wiley, Immunity **11**, 45 (1999).
[29] X. Yang, M. Gao, G. Chen, B. G. Pierce, J. Lu, N.-P. Weng, and R. A. Mariuzza, J. Biol. Chem. **290**, 29106 (2015).
[30] S. Gras, Z. Chen, J. J. Miles, Y. C. Liu, M. J. Bell, L. C. Sullivan, L. Kjer-Nielsen, R. M. Brennan, J. M. Burrows, M. A. Neller, R. Khanna, A. W. Purcell, A. G. Brooks, J. McCluskey, J. Rossjohn, and S. R. Burrows, Journal of Experimental Medicine **207**, 1555 (2010).
[31] Y. C. Liu, Z. Chen, M. A. Neller, J. J. Miles, A. W. Purcell, J. McCluskey, S. R. Burrows, J. Rossjohn, and S. Gras, J. Biol. Chem. **289**, 16688 (2014).
[32] M. C. C. Raman, P. J. Rizkallah, R. Simmons, Z. Donnellan, J. Dukes, G. Bossi, G. S. Le Provost, P. Todorov, E. Baston,

E. Hickman, T. Mahon, N. Hassan, A. Vuidepot, M. Sami, D. K. Cole, and B. K. Jakobsen, Sci. Rep. **6**, 18851 (2016).

[33] B. J. Cameron, A. B. Gerry, J. Dukes, J. V. Harper, V. Kannan, F. C. Bianchi, F. Grand, J. E. Brewer, M. Gupta, G. Plesa, G. Bossi, A. Vuidepot, A. S. Powlesland, A. Legg, K. J. Adams, A. D. Bennett, N. J. Pumphrey, D. D. Williams, G. Binder-Scholl, I. Kulikovskaya, B. L. Levine, J. L. Riley, A. Varela-Rohena, E. A. Stadtmauer, A. P. Rapoport, G. P. Linette, C. H. June, N. J. Hassan, M. Kalos, and B. K. Jakobsen, Sci. Transl. Med. **5**, 197ra103 (2013).
